# Exon Junction Complex dependent mRNA localization is linked to centrosome organization during ciliogenesis

**DOI:** 10.1101/2020.10.28.358960

**Authors:** Oh Sung Kwon, Rahul Mishra, Adham Safieddine, Emeline Coleno, Quentin Alasseur, Marion Faucourt, Isabelle Barbosa, Edouard Bertrand, Nathalie Spassky, Hervé Le Hir

## Abstract

Exon junction complexes (EJC) mark untranslated spliced mRNAs and are crucial for the mRNA lifecycle. An imbalance in EJC dosage alters mouse neural stem cell (mNSC) division and is linked to human neurodevelopmental disorders. In quiescent mNSC and immortalized human retinal pigment epithelial (RPE1) cells, centrioles form a basal body for ciliogenesis. Here, we report that EJCs accumulate at basal bodies of mNSC or RPE1 cells and decline when these cells differentiate or resume growth. A high-throughput smFISH screen identifies two transcripts accumulating at centrosomes in quiescent cells, *NIN* and *BICD2*. In contrast to *BICD2*, the localization of *NIN* transcripts is EJC-dependent. *NIN* mRNA encodes a core component of centrosomes required for microtubule nucleation and anchoring. We find that EJC down-regulation impairs both pericentriolar material organization and ciliogenesis. An EJC-dependent mRNA trafficking towards centrosome and basal bodies might contribute to proper mNSC division and brain development.

## Introduction

Messenger RNAs result from a succession of maturation steps that modify transcript extremities and excise introns. These processes are tightly coupled to the transcription machinery, and ultimately lead to the packaging of mature mRNAs into large ribonucleoparticles composed of numerous RNA-binding proteins (RBPs)^1^. Each messenger ribonucleoprotein (mRNP) particle is composed of ubiquitous RBPs including cap-binding proteins, exon junction complexes (EJC) and polyA-binding proteins, as well as hundreds of additional common and cell-specific RBPs^2–6^. These RBPs densely pack the mRNP particles^7,8^ and govern the fate and the functions of mRNAs^1^.

EJCs are deposited upstream exon-exon junctions by the splicing machinery and are potentially present in multiple copies along transcripts^9,10^. The EJC core complex is composed of four proteins: the RNA helicase eIF4A3 (eukaryotic initiation factor 4A3 or DDX48), the heterodimer MAGOH/Y14 (or RBM8) and MLN51 (Metastatic Lymph Node 51 or CASC3)^11,12^. At the center, eIF4A3 clamps RNA to ensure an unusually stable binding^13,14^ A dozen of additional factors bind directly or indirectly the EJC core and constitute EJC peripheral factors^15^. mRNP particles are largely remodeled upon translation in the cytoplasm^7,8^ and EJCs are disassembled at this step by scanning ribosomes^16^. Therefore, EJCs mark a precise period in the mRNA lifecycle between nuclear splicing and cytosolic translation. During this period, EJCs contribute to splicing regulation and to the recruitment of nuclear export factors^15,17^ In the cytoplasm, EJCs are intimately linked to mRNA translation and stability. First, EJCs enhance the translational efficiency of newly made mRNP by communicating with the translation machinery^15,18,19^. Second, EJCs serve as a signal for nonsense-mediated mRNA decay (NMD), when translation termination occurs before the last exon-exon junction. Thus, NMD couples the translation and degradation machineries to eliminate transcripts encoding truncated proteins or to regulate the stability of specific transcript in a translation-dependent manner^20^.

The implication of EJCs in several crucial steps of gene expression explains why its components are essential for cellular viability^21^. In several organisms, a precise dosage of EJC components is required for proper development^22–28^. In humans, mutations leading to hypomorphic expression of Y14 and eIF4A3 are associated to two distinct syndromes with common neurodevelopmental phenotypes^29^. The thrombocytopenia with absent radius (TAR) syndrome is associated with a reduction of Y14 expression and it presents some defects in limb development and platelet production^30^. In the case of eIF4A3, it is linked to the autosomal recessive Richieri-Costa-Pereira syndrome (RCPS) presenting both limb and craniofacial dysmorphisms^31^. Copy number variants of EJC and NMD factors were also found in patients with intellectual disabilities^32^. A major step in understanding the link between EJC dosage affection and brain development and function in mammals derived from mouse genetics. A pioneer mutagenesis screen unraveled that MAGOH haploinsufficiency results in smaller body size and microcephaly by regulating division of Neural Stem Cells (NSC)^33^. A conditional *Magoh* allelic knock-out leading to NSC-specific reduction in MAGOH expression confirmed its importance for cortical development. In these cells, NSC mitosis is delayed, leading to a decrease of intermediary progenitors (IP), a premature generation of neurons and an increased apoptosis of their progeny^33–35^. Remarkably, the generation of *Rbm8a* (encoding Y14) as well as *eIF4A3* conditional haplo-insufficiency in mNSC phenocopied the effects observed with *Magoh* on embryonic neurogenesis, with a notable microcephaly^36,37^ However, a *Mln51* conditional haploinsufficiency only partially phenocopied the three other EJC core components with less profound neurodevelopmental disorders, suggesting a more tissue-specific involvement of MLN51^38^. EJC-associated NMD factors have also been associated to NSC maintenance and differentiation^39–41^. A proper dosage of fully assembled EJCs, and not only its free components, is thus clearly essential for NSC division, differentiation and brain development. However, the precise mechanisms at play remain elusive.

These observations prompted us to study EJC core proteins in primary cultures of radial glial mNSC, which are quiescent monociliated cells. Centrosomes are composed of a pair of centrioles and a matrix of pericentriolar material (PCM) that nucleates microtubules and participates in cell cycle and signaling regulation^42^. When cells exit the cell cycle, the centriole pair migrates to the cell surface, and the mother centriole constitutes a basal body for primary cilium formation^42^. We observed that EJC core proteins concentrate around centrosomes at the base of primary cilia both in mNSCs and human retinal pigment epithelial (RPE1) cells. This centrosomal accumulation of EJC proteins is predominant during the quiescent state as it diminishes upon cell differentiation or cell-cycle re-entry. The accumulation of EJC complexes around centrosomes is RNA-dependent and ensured by a microtubule-dependent pathway. A single molecule FISH (smFISH) screen identifies two mRNAs, *NIN* and *BICD2* localizing at centrosome in quiescent RPE1 cells. Remarkably, both EJC and translation are essential for *NIN* mRNA localization. Down-regulation of EJC impaired ciliogenesis and organization of the PCM, establishing a potential link between the molecular and physiological functions of the EJC.

## Results

### EIF4A3 and Y14 label centrosomes in quiescent mNSC

Reduced expression of any of the EJC core components in mice induces defects in NSC division and differentiation^29^. This prompted us to study the expression of EJC core proteins in mNSCs. We first investigated primary cultures of glial progenitors isolated from newborn mice forebrain^43^. Upon serum starvation, quiescent mono-ciliated radial glial cells differentiate into ependymal cells^44^. Ependymal cells are multi-ciliated and are present at the surface of brain ventricles. Beating of their cilia contributes to the flow of cerebrospinal fluid. In radial glial cells, the primary cilium grows from the basal body docked at the membrane. During differentiation, amplification of centrioles leads to the production of multiple cilia at the surface of ependymal cells^45^.

Antibodies against FGFR1 Oncogene Partner (FOP) label the distal end of centrioles of mono and multiciliated cells and the pericentriolar area^46,47^, whereas antibodies against polyglutamylated tubulin decorate both centrioles and cilia^48^. Both antibodies clearly distinguished the mono- (Fig. 1a, c) and multi-ciliated (Fig. 1b, d) states of mNSCs and ependymal cells, respectively. We investigated the localization of the EJC core components eIF4A3 and Y14. As previously observed in other cells^49–51^, eIF4A3 and Y14 were mainly nuclear in both mono- and multi-ciliated mNSCs (Fig. 1a-d). However, we noticed that both eIF4A3 and Y14 concentrate around the centrosome at the base of primary cilia in the majority of quiescent mNSCs (Fig. 1a, c, e-h and Supplementary Fig. 1a, b). In contrast, ependymal cells do not show a strong eIF4A3 and Y14 staining around centrioles (Fig. 1b, d, e-h and Supplementary Fig. 1c-d). The reduced concentration of both proteins around centrioles in ependymal cells was not due to an overall lower expression of the two proteins as the nuclear signals of eIF4A3 and Y14 increased by 1.5 fold in ependymal cells compared to quiescent mNSCs (Supplementary Fig. 1e, f).

**Figure 1.**
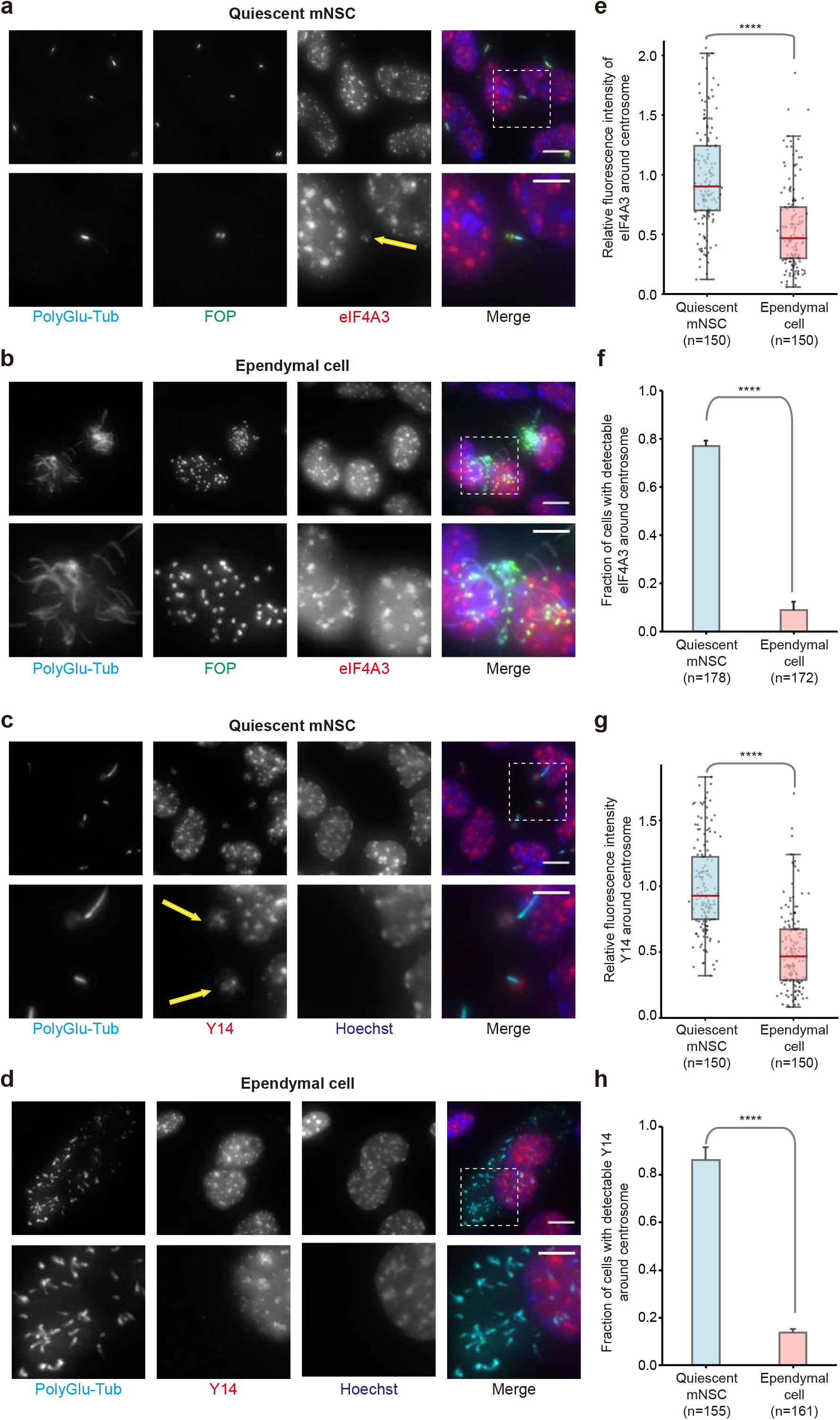
EJC core components, eIF4A3 and Y14, strongly localize around centrosomes in quiescent mNSC, but not in differentiated ependymal cells. Quiescent mNSC (a,c) and multiciliated ependymal cells (b, d) were stained for eIF4A3 (a, b) or Y14 (c, d). Centrosomes were labeled by FOP antibody. Primary cilia and centriole were stained by poly-glutamylated tubulin antibody. Nuclei were stained by Hoechst. Images result from maximum intensity projections of 12 z-stacks acquired at every 0.5 μm. Lower panels show enlarged images that are marked by white dashed square in the upper panel. Scale bars in the upper and lower panels are 5 μm and 3 μm, respectively. Fluorescence intensities for eIF4A3 (e) and for Y14 (h) were quantified in 2 μm circles around centrosomes and plotted as fluorescence intensities relative to the average fluorescence intensity in quiescent mNSC (set as 1.0). The red lines mark the median values and values between the 25th lower percentile and 75th higher percentile are in the box. Whiskers above and below the box correspond 0.35th lower percentile and 99.65th higher percentile, respectively. The fraction of cells with detectable centrosomal eIF4A3 (f) or Y14 (h) was determined in either quiescent mNSC or ependymal cells. Three independent experiments were performed. The number of cells analyzed for each independent experiment is provided (e-h). Error bars correspond to S.D. **** P ≤ 0.0001, Mann-Whitney test (e, g) and two-tailed t-test (f, h).

Together, these data showed that at least two EJC core proteins accumulate in the vicinity of centrioles in monociliated mNSCs and this cytoplasmic localization decreases upon differentiation into ependymal cells.

### EJC core components accumulate around centrosomes in ciliated quiescent RPE1 cells

To test the generality of this observation, we investigated the localization of EJC core proteins in the telomerase-immortalized Retinal Pigment Epithelial cell line hTERT-RPE1 (RPE1 cells), a popular cellular model to study primary cilia^52,53^. After 2 days of serum starvation, around 80% of RPE1 cells possessed a primary cilium, compared to only 9% in proliferating cells cultivated with serum (Fig. 2a-c). Y14 and MAGOH form a stable and obligated heterodimer^54,55^. Given that no antibodies against MAGOH provided specific immunofluorescence signals, we did not analyze MAGOH localization. And, as previously observed in other cells^51,56^, MLN51 is mainly detected in the cytoplasmic compartment of RPE1 cells (Supplementary Fig. 2e, f). It generated a background preventing the detection of its potential enrichment around centrosomes. As expected, eIF4A3 and Y14 were mainly localized in the nuclear compartment where they concentrated in nuclear speckles, corresponding to punctuate domains enriched in splicing factors and labeled by SC35 and/or 9G8 antibodies (Fig. 2a, b, Supplementary Fig. 1g-j and Supplementary Fig. 2a, b). Remarkably, in a large fraction of quiescent RPE1 cells, eIF4A3 and Y14 also concentrated around centrioles (Fig. 2a,d and Supplementary Fig. 2a, c). In contrast, eIF4A3 and Y14 were not accumulating around the centrosome of proliferating cells (Fig. 2b and Supplementary Fig. 2b). The relative fluorescence intensity of eIF4A3 and Y14 around centrosome was 1.5 times higher in quiescent cells than in proliferating cells (Fig. 2e and Supplementary Fig. 2d). In quiescent RPE1 cells, eIF4A3 and Y14 both accumulate around centrosomes at the base of primary cilia like in quiescent NSC.

**Figure 2.**
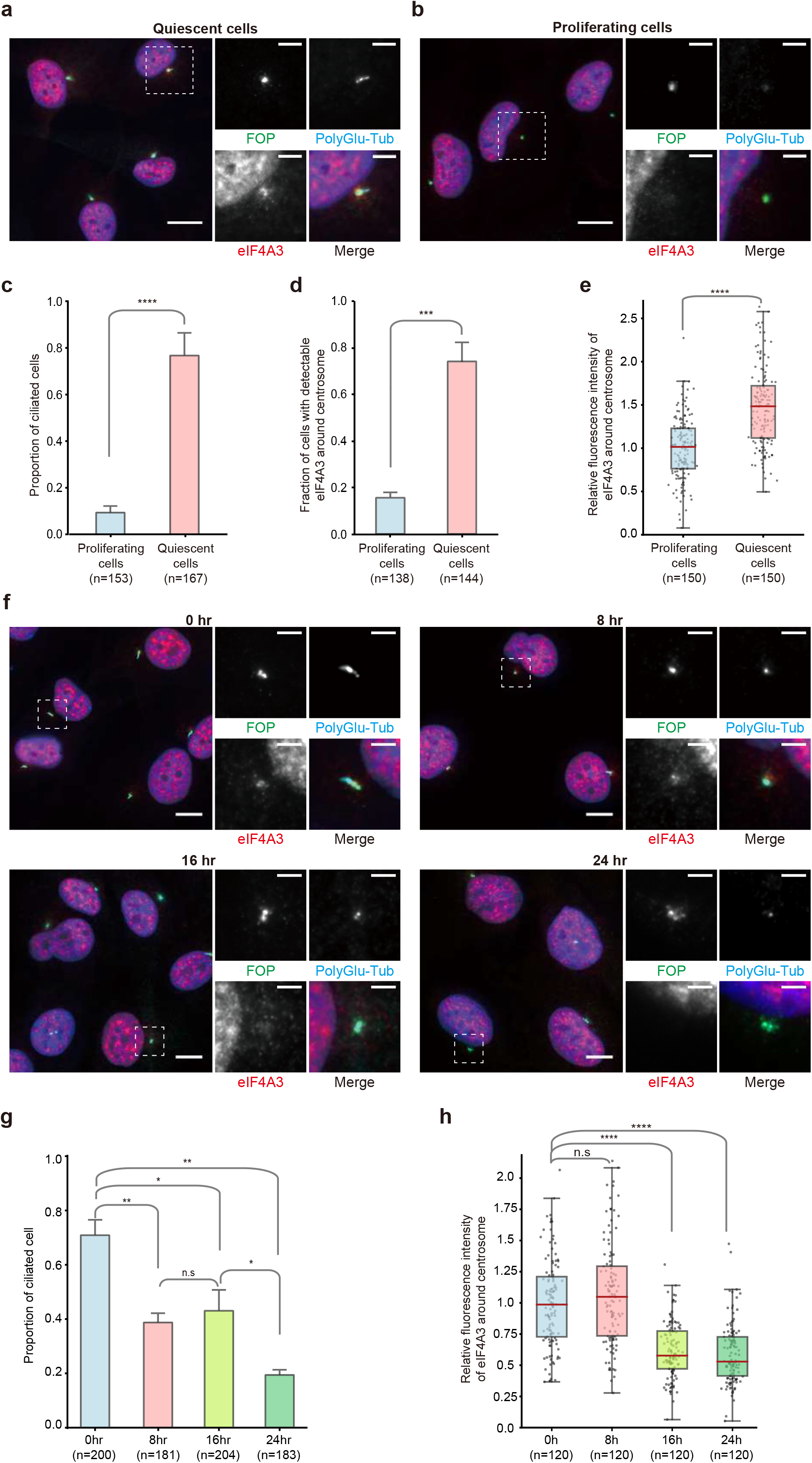
EJC core component accumulates around centrosomes in quiescent RPE1 cells and decreases upon cell cycle re-entry. Proliferating (a) and quiescent (b) RPE1 cells and RPE1 cells incubated with 10 % serum containing media during indicated times after quiescence (f) were stained for eIF4A3. Centrosomes were labeled by FOP antibody and primary cilia, and centriole were stained by poly-glutamylated tubulin antibody. Nuclei were stained by Hoechst. Right panels show enlarged images of the white dashed square in the left panel. Scale bars in the left panels are 10 μm, and scale bars in right panels are 3 μm (a, b, f). The proportion of ciliated cells was determined in either proliferating or quiescent cell populations (c) and the cell populations incubated with serum containing media during indicated incubation times (g). The fraction of cells with detectable eIF4A3 (d) was determined in either proliferating or quiescent RPE1 cells. Quantifications of eIF4A3 fluorescence intensities (e, h) were performed as described in the legend of figure 1 except that average fluorescence intensities of eIF4A3 in proliferating cells (e) and cells with 0 hr incubation (h) are set to 1.0. Error bars correspond to S.D. n.s P > 0.05, * P ≤ 0.05, ** P ≤ 0.01, *** P ≤ 0.001, and **** P ≤ 0.0001, Mann-Whitney test (e, h) and two-tailed t-test (c, d, g). Three independent experiments were performed.

We next investigated cell cycle-dependent localization of EJC proteins. For this, we followed eIF4A3 and Y14 signals during a 24h time-course triggered by serum addition to quiescent RPE1 cells (Fig. 2f-h and Supplementary Fig. 3a, b). As previously reported^57^, the proportion of monociliated cells decreased following a two-steps mode, with roughly 20% of ciliated cells left after 24 hours (Fig. 2g). The amount of eIF4A3 and Y14 started to decrease after 8 hours of serum addition and was similar to the amount observed in unsynchronized proliferating cells (Fig. 2f, h and Supplementary Fig. 3a, b). As the number of cells in S phase peaked at 16 hours after serum addition, accumulation of EJC proteins around centrosomes most likely declined during the S phase (Supplementary Fig. 3c, d).

Serum starvation is a kind of stress that can induces translational repression^58–61^. A short sodium arsenite treatment induced the formation of stress granules detected by TIA-1 and eIF4E antibodies (Supplementary Fig. 4a, b). However, EJC accumulation around centrosome did not correspond to stress-induced foci because serum starvation did not lead to accumulation of the stress granule protein TIA-1 in RPE1 cells. We also investigated the impact of translation inhibition by incubating RPE1 cells with either puromycin that dissociates translating ribosomes or cycloheximide that stalls elongating ribosomes onto mRNAs. Both treatments weakly increased the centrosomal accumulation of eIF4A3 and Y14 in quiescent RPE1 cells and had little effect in proliferating cells (Supplementary Fig. 4c-j). Therefore, the concentration of EJC proteins at the base of RPE1 primary cilia does not result from stress or partial translation inhibition triggered by serum starvation.

### RNA-dependent accumulation of assembled EJCs around centrosomes

One question raised by these results was whether eIF4A3 and Y14 accumulate around centrosomes independently or not. Dual labeling of eIF4A3 and Y14 showed that they colocalize around centrosomes (Fig. 3a). The relative fluorescence intensities of eIF4A3 and Y14 followed similar patterns when plotted along lines crossing either nuclear speckles where the EJC subunits are concentrated (Fig. 3b), or centrosomes (Fig. 3c). Analysis of 60 individual centrosomes and speckles indicated a very high correlation of localization of the two proteins in both places (Fig. 3d). To further support the hypothesis that eIF4A3 and Y14 co-exist in assembled EJCs near centrosomes in quiescent cells, we down-regulated the expression of either eIF4A3 or Y14 by RNA interference. RT-qPCR, Western blotting and immunofluorescence monitoring showed that silencing of one protein did not affect the expression of the other one (Fig. 3e-i and Supplementary Fig. 5a, b). In contrast, down-regulation of Y14 strongly reduced eIF4A3 intensity around centrosomes (Fig. 3h, j), and conversely down-regulation of eIF4A3 strongly reduced Y14 accumulation around centrosomes (Supplementary Fig. 5a, c).

**Figure 3.**
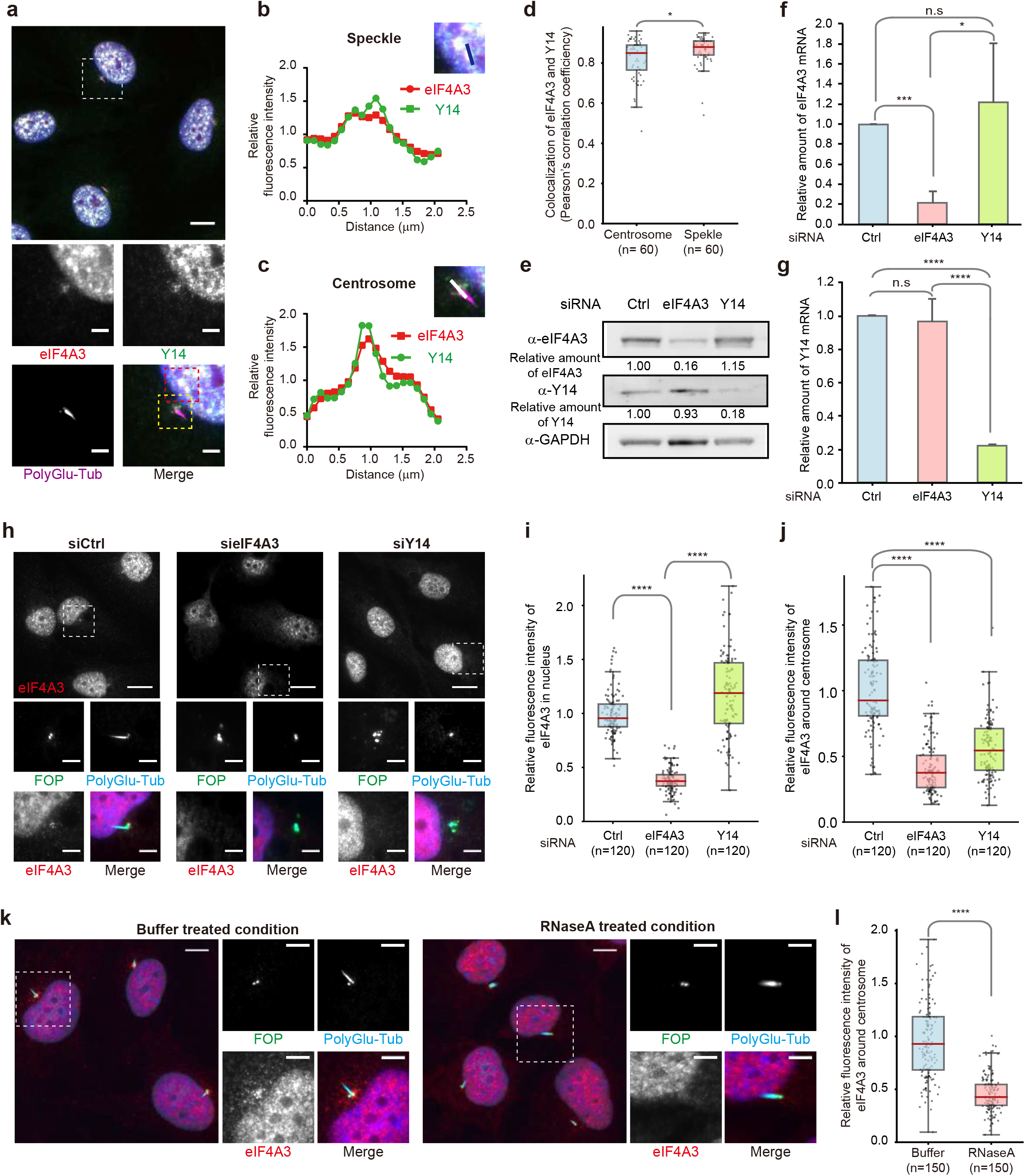
RNA-dependent assembled EJCs localize around centrosomes in quiescent cells. Quiescent RPE1 cells were immunolabeled by eIF4A3 and Y14 (a). Quiescent RPE1 cells transfected with indicated siRNAs (h) and permeabilized quiescent cells incubated with RNAse A or not prior to fixation (k) were stained for eIF4A3. Primary cilia and centrioles were stained by poly-glutamylated tubulin antibody (a, h, k) and centrosomes were labeled by FOP antibody (h, k). Nuclei were stained by Hoechst. Lower (a, h) or right panels (k) show enlarged images of the white dashed square in upper (a, h) or left panel (k). Images from red and yellow dashed squares in lower panel (a) are depicted in b and c, respectively. Scale bars in upper (a, h) or left (k) panels and lower (a, h) or right panels (k) panels are 10 μm and 3 μm, respectively. Relative fluorescence intensity of eIF4A3 and Y14 along the line on nuclear speckle (b) and centrosome (c) were plotted, and average fluorescence intensity on the line is set to 1.0. Colocalization of eIF4A3 and Y14 was analyzed in a 2 μm circle around centrosomes and nuclear speckles (d) and plotted as described in figure 1 except that perfect colocalization is set to 1.0. Knock down efficiency of siRNAs was determined by either Western blotting (e) or RT-qPCR (f, g). Relative protein (e) or RNA level of eIF4A3 (f) and Y14 (g) normalized by GAPDH is depicted. Error bars correspond to S.D. Relative fluorescence intensities of eIF4A3 in the nucleus (i) and those for eIF4A3 around centrosome (j, l) were performed as described in the legends of figure 1 and supplementary figure 1 except that the average fluorescence intensity of eIF4A3 in Ctrl siRNA (i, j) or buffer (l) treated cells is set to 1.0. n.s P > 0.05, * P ≤ 0.05, *** P ≤ 0.001, and **** P ≤ 0.0001, Mann-Whitney test (d, i, j, l) and twotailed t-test (f, g). Three independent experiments were performed.

Since EJCs are assembled onto RNA^12,51^, we next investigated whether their presence around centrosomes depends on RNA. Quiescent RPE1 cells were permeabilized, incubated with RNase A before fixation and stained with antibodies. As a positive control, we showed that the number of P-bodies (cytosolic RNP granules involved in mRNA storage^62^) was reduced by 4 fold upon such treatment (Supplementary Fig. 5d-f)., This short RNaseA treatment slightly reduced the amount of eIF4A3 and Y14 in the vicinity of these foci (Supplementary Fig. 5j, k), as expected given that EJCs are assembled around nuclear speckles^51^. Remarkably, RNase A strongly reduced the amount of both eIF4A3 and Y14 around centrosomes (Fig. 3k, l and Supplementary Fig. 5g-i).

Together, the interdependent centrosomal colocalization of eIF4A3 and Y14, and its susceptibility to RNase strongly support that these proteins accumulate around centrosomes of quiescent cells as part of assembled EJC complexes.

### Microtubule-dependent transport of centrosomal EJCs

Multiple mechanisms allow the transport and/or the concentration of transcripts in specific cellular locations^63^. Active transport of mRNP particles notably use cytoskeleton structures^64,65^. Centrosomes function as the major microtubule-organizing centers^42^. Therefore, we first questioned whether the accumulation of EJCs around centrioles in quiescent cells relies on the microtubule network. When quiescent RPE1 cells were treated with nocodazole for two hours, a well-known microtubule destabilizer, microtubules disappeared (Supplementary Fig. 6a, b). However, β-tubulin immunostaining of both centrioles and cilia was not significantly affected (Fig. 4a). In contrast, this treatment reduced the fluorescence intensities of eIF4A3 and Y14 around centrosomes, by 60% and 50% respectively (Fig. 4a, d and Supplementary Fig. 6d, g). These observations indicate that EJCs accumulate around centrosomes in a microtubule-dependent manner. Given that centrosome nucleate the minus-ends of microtubules, minus-end directed motors and notably cytoplasmic dynein might transport EJC-bound particles to centrosomes. To test this hypothesis, we incubated quiescent RPE1 cells for 90 min with Ciliobrevin D, a cell-permeable inhibitor of dynein. Indeed, such treatment efficiently disrupted the Golgi network immunostained with GM130 antibodies, as previously reported^66^ (Supplementary Fig. 6c). Interestingly, the Ciliobrevin D treatment also reduced the fluorescence intensities of eIF4A3 and Y14 around centrosome by 40 % (Fig. 4b, e and Supplementary Fig. 6e, h) showing that centrosomal EJCs concentration requires dynein motors.

**Figure 4.**
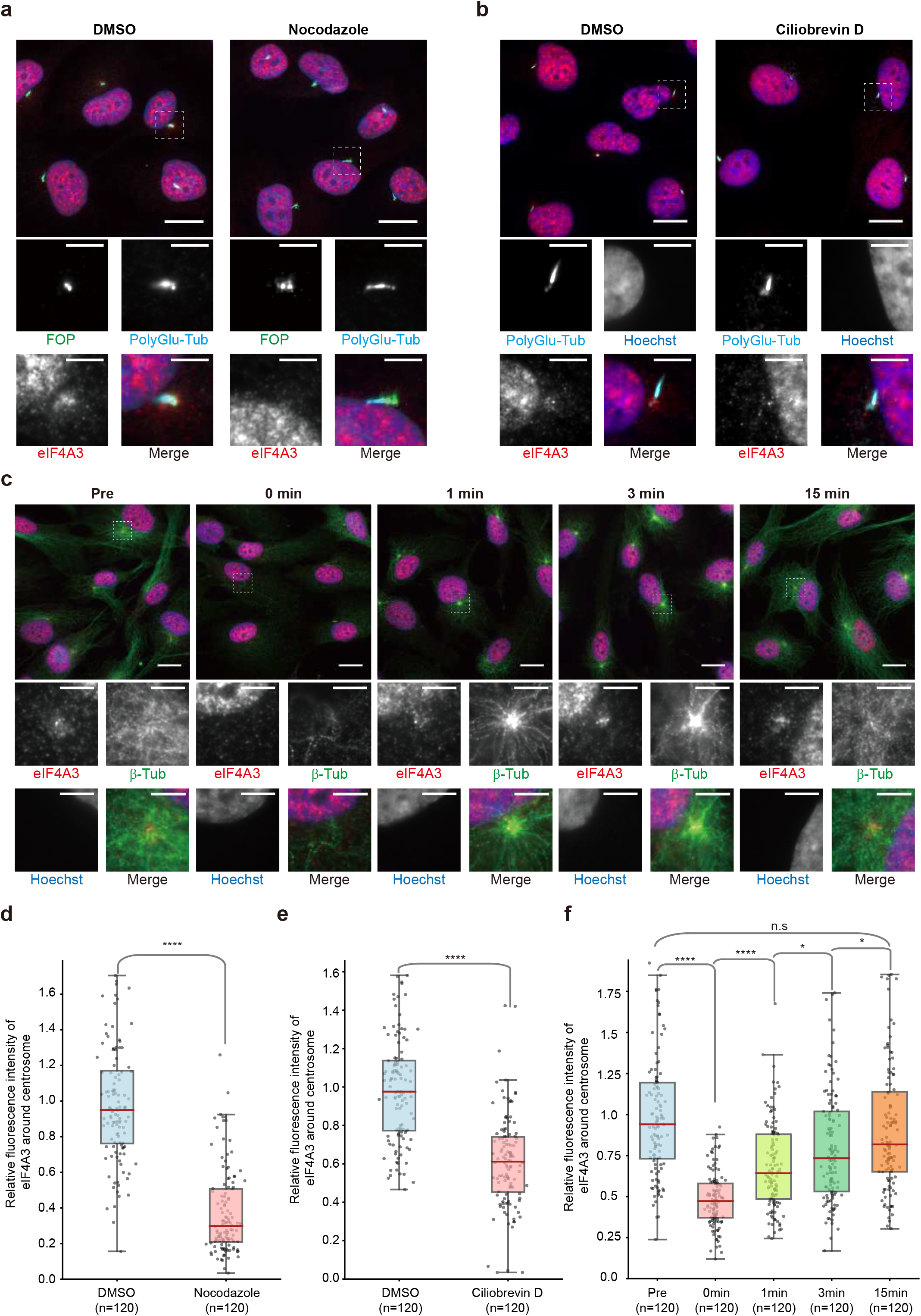
An active microtubule-dependent transport is required to maintain EJC localization around centrosomes. eIF4A3 antibody stained quiescent RPE1 cells treated with either DMSO or Nocodazole (a), either DMSO or CiliobrevinD (b), and chilled quiescent cells subjected to a microtubule regrowth assay (c). Centrosomes were labeled by FOP antibody and primary cilia and centriole were stained by poly-glutamylated tubulin antibody (a, b). Microtubules were stained by β-tubulin antibody (c). Nuclei were stained by Hoechst. Lower panels show enlarged images marked by white dashed square in the upper panel. Scale bars in the upper and lower panels are 10 μm and 3 μm, respectively. Quantification of fluorescence intensities of eIF4A3 (d-f) were performed as described in the legend of figure 1. The average fluorescence intensities for eIF4A3 in DMSO treated cells (d, e) or in pre-incubated quiescent cells (f) are set to 1.0. n.s P > 0.05, * P ≤ 0.05, and **** P ≤ 0.0001, Mann-Whitney test. Three independent experiments were performed.

Microtubules form a dynamic network undergoing permanent polymerization and depolymerization. To further investigate the dynamic aspect of EJC transport to centrosomes, we performed microtubule regrowth assays. When quiescent cells were ice-chilled, the microtubule network labeled with β-tubulin or α-tubulin antibodies almost completely disappeared, and the amounts of centrosomal eIF4A3 and Y14 were reduced by 2 fold (Fig. 4c, f and Supplementary Fig. 6f, i). Placing cells back at 37°C induced microtubule regrowth. Already one minute after addition of 37°C media, astral structures reappeared at microtubule organizing centers and 15 minutes later, the microtubule network was almost completely reconstituted (Fig. 4c and Supplementary Fig. 6f). Remarkably, the intensity of eIF4A3 and Y14 at centrosome already increased after one minute back at 37°C and reached almost initial levels after 15 minutes (Fig. 4c, f and Supplementary Fig. 6f, i).

Together, these data indicated that an important proportion of EJC complexes are rapidly transported to centrosomes of quiescent RPE1 in a microtubule- and dynein-dependent manner.

### Basal body localization of *NIN* mRNAs but not *BICD2* mRNAs is both EJC- and translation-dependent

Finding EJCs assembled on RNA prompted us to search for transcripts localized around centrosomes in ciliated RPE1 cells. We used a high-throughput smFISH strategy (schematized in Fig. 5a; see also Safieddine et al.^67^) to screen about 700 mRNAs encoding centrosome- and cilium-related proteins (Supplementary Table 1). Briefly, we generated 50 to 100 distinct single-stranded RNA probes for each mRNA. The probes were flanked by two overhangs that hybridize with fluorescently labeled locked nucleic acids (LNA). The probe mixtures were hybridized on fixed cells following a smiFISH procedure as described previously^68^. We used RPE1 cells stably expressing centrin1-GFP for centrosome labeling and antibodies against Arl13b to stain primary cilia (Fig. 5a and Supplementary Fig. 7a). Among the different mRNAs investigated, we found 21 mRNAs that exhibit non-random intracellular distribution (Supplementary table 1). For example, *CHD3* (Chromodomain Helicase DNA binding protein 3; a component of NuRD chromatin remodeling complex^69^) accumulated in cytoplasmic protrusions while *NEK9* mRNA (also known as Nercc1; serine/threonine kinase controlling centrosome separation during prophase^70^) was distributed randomly throughout the cytoplasm (Supplementary Fig. 7a). Among the different mRNAs investigated, 18 mRNAs exhibited non-random intracellular distribution. Remarkably, two mRNAs, *BICD2* and *NIN*, specifically concentrated around centrosomes at the base of cilia (Supplementary Fig. 7a). Bicaudal D2 (BICD2) is a dynein adaptor involved in RNP particles and vesicles trafficking along microtubule network^71^. Ninein (NIN) is a core component of centrosomes required for microtubule nucleation and anchoring to centrosome^72,73^. While we screened a large fraction of mRNAs corresponding to the centrosomal and cilium proteomes, the screen was not exhaustive. Hence, additional mRNAs might localize there (see Discussion).

**Figure 5.**
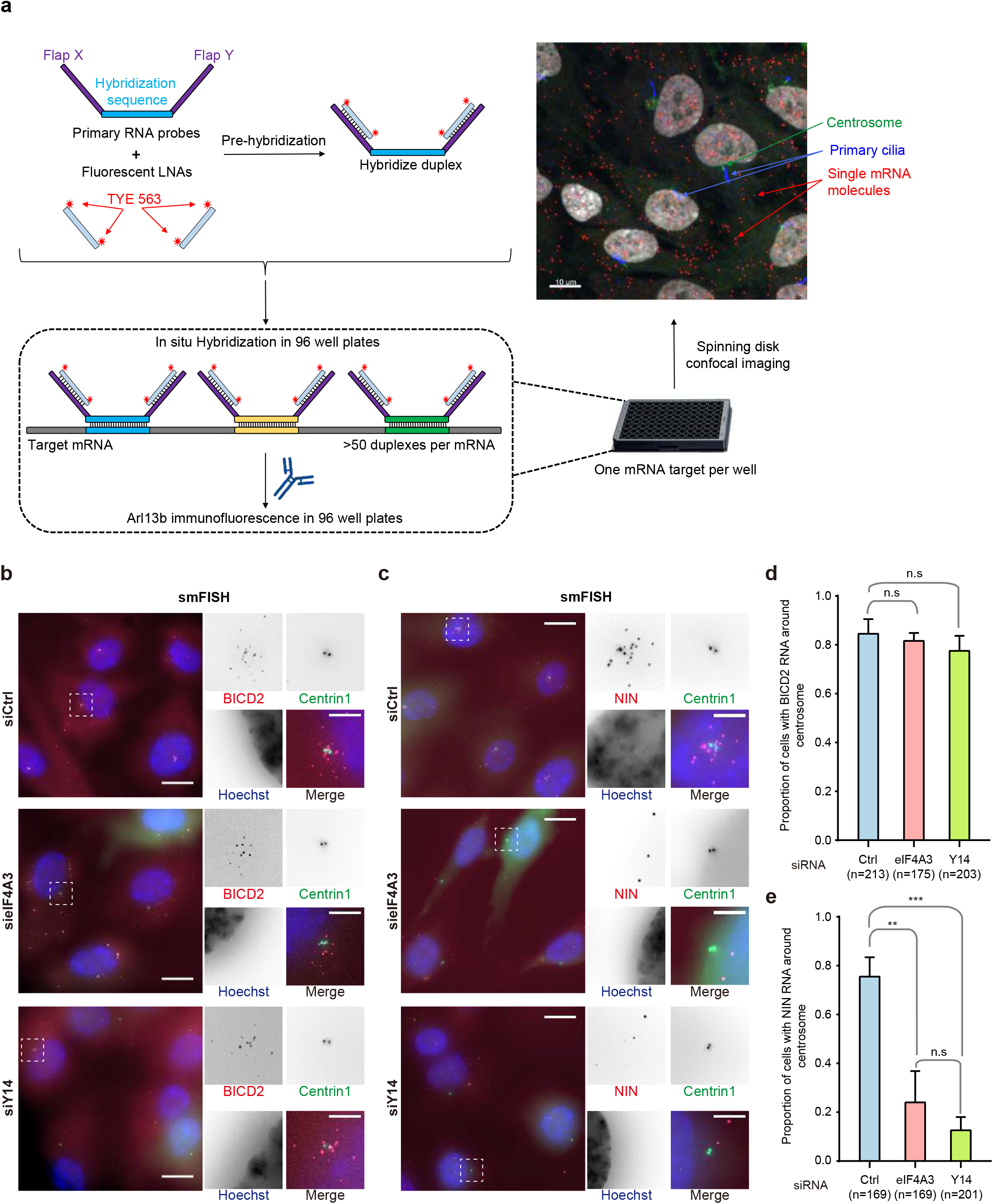
EJC is required for centrosomal localization of *NIN* mRNA. Summary of the high-throughput smiFISH pipeline (a). Top left: primary RNA probes contain a hybridization sequence that is complementary to the target mRNA flanked by two overhangs named Flap X and Y. Each Flap was annealed to a locked nucleic acid (LNA) oligo labeled with two TYE 563 molecules in a pre-hybridization step. Bottom: Duplexes were then hybridized to the mRNA of interest followed by immunofluorescence against Arl13b to label primary cilia in 96 well plates. Plates were finally imaged with a spinning disk confocal microscope. Top right: A micrograph showing a typical field of view from the screen. Red dots correspond to a single mRNA molecule. Scale bar represents 10 μm. Quiescent RPE1 cells stably expressing centrin1-GFP were stained by probes against *BICD2* mRNA (b) or *NIN* mRNA (c) after knock-down of either eIF4A3 or Y14. Nuclei were stained by Hoechst. Images are resulted from maximum intensity projections of 14 z-stacks acquired at every 0.5 μm. Right panels show enlarged images of the white dashed square in the left panels. Scale bars in the left panels are 10 μm, and scale bars in right panels are 3 μm (b, c). Proportion of cells displaying centrosomal *BICD2* (d) or *NIN* (e) RNA pattern was depicted. Error bars correspond to S.D. n.s P > 0.05, ** P ≤ 0.01, and *** P ≤ 0.001, two-tailed t-test. Three independent experiments were performed.

Next, to determine whether *BICD2* and *NIN* transcripts were associated to EJCs, we performed RNA immunoprecipitations. RT-qPCR showed that both transcripts were efficiently and specifically precipitated with eIF4A3 and Y14 antibodies but not with antibodies against the unrelated protein Rab5 (Supplementary Fig. 8a). This enrichment was specific because the intron-less *SFM3B5* and *SDHAF1* transcripts were not precipitated under the same conditions (Supplementary Fig. 8a). Therefore, a significant proportion of *BICD2* and *NIN* mRNAs are bound to EJCs in quiescent RPE1 cells.

We next tested whether the localization of *BICD2* and *NIN* mRNAs was EJC-dependent. Neither eIF4A3 or Y14 knock-down affected the centrosomal localization of *BICD2* mRNA (Fig. 5b, d). In contrast, both knock-downs strongly perturbed *NIN* mRNA localization that became more dispersed (Fig. 5c, e). Measurement of *NIN* mRNAs expression by RT-qPCR showed that EJC knock-downs did not alter its overall expression (Supplementary Fig. 8b). Therefore, EJCs actively participates to the centrosomal localization of *NIN* transcripts.

We and others previously observed that the localization of *PCNT* and *ASPM* mRNAs to centrosome during early mitosis is translation-dependent^74,75^. Although cycloheximide treatment had no effect, puromycin treatment prevented the accumulation of both *NIN* and *BICD2* mRNAs around centrosomes in quiescent RPE1 cells (Supplementary Fig. 7b-e).

Taken together, our data suggest that EJCs contribute to the transport and localization at centrosomes of *NIN* transcripts undergoing translation, whereas the localization of *BICD2* mRNA only requires translating ribosome.

### EJC protein depletion impairs both centrosome organization and ciliogenesis

Ninein (NIN) is a core component of centrosomes located at the proximal end of each centriole and at sub-distal appendages of mother centrioles. It contributes to microtubule nucleation and anchoring to centrosomes^72,73^. We investigated the NIN protein by immunofluorescence. We found that either eIF4A3 or Y14 knock-down reduced by half the amount of NIN protein detected around centrosomes (Fig. 6a, b). This observation prompted us to analyze other centrosomal components. We observed that knock-down of either eIF4A3 or Y14 also had a strong effect on PCM-1 and FOP localizations in quiescent RPE1 cells (Fig. 6c-e). In control cells, PCM-1 labeling showed that the PCM and centriolar satellites were mainly concentrated around centrioles and radially distributed from the centrosome in a punctuated manner. eIF4A3 knock-down led to the appearance of PCM-1 dots a few microns away from centrioles, with a dispersed and scattered pattern. A Y14 knock-down had the same impact though less pronounced. In control cells, FOP staining was very focused although an extended punctuated staining is seen in 15% of the cells. However, such an extended punctuated FOP staining was observed in 80% of the cells after knock-down of either eIF4A3 or Y14 (Fig. 6c, d). The specific role of EJC in centrosome organization was further supported by the fact that down regulation of MAGOH showed similar effects (Supplementary Fig. 9). In contrast, the down-regulation of MLN51 showed no impact suggesting that EJC-related functions of MLN51 are most likely transcript and/or cell-specific, as previously reported^38,76^. Thus, EJC depletion triggers defects in centriolar satellite transport.

**Figure 6.**
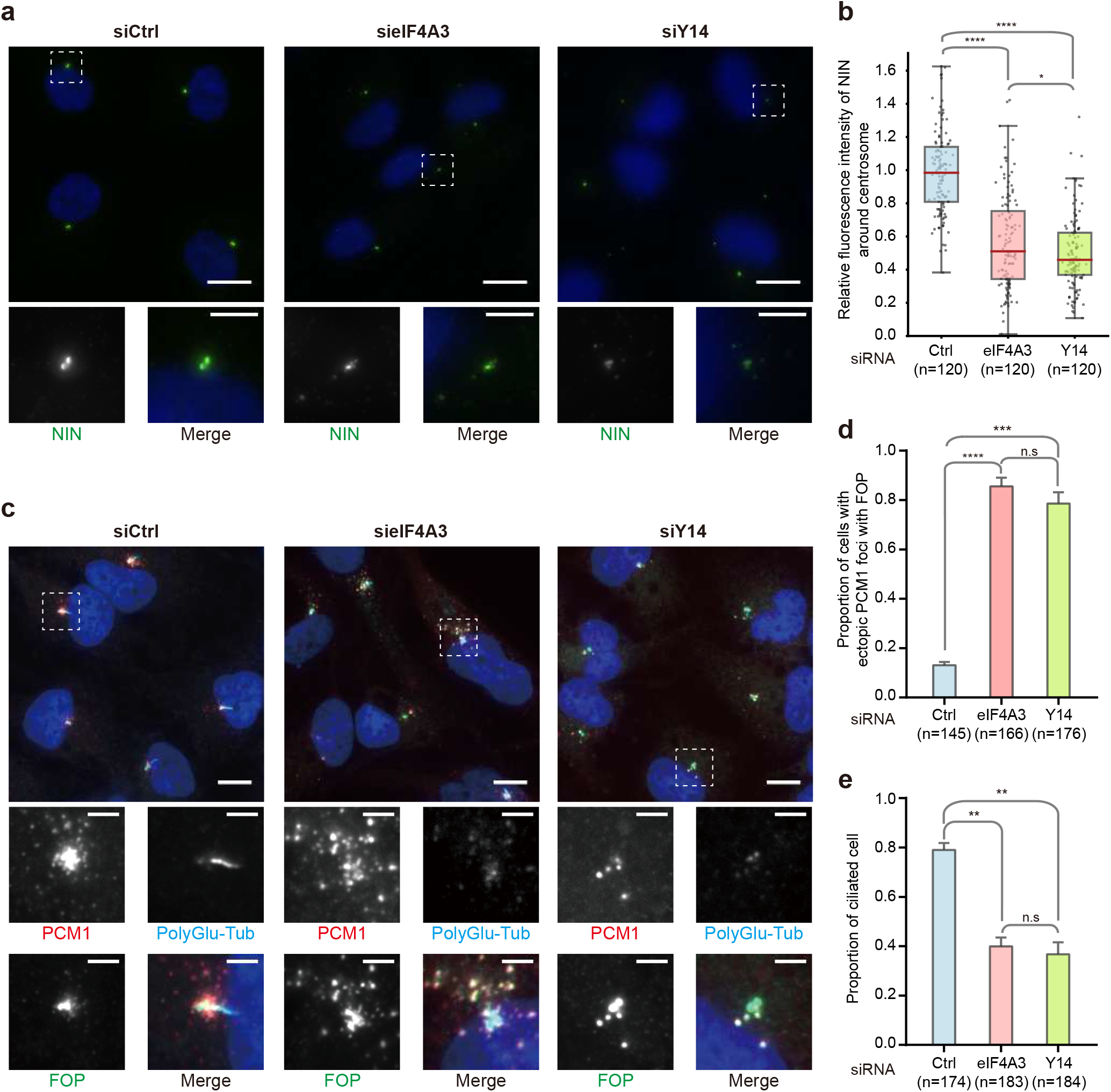
Knock-down of EJC components impairs centrosome structure and primary cilia formation. Quiescent RPE1 cells transfected with siRNAs against eIF4A3 or Y14 were stained for NIN (a) or PCM1 (centriolar satellite protein), FOP, and poly-glutamylated tubulin (c). Nuclei were stained by Hoechst. Lower panels are enlarged images marked by white dashed square in the upper panels. Scale bars in the upper and lower panels are 10 μm and 3 μm, respectively (a, c). Images were processed by maximum intensity projections of 15 z-stacks acquired at every 0.5 μm (a). Quantification of fluorescence intensities of NIN were performed as described in the legend of figure 1. The average fluorescence intensity of NIN in Ctrl siRNA treated cells is set to 1.0 (b). Proportion of cells with ectopic centriolar satellite with FOP upon the siRNA treatments indicated (d). Proportion of ciliated cells upon the indicated siRNA treatments (e). Error bars correspond to S.D (d, e). n.s P > 0.05, ** P ≤ 0.01, *** P ≤ 0.001, and **** P ≤ 0.0001, Mann-Whitney test (b) and Two tailed t-test (d, e). Three independent experiments were performed.

We also observed that eIF4A3 or Y14 knock-downs reduced both the γ-tubulin and PCNT fluorescence intensities at centrosomes (Supplementary Fig. 8c-e), in agreement with the fact that NIN and PCM-1 are important for deposition of γ-tubulin and PCNT on centrosome, respectively^72,77^. However, although EJC knock-down impairs the localization of some centrosome components, it did not induce major changes in the microtubule network revealed by β-tubulin labeling (Supplementary Fig. 8f).

Ciliogenesis is linked to basal body integrity. As knock-down of either eIF4A3 or Y14 decreased the number of ciliated cells by more than 50% (Fig. 6c, e), an imbalance in EJC dosage in quiescent RPE1 cells strongly impairs both the organization of centrosomes and ciliogenesis.

## Discussion

Here, we show the accumulation of EJC core proteins around basal bodies, which are formed by centrioles at the base of cilia. This was observed both in primary mNSCs and quiescent RPE1 cells. These EJC proteins are assembled on RNA and localized in a microtubule-dependent manner revealing the enrichment of untranslated or partially translated EJC-bound transcripts at centrosomes. A large smFISH screen identifies *BICD2* and *NIN* mRNAs near the base of primary cilia in quiescent RPE1 cells. Knock-down of any EJC core protein prevents *NIN* mRNAs transport but not *BICD2* mRNAs transport. Thus, the EJC plays a crucial role for the spatial enrichment of specific mRNAs at a specific location in human cells. In addition, we provide evidences that an EJC imbalance affects the centrosomal accumulation of *NIN* protein and other structural components such as pericentrin and PCM1. Thus, the EJC is associated to defects in centrosomal organization and ciliogenesis.

We provide several complementary evidences that assembled EJCs accumulate in a RNA-dependent manner around basal bodies. EJCs are deposited by the nuclear splicing machinery and remain stably bound to transcripts until their translation in the cytoplasm^15,17^ So far, there is no evidence that a splicing-independent EJC assembly may occur. Therefore, the RNA-dependent enrichment of EJCs at the centrosomes signals the local concentration of spliced transcripts both in mNSC and in human RPE1. Previous studies reported the presence of RNAs in the centrosomal area in different organisms including *Tetrahymena pyriformis, Paramecium tetraurelia, Spisula solidissima, Ilyanassa obsolete, Dano rerio, Xenopus laevis* and *Drosophila melanogaster*^74,78–81^.

More recently, four transcripts encoding the central PCM component, Pericentrin (PCNT), Abnormal spindle-like microcephaly-associated protein (ASPM), the nuclear mitotic apparatus protein 1 (NUMA1) and the Hyaluronan Mediated Motility Receptor (HMMR) were detected by single molecule approaches around centrosome of HeLa cells during cell division^67,74,75^. Here, we identified *BICD2* and *NIN* transcripts as two additional mRNAs concentrated around centrosome at the base of cilia in quiescent RPE1 cells. BICD2, an activating adaptor of dynein, participates to the traffic of both Golgi vesicles and RNP particles along microtubules^71^. Ninein, localized to the proximal end of both mother and daughter centrioles, is a core component of subdistal appendages of mother centrioles^73^. Ninein is important for both microtubule anchoring and nucleation at the centrosome^72^. The presence of several transcripts around centrosomes at different stages of the cell cycle^74,75^, the detection of centrosomal EJCs in proliferating cells and their accumulation during quiescence are all echoes of a major, spatially restricted and dynamic post-transcriptional program crucial for the centrosome functions, both during cell division and cilia formation.

Various mechanisms can lead to mRNP enrichment at particular subcellular locations such as an active transport along cytoskeletal tracks, passive diffusion coupled to site specific anchoring or local protection from degradation^63,64,82^. It has long been considered that most localized mRNPs are transported in a translationally repressed state to prevent ectopic expression of the encoded protein and/or favor the assembly of protein complexes. However, there are growing evidences of widespread co-translational transports^83,84^. Recently, a large dual protein-mRNA screen in human cells revealed that the majority of the transcripts displaying specific cytoplasmic locations reach their destination in a translation-dependent manner^75^. Co-translational mRNA transport is notably essential for the targeting of membrane and secreted proteins to the endoplasmic reticulum. In this case, the cytosolic translation of transcripts is arrested after translation of a signal sequence that mediates the transport of the ribosome-bound mRNP to the endoplasmic reticulum where translation resumes after translocation of the nascent polypeptide^85^. The delivery of *PCNT* and *ASPM* mRNAs to centrosomes requires active polysomes as well as microtubules and dynein activity^74,74^, and direct visualization of single polysomes in live cells with the SunTag showed that the ASPM and NUMA1 polysomes are actively transported to mitotic centrosomes^67^. Recently, a large dual protein-mRNA screen in human cells revealed that the majority of the transcripts displaying specific cytoplasmic locations reach their destination in a translation-dependent manner^75^. Here, we complete this list by showing that the accumulation *NIN* and *BICD2* mRNA around centrosome at the base of primary cilia is highly sensitive to puromycin treatment but not to cycloheximide treatment (Supplementary Fig. S7), strongly suggesting that nascent peptides are necessary for correct targeting. BICD2 and NIN proteins are direct partners of dynein^86^ and the N-terminal region of Ninein is important for protein targeting to mother centrioles^72^. It is tempting to speculate that BICD2 and NIN nascent peptides somehow contribute to the co-translational and dynein-dependent delivery of their transcripts to achieve protein synthesis at their final destination.

So far, only one example of transcript localization requiring the EJC is known. It is the *oskar* mRNA that is transported from nurse cells to the posterior pole of *Drosophila melanogaster* oocytes^18^. Here, we report a second EJC-dependent localized transcript, the *NIN* mRNA. It is the first described in mammals, revealing that this phenomenon is not an exception restricted to fly. The EJC is likely involved in the subcellular localization of other transcripts yet to be identified. Multiple cis- and trans-acting factors participate to the active transport and the translational repression of *oskar* mRNP before it reaches the posterior pole of embryo where the protein Osk is produced^63,87,88^. In this multistep pathway, the EJC is only one of the actors and its precise role remains unclear. In contrast to *oskar* mRNA, the localization of *NIN* mRNA requires ongoing translation in addition to the EJC. The combination of these two signals is at first surprising because EJCs deposited on the mRNA ORF are expected to be disassembled by scanning ribosomes. Ribosomes might be halted before reaching the end of the *NIN* mRNA ORF. The differential sensitivity of *NIN* mRNA localization to cycloheximide and puromycin, suggests that a nascent NIN polypeptide bound to halted ribosomes cooperates with downstream EJCs for *NIN* mRNA targeting to centrosome where translation would resume. It is also worth noting that *NIN* mRNA is 10 kb long and has 50 introns, thus possibly requiring a particular EJC-driven packaging for transport. Although the molecular mechanisms involved remain unclear, the differential EJC-requirement for *BICD2* and *NIN* mRNAs localization indicate that different pathways orchestrate the ballet of transcripts accumulating around centrosome during the different phases of the cell cycle^74,75^ (Fig. 7).

**Figure 7.**
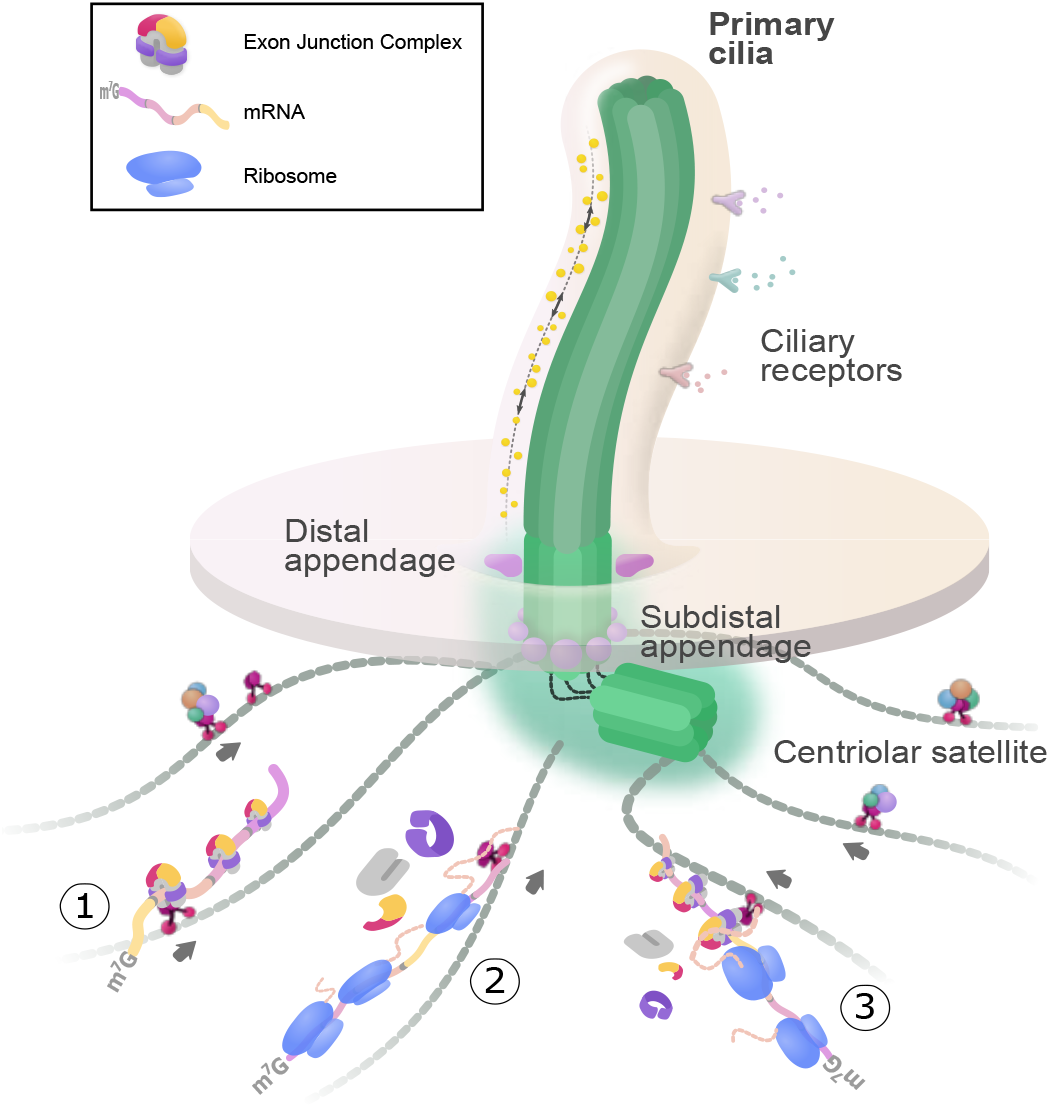
Multiple pathways for RNA localization toward centrosome. During quiescence, mother centriole tightly attaches to plasma membrane by distal appendages to nucleate the formation of primary cilia. To relay the signals from environment, several receptors are embedded at ciliary membrane. Centriolar satellites deliver components of centrosome and primary cilia. Centrosomal transcripts could be localized by ➀ EJC and/or other RNPs mediated pathway, ➁ polysome dependent pathway, and ➂ polysome and EJC dependent pathway.

In quiescent cells, the oldest centriole of the centrosome converts into a basal body that nucleates a non-motile primary cilium. This organelle serves as a cellular antenna and constitutes a signaling hub for both chemical and mechanical external stimuli leading to cell-fate decisions such as cell cycle re-entry or cell differentiation^89^. In the brain, primary cilia sense signaling molecules present in the cerebral spinal fluid^90^. Untranslated and partially translated mRNAs unmasked by EJCs and parked around the basal body might wait for external signals to synthesize their protein products on request and contribute to centrosome organization, ciliogenesis and cilia functions. Here, we show that downregulation of EJC core factors in RPE1 cells affects the localization of *NIN* mRNAs and consequently, the amount of NIN protein at centrosomes (Fig. 6). Furthermore, it leads to centriolar satellite scattering, pericentriolar assembly/composition defects and to abnormal ciliogenesis (Fig. 6 and Supplementary Fig. 9). Centriolar satellites are dynamic granules transported towards centrosomes along microtubules by a dynein-dependent mechanism^91^ that are essential for centrosome assembly as well as ciliogenesis^77,92^. Therefore, the targeting of EJC-bound transcripts such as *NIN* mRNA toward centrosomes and possibly their local translation is critical for centrosome structure and cilia formation.

Brain development is particularly susceptible to centrosome dysfunction and defects in several centrosome components are associated with microcephaly^93,94^. Interestingly, mouse haplo-insufficiencies in EJC core factors are all associated with defects in neural stem cell division^29,37^. A reduction in intermediary progenitors and an increased apoptosis of progeny are observed and result in neurogenesis defaults and ultimately microcephaly. In addition, a deficient Ninein expression in embryonic mouse brain causes premature depletion of progenitors^95^. Thus, it is tempting to speculate that EJC-linked neurodevelopmental abnormalities observed in mouse models as well as in human syndromes at least in part originate from centrosomal and primary cilia dysfunctions in NSC, triggered by a defective post-transcriptional EJC-dependent gene regulation.

## Acknowledgements

We thank Alice Lebreton and Dominique Weil for antibodies and Alexandre Benmerah for RPE1 cells. We thank Olivier Bensaude for critical readings of the manuscript and scientific discussions, and Nathalie Delgehyr for scientific inputs. This study was supported by the ANR (Agence Nationale de la Recherche), with the grants differEnJCe (ANR-13-BSV8-0023), spEJCificity (ANR-17-CE12-0021) to HLH, Hi-FISH (ANR-14-CE10-30) to EB, by FRM (Fondation pour la Recherche Médicale; grant bioinformatics) to EB, by LNCC (Ligue Nationale Contre le Cancer to EB (equipe labélisée), by the European Commission (Marie Curie ITN RNPnet) to HLH, the European Research Council (ERC Consolidator grant 647466) to NS, by the program «Investissements d’Avenir» launched by the French Government and implemented by ANR (ANR–10–LABX-54 MEMOLIFE and ANR–10–IDEX–0001–02 PSL* Research University) to HLH and NS, and by continuous financial support from the Centre National de Recherche Scientifique, the Ecole Normale Supérieure and the Institut National de la Santé et de la Recherche Médicale, France.

## Author contribution statement

HLH, NS and OSK conceived the project. RM and MF prepared primary cell cultures of mNSC and their differentiation into ependymal cells. OSK, RM and MF performed immunofluorescence microscopy on mNSC. OSK conducted immunofluorescence microscopy on RPE1 cells. IB performed Western blotting and QA performed RIP. AS and EC performed high-throughput smFISH screen. OSK performed microtubule regrowth assays, Sunset analyses, flow cytometry experiments, smFISH experiments, all RPE1 treatments, image analyses including quantifications. EB, HLH, NS, OSK and RM analyzed the data. OSK prepared the Figures. OSK and HLH wrote the manuscript that was reviewed and edited by EB and NS.

## Materials and methods

### Animals

All animal studies were performed in accordance with the guidelines of the European Community and French Ministry of Agriculture and were approved by the Direction départementale des Services Vétériniares de Paris (Approval number APAFIS#9343-201702211706561 v7). The mice used in this study have already been described and include: RjOrl:SWISS (Janvier Laboratories).

### Primary ependymal cell cultures and differentiation

mNSC and ependymal cells were prepared following previous reports^43,44^ Newborn mice (P0–P2) were killed by decapitation. The brains were dissected in Hank’s solution [10 % Hanks balanced salt solution (GIBCO), 5 % HEPES (GIBCO), 5 % sodium bicarbonate (GIBCO), 1 % penicillin/streptomycin (P/S) (GIBCO)] and the extracted ventricular walls were manually cut into pieces. The telencephalon was incubated in enzymatic digestion solution [DMEM glutamax, 2.8 % (v/v) papain (Worthington 3126), 1.4 % (v/v) of 10 mg/ml DNase I, 2.25 % (v/v) of 12 mg/ml cysteine] for 45 min at 37 °C in a humidified 5 % CO_2_ incubator. Digestion was inactivated by addition of trypsin inhibitors [Leibovitz Medium L15 (GIBCO), 50 μg/ ml BSA, 1 mg/ml trypsin inhibitor (Worthington), 2 % (v/v) 10 mg/ml DNase I (Worthington)]. Cells were washed with L15 medium and resuspended by DMEM glutamax supplemented with 10 % fetal bovine serum (FBS) and 1 % P/S. Ependymal progenitors proliferated until cells are confluent (4-5 days) in a Poly-Llysine (PLL)-coated flask. Then, cells were shacked (250 rpm) at RT overnight before treatment with trypsin-EDTA. Then, 1.5 × 10^5^ −2 × 10^5^ cells were plated on the PLL coated coverslip and cultivated in DMEM glutamax 10 % FBS, 1 % P/S. The next day, medium was replaced by serum-free DMEM glutamax 1 % P/S, to trigger ependymal differentiation gradually *in vitro* (DIV 0). Cells were fixed with 4 % paraformaldehyde at DIV 1 day and DIV 6 day for quiescent mNSC and ependymal cells, respectively.

### RPE1 cell culture and modulations

RPE1 cells were cultivated in DMEM-F12 1:1 (Invitrogen) supplemented with 10 % of fetal bovine serum (FBS, PAN™ BIOTECH), and 1 % penicillin and streptomycin. To induce quiescence, RPE1 cells were washed twice with DPBS and incubated for 48 hours with serum free DMEM-F12^96^. To repress protein synthesis 100 μg/ml of cycloheximide (TOKU-E) and 300 μM of puromycin (InVivoGen) were added for 2 hours before EJC IF. 100 μg/ml of puromycin was added for 20 mins before smiFISH. To disrupt microtubules, 3 μg/ml of nocodazole (Sigma) in DMSO was added to the cells. To prepare immunofluorescence sample, 5 × 10^4^ of RPE1 cells were plated on PLL coated coverslips (VWR) one day before quiescence induction and 3 × 10^4^ cells were plated for proliferating condition.

### Microtubule regrowth assay

To destabilize the microtubule structure in the cell, quiescent RPE1 cells were left on ice for 30 min. The cold medium was next replaced with pre-warmed medium and the cells were incubated in 37°C. After indicated incubation times, cells were washed in PBS and fixed.

### siRNA transfection

RPE1 cells plated on day 0 were transfected on day 1 with control siRNAs (5’-UGAAUUAGAUGGCGAUGUU-3’), eIF4A3 siRNA (5’-AGACAUGACUAAAGUGGAA-3’), and Y14 siRNA (5’-GGGUAUACUCUAGUUGAAUUUCAUAUUCAACUAGAG-3’) with Lipo2000 (Invitrogen) in Optimem (Gibco). After three hours, cells are replaced in DMEM with FBS and processed on day 4. If required, cells were serum-starved on day 2.

### Antibodies

For Western blot: α-Puromycin (Merck 1:12,500), α-mouse antibody conjugated with HRP (Bethyl). For immunofluorescence: α-FOP (Abnova, Mouse IgG2b, 1:1000), α-Polyglutamylated tubulin (ADIPOGEN, Mouse IgG1, 1:500), α-Y14 (Santacruz, mouse IgG2b, 1:50), α-eIF4A3 (affinity purified from rabbit serum^97^, 1:2000), α-MLN51 (affinity purified from rabbit serum^97^, 1:500), α-β tubulin (Biolegend, Mouse 1:1000), α-EDC4 (Santacruz, Mouse, 1:1000), α-DDX6 (Novus, Rabbit, 1:1000), α-Pericentrin (Covance, Rabbit 1:500), α-γ-tubulin (Sigma, Mouse 1:500), α-PCM1(Cell signaling, Rabbit 1:600), α-9G8 (described previously^98^, Rabbit 1:1000), α-SC35 (described previously^98^, Mouse 1:1000), α-NIN (Institut curie, Human 1:200), anti-Arl13b (proteintech, Rabbit, 1:4500) α-Rabbit Alexa594 (ThermoFisher, 1:400), α-Mouse IgG1 Alexa488 (ThermoFisher, 1:500), α-Mouse IgG2b Alexa647 (ThermoFisher, 1:400), α-Mouse Alexa488 (Thermofischer, 1:500), α-Rabbit Cy5 (Jackson ImmunoResearch, 1:800).

### SUnSET analysis

Cells were incubated with 4.5 μM Puromycin (InvivoGen) for 15 min at 37°C. After washing with PBS, cells were scraped, pelleted at 0.5 rcf and the pellet was lysed in RIPA buffer (20 mM Tris-Cl pH 7.5, 150 mM NaCl, 1 mM Na2EDTA, 1 mM EGTA, 1 % NP-40, 1 % Sodium deoxycholate) supplemented with protease inhibitor mix (Millipore). Protein concentrations were determined by Bradford protein assay. 25 μg of proteins was electrophoresed in a 12 % SDS-polyacrylamide gel. Proteins were electrotransferred onto a 0.2 μm nitrocelluose membrane (GE Healthcare) and blocked with 5 % skim milk in 0.1 % Triton in Tris Buffer Saline (T-TBS) for 30 min at RT. The membranes were incubated overnight at 4 °C with α-Puromycin antibody in blocking solution. An α-mouse antibody conjugated with HRP was incubated for 2 hours at RT. Puromycilated peptides were visualized by chemiluminescence with SuperSignal West Pico PLUS (Thermo Scientific). Total protein on the membrane was stained with PierceTM reversible protein stain kit (Thermo) by manufacturer’s instruction.

### Flow cytometry

Cells were trypsinized and resuspended with DPBS. Resuspended cells were permeabilized by incubating with extraction buffer (0.2 % Triton X-100 in PBS) for 5 min in ice, fixed by incubating with fixation buffer (2 % PFA in PBS) for 15 min in RT and stored in storage buffer (3 % FBS, 0.09 % sodiumazide in DPBS). Cells were labeled by 10 μg/ml of HOECHST 33258 for 30 min in RT. Cell cycle of cells (> 3 × 10^4^) were determined by HOECHST fluorescence in single cell by ZE5 cell analyzer (BioRad) and data were processed by Flowjo.

### Immunofluorescence

Cells on coverslips were washed with DPBS, fixed with 4 % paraformaldehyde in PBS for 10 min at RT, permeabilized with 0.1 % Triton X-100 in PBS for 2 min at RT, blocked with 1 % BSA in PBS for 30 min in RT, incubated for 1 hr with primary antibodies diluted in blocking solution and nuclei were stained with HOECHST 33258, 1 μg/ml in blocking solution for 5 min at RT. Coverslips were next incubated with secondary antibodies for 1 hr in RT and mounted with Fluoromount-G (Invitrogen) on slideglasses. For RNaseA treatment, RNAse A (Sigma) was prepared by dissolving 10 mg/ml RNaseA in 10 mM Tris-Cl pH 7.5, 15 mM NaCl. To inactivate contaminating DNases, the RNaseA solution was heated at 98°C for 15 min. Coverslips were washed with PBS followed by a wash with CSK buffer (10 mM PIPES pH 7.0, 100 mM NaCl, 300 mM sucrose, 3 mM MgCl2) and permeabilized with 0.5% Tween-20 for 5 min at RT. Coverslips were incubated with 5 mg/ml of RNaseA at 37°C for 15 min and next washed two times with PBS followed by 10 min incubation with 4 % paraformaldehyde in PBS. Cells were additionally washed with PBS for three times and immunofluorescence was performed as described above.

### High-throughput single molecule inexpensive fluorescent *in situ* hybridization with immunofluorescence (HT-smiFISH-IF) and conventional smiFISH

The 711 screened genes were selected by their GO term in the “Component” categorie. We included all human genes whose GO Component included one of the following term: “centrosome”, “centriole”, “pericentriolarmaterial”, “cilium”, “microtubule”,”equatorialcellcortex”, “midbody”, “spindle”, “mitoticspindle”, “celldivisionsitepart”. This represents 732 human genes and data were obtained for 711 of them.

RPE1 cells stably expressing centrin1-GFP were seeded in 96-well glass bottom plates (SensoPlates, Greiner) and induced quiescence the next day by a 24 hours culture in Dulbecco’s modified Eagle’s Medium (DMEM, Gibco) with 0.25% % fetal bovin e serum (FBS, Sigma-Aldrich). Cells were then directly fixed for 20 min at RT with 4% paraformaldehyde (Electron Microscopy Sciences) diluted in PBS, and permeabilized with 70% ethanol overnight at 4°C.

To generate primary RNA probes used in the high-throughput smiFISH screen and conventional smiFISH experiments, a pool of DNA oligonucleotides (GenScript) was used. The oligonucleotide design was based on the Oligostan script^68^ with each oligo having a gene-specific segment that will hybridize to the mRNA of interest, flanked by two common overhangs named Flap X and Flap Y Sequence of probes used in screen is depicted in Supplementary table 2. Briefly, a first series of PCR was performed using gene-specific barcodes placed at the extremities of each oligo to amplify specific probe sets using a hot start Phusion DNA Polymerase (Thermo Fisher Scientific, F549L). A second series of PCR was done to add the T7 RNA polymerase promoter using the following primers: FLAP Y sequence with the addition of the T7 sequence at its 5’ end (5’ TAATACGACTCACTATAGGGTTACACTCGGACCTCGTCGACATGCATT-3’), and the reverse complement sequence of FLAP X (5’-CACTGAGTCCAGCTCGAAACTTAGGAGG-3’). This PCR reaction was carried out with GoTaq G2 hot start DNA Polymerase (Promega, F549L). All PCR reactions were in 96-well plates with a Freedom EVO 200 (Tecan) robotic platform. PCR products were checked by capillary electrophoresis on a Caliper LabChip GX analyzer (PerkinElmer). The products of the second PCR were purified with a NucleoSpin 96 PCR Clean-up kit (Macherey-Nagel), lyophilized, and resuspended in DNase/RNase-free distilled water (Invitrogen). *In vitro* transcription was subsequently performed with T7 RNA Polymerase and the obtained primary probes were analyzed by capillary electrophoresis using a Fragment Analyzer instrument (Advanced Analytical).

50 ng of primary probes (total amount of the pool of probes) and 25 ng of each of the secondary probes (LNA oligonucleotides targeting FLAP X and FLAP Y labeled with TYE 563, Qiagen) were pre-hybridized in either 100μL of 1X SSC for conventional smiFISH, or in the following pre-hybridization buffer: 1X SSC, 7.5 M urea (Sigma-Aldrich), 0.34 μg/mL tRNA, 10% Dextran sulfate. Pre-hybridization was performed on a thermocycler with the following program: 90°C for 3 min, 53°C for 15 min, up until probe usage. Plates with fixed cells were washed with PBS and hybridization buffer (1X SSC, 7.5 M urea). For conventional smiFISH, the pre-hybridized mixture was diluted in the same pre-hybridization buffer as above. Hybridization was then carried out overnight at 48°C. The next day, plates were washed eight (screen) or three (conventional smiFISH) times for 20 minutes each in 1xSSC 7.5M urea at 48°C, followed by three PBS rinses. The samples that were not processed into immunofluorescence were directly mounted on slide glass with Vectashield mounting medium with Dapi (Vector laboratories).

For post HT-smiFISH immunofluorescence, cells were permeabilized with 0.1% Triton-X100 in PBS for 10 minutes at room temperature and washed twice with PBS. For cilia labelling, plates were incubated overnight at 4°C with an anti-Arl13b antibody diluted in 0.1% Triton X-100 PBS. The next day, plates were washed three times with PBS, and incubated with a Cy5-labeled goat anti-rabbit secondary antibody in 0.1% Triton X-100 PBS. After 2 hour of incubation at room temperature, plates were washed three times with PBS. To label DNA, cells were then stained with 1 μg/mL Dapi diluted in PBS, and finally mounted in 90% glycerol (VWR), 1 mg/mL p-Phenylenediamine (Sigma-Aldrich), PBS pH 8.

### Fluorescence microscopy and image analysis

Images were acquired with an epifluorescence microscope (Nikon ECLIPSE Ti) equipped with a plan APO VC 60 X objective (N.A 1.4, Nikon), CCD camera (ORCA Flash 4.0, Hamamatsu) and operated by Micro-Manager (MM studio). Maximum intensity projection of z-stacks was processed by Fiji. HT-smiFISH-IF were were imaged on an Opera Phenix High-Content Screening System (PerkinElmer), with a 63x water-immersion objective (NA 1.15).

Centrosomes are detected by poly-glutamylated tubulin and/or FOP staining. Cilia are detected poly-glutamylated tubulin staining. In multiciliated ependymal cell, centrioles were chosen at random in multiciliated ependymal cell. P-bodies are identified by both of DDX6 and EDC4. The number of P-body per cell was determined in images that are processed by maximum intensity projection from 6 z-stacks acquired at every 1 μm. Fluorescence intensities were measured in 2 μm diameter circles around centrosomes (RPE1, or mouse NSC) or base of cilia (ependymal cells) were determined by Image J. Centrosomes and base of cilia that do not overlap the nucleus were selected to exclude nuclear background interference. The background was defined as the lowest pixel intensity in the circle and subtracted from the average fluorescence intensity. Pearson’s correlation coefficient for colocalization was determined in the selected area by Coloc2 plugin in Fiji. Box plots and bar graphs are made by using matplotlib and GraphPad Prism 7, respectively.

