## Supplementary Figures for "Exon Junction Complex dependent mRNA localization is linked to centrosome organization during ciliogenesis"

### Fig.S1

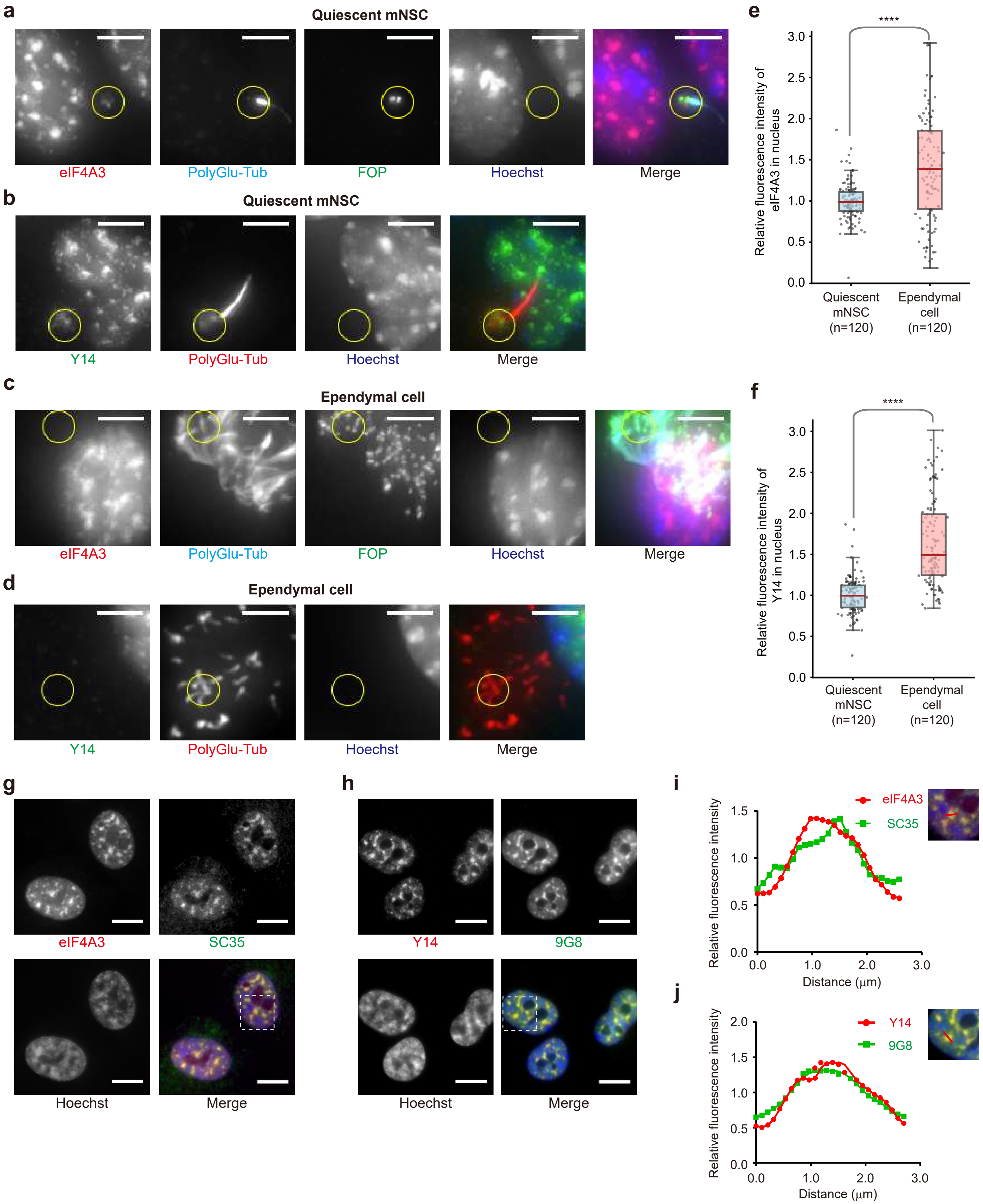

a
Quiescent mNSC
eIF4A3
PolyGlu-Tub
FOP
Hoechst
Merge
b
Quiescent mNSC
Y14
PolyGlu-Tub
Hoechst
Merge
c
Ependymal cell
eIF4A3
PolyGlu-Tub
FOP
Hoechst
Merge
d
Ependymal cell
Y14
PolyGlu-Tub
Hoechst
Merge
e
****
Relative fluorescence intensity of eIF4A3 in nucleus
0.0
0.5
1.0
1.5
2.0
2.5
3.0
Quiescent mNSC (n=120)
Ependymal cell (n=120)
f
****
Relative fluorescence intensity of Y14 in nucleus
Quiescent mNSC (n=120)
Ependymal cell (n=120)
g
eIF4A3
SC35
Hoechst
Merge
h
Y14
9G8
Hoechst
Merge
i
Relative fluorescence intensity
Distance (μm)
eIF4A3
SC35
j
Relative fluorescence intensity
Distance (μm)
Y14
9G8

Fig.S2

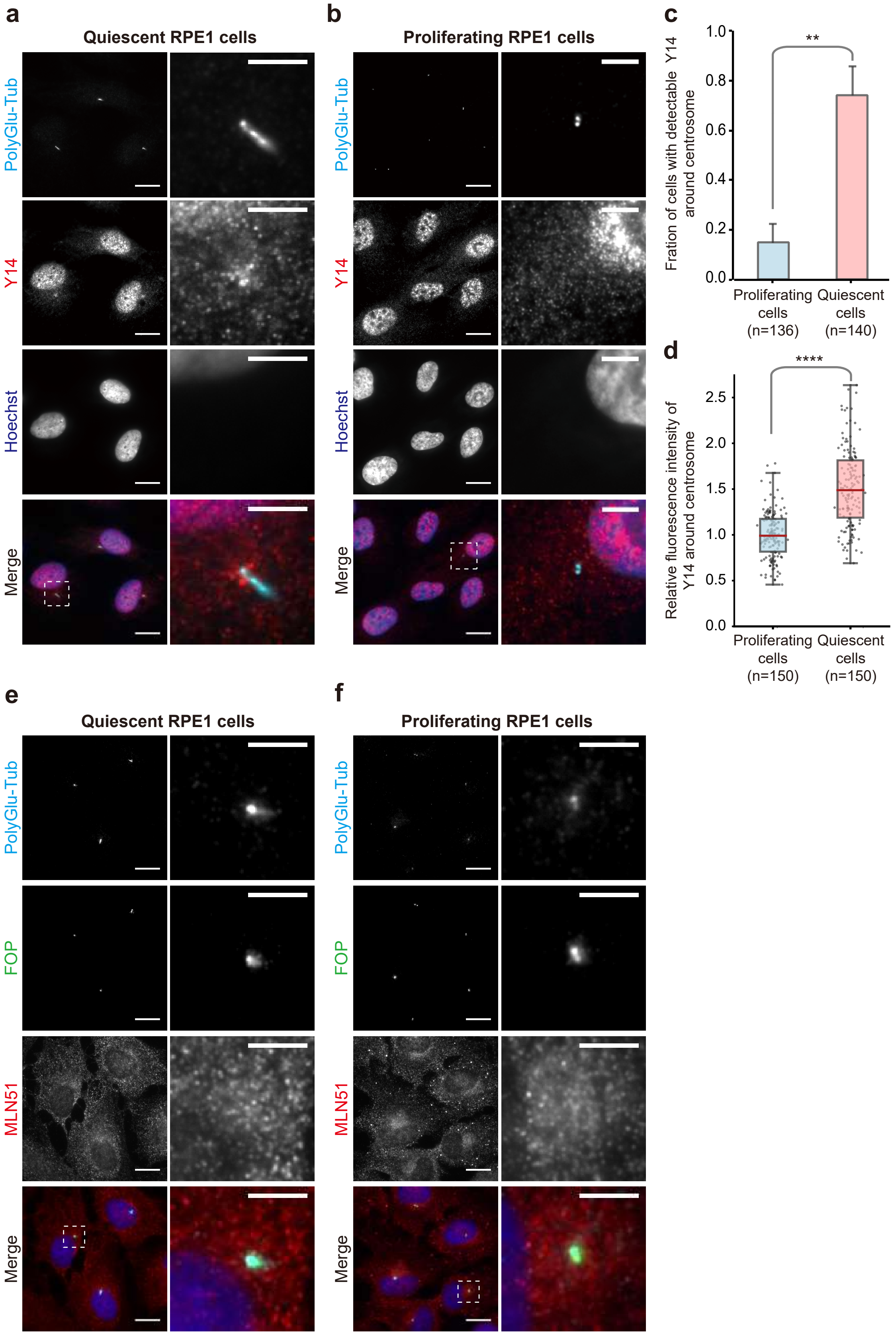

a
Quiescent RPE1 cells
PolyGlu-Tub
Y14
Hoechst
Merge
b
Proliferating RPE1 cells
PolyGlu-Tub
Y14
Hoechst
Merge
c
Fration of cells with detectable Y14 around centrosome
1.0
0.8
0.6
0.4
0.2
0.0
**
Proliferating cells (n=136)
Quiescent cells (n=140)
d
Relative fluorescence intensity of Y14 around centrosome
2.5
2.0
1.5
1.0
0.5
0.0
****
Proliferating cells (n=150)
Quiescent cells (n=150)
e
Quiescent RPE1 cells
PolyGlu-Tub
FOP
MLN51
Merge
f
Proliferating RPE1 cells
PolyGlu-Tub
FOP
MLN51
Merge

Fig.S3

a

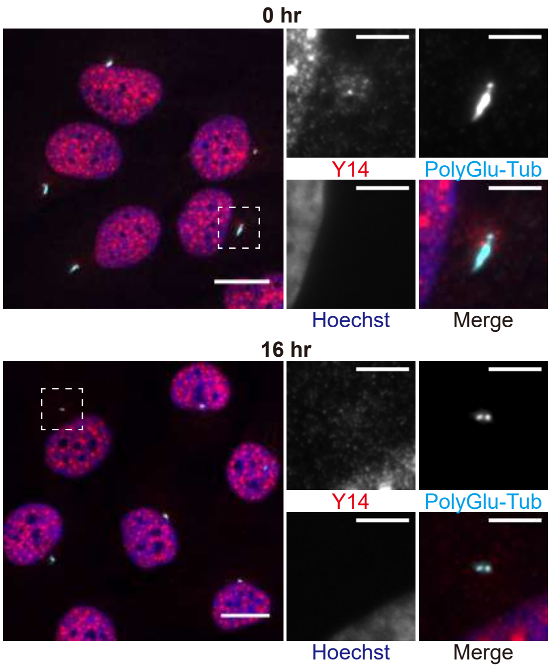


b

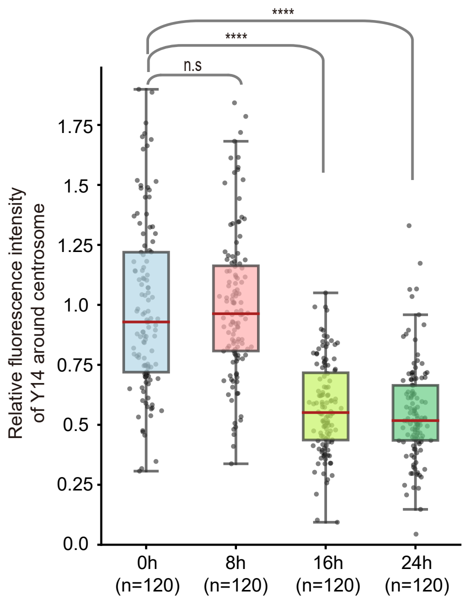


c

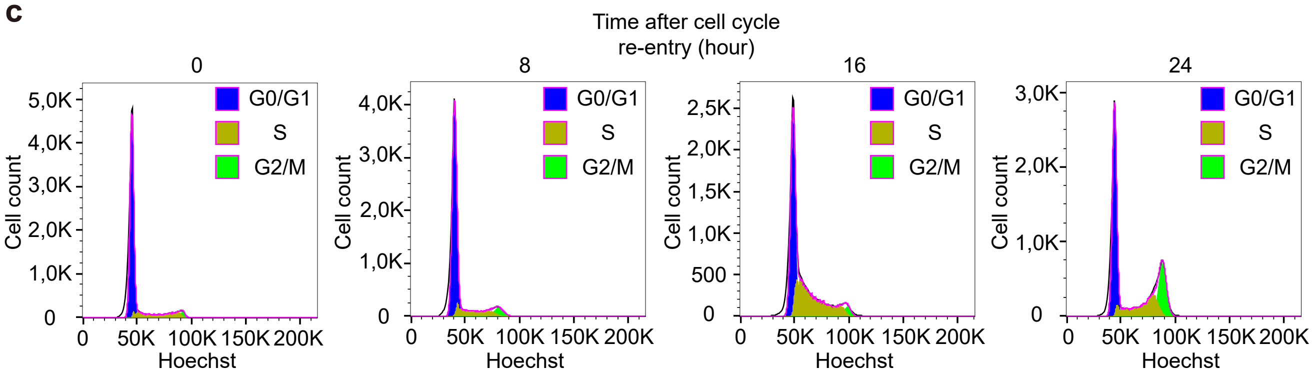


d

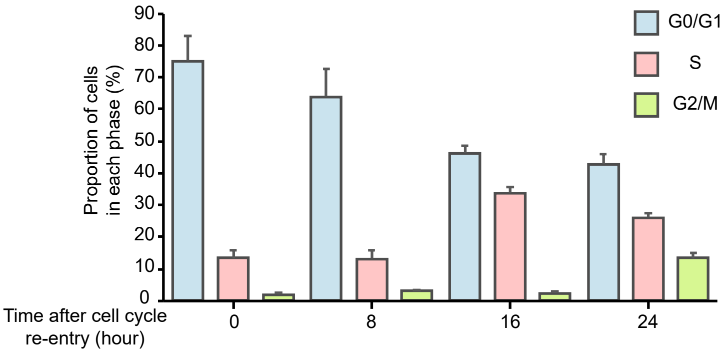


e

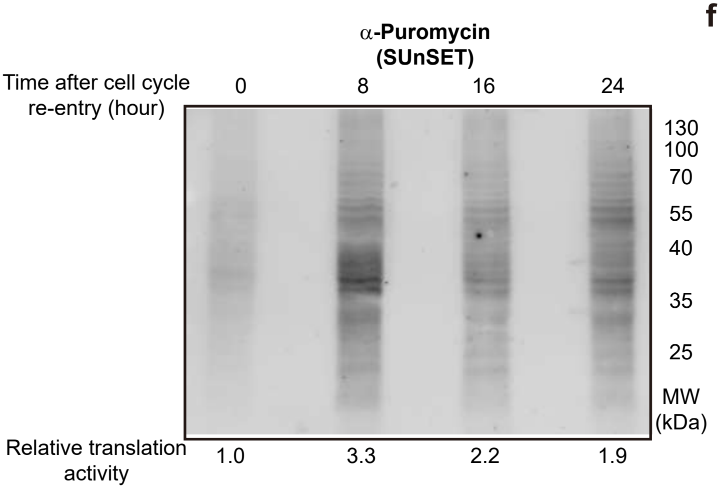


f

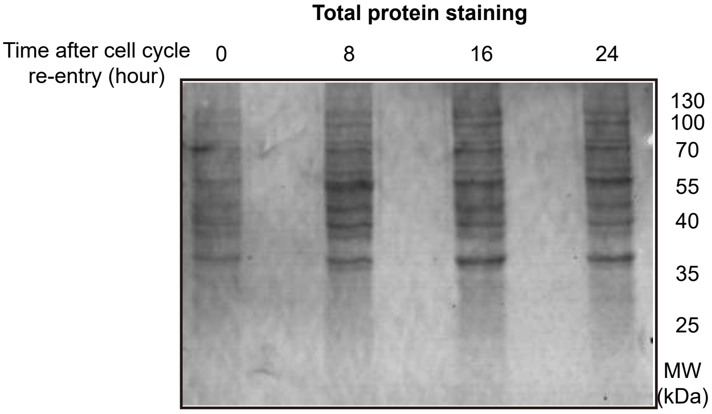

Fig.S4

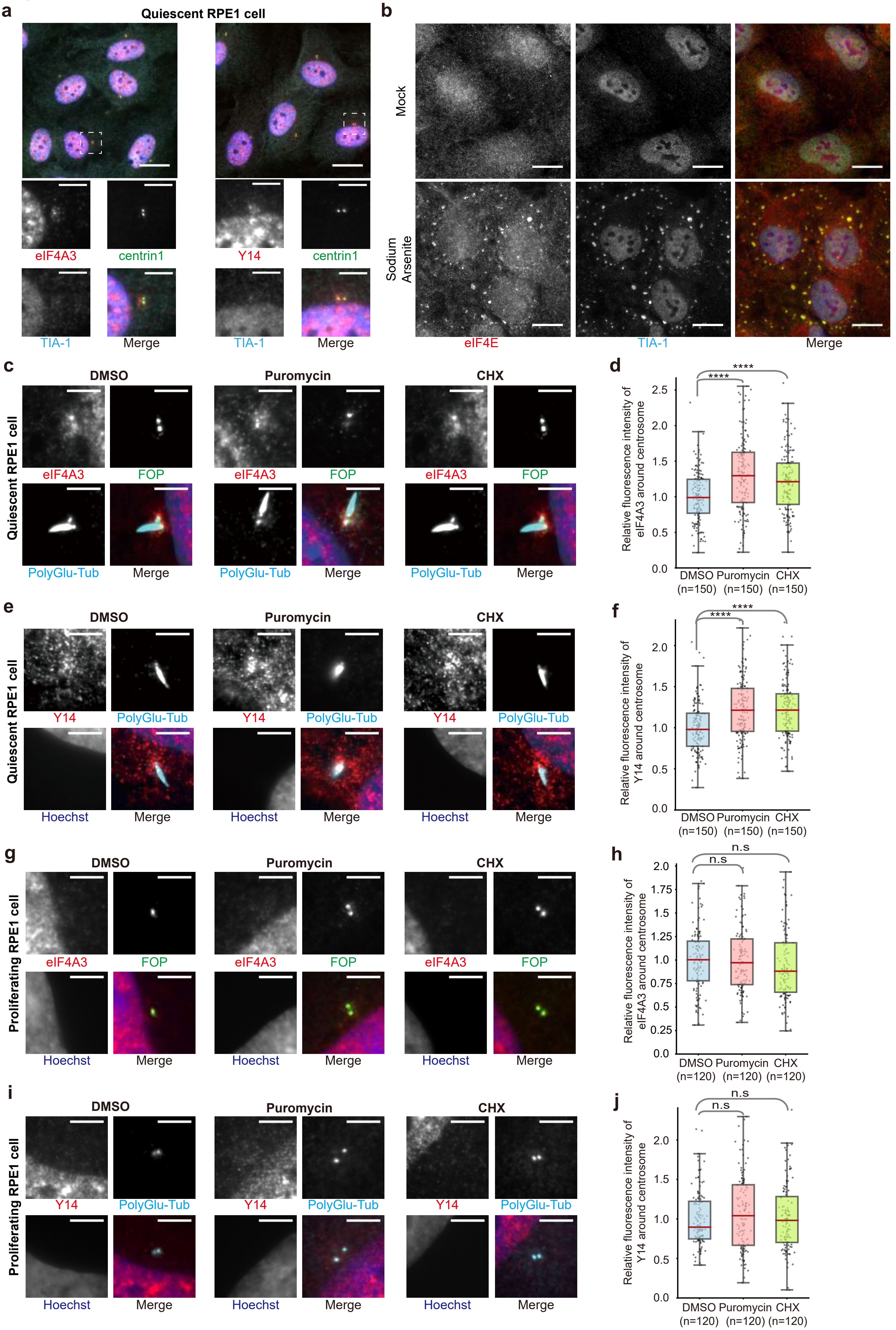
a
Quiescent RPE1 cell
eIF4A3
centrin1
Y14
centrin1
TIA-1
Merge
TIA-1
Merge
b
Mock
Sodium Arsenite
eIF4E
TIA-1
Merge
c
DMSO
Puromycin
CHX
Quiescent RPE1 cell
eIF4A3
FOP
PolyGlu-Tub
Merge
d
Relative fluorescence intensity of eIF4A3 around centrosome
****
****
DMSO (n=150)
Puromycin (n=150)
CHX (n=150)
e
DMSO
Puromycin
CHX
Quiescent RPE1 cell
Y14
PolyGlu-Tub
Hoechst
Merge
f
Relative fluorescence intensity of Y14 around centrosome
****
****
DMSO (n=150)
Puromycin (n=150)
CHX (n=150)
g
DMSO
Puromycin
CHX
Proliferating RPE1 cell
eIF4A3
FOP
Hoechst
Merge
h
Relative fluorescence intensity of eIF4A3 around centrosome
n.s
n.s
DMSO (n=120)
Puromycin (n=120)
CHX (n=120)
i
DMSO
Puromycin
CHX
Proliferating RPE1 cell
Y14
PolyGlu-Tub
Hoechst
Merge
j
Relative fluorescence intensity of Y14 around centrosome
n.s
n.s
DMSO (n=120)
Puromycin (n=120)
CHX (n=120)

Fig.S5

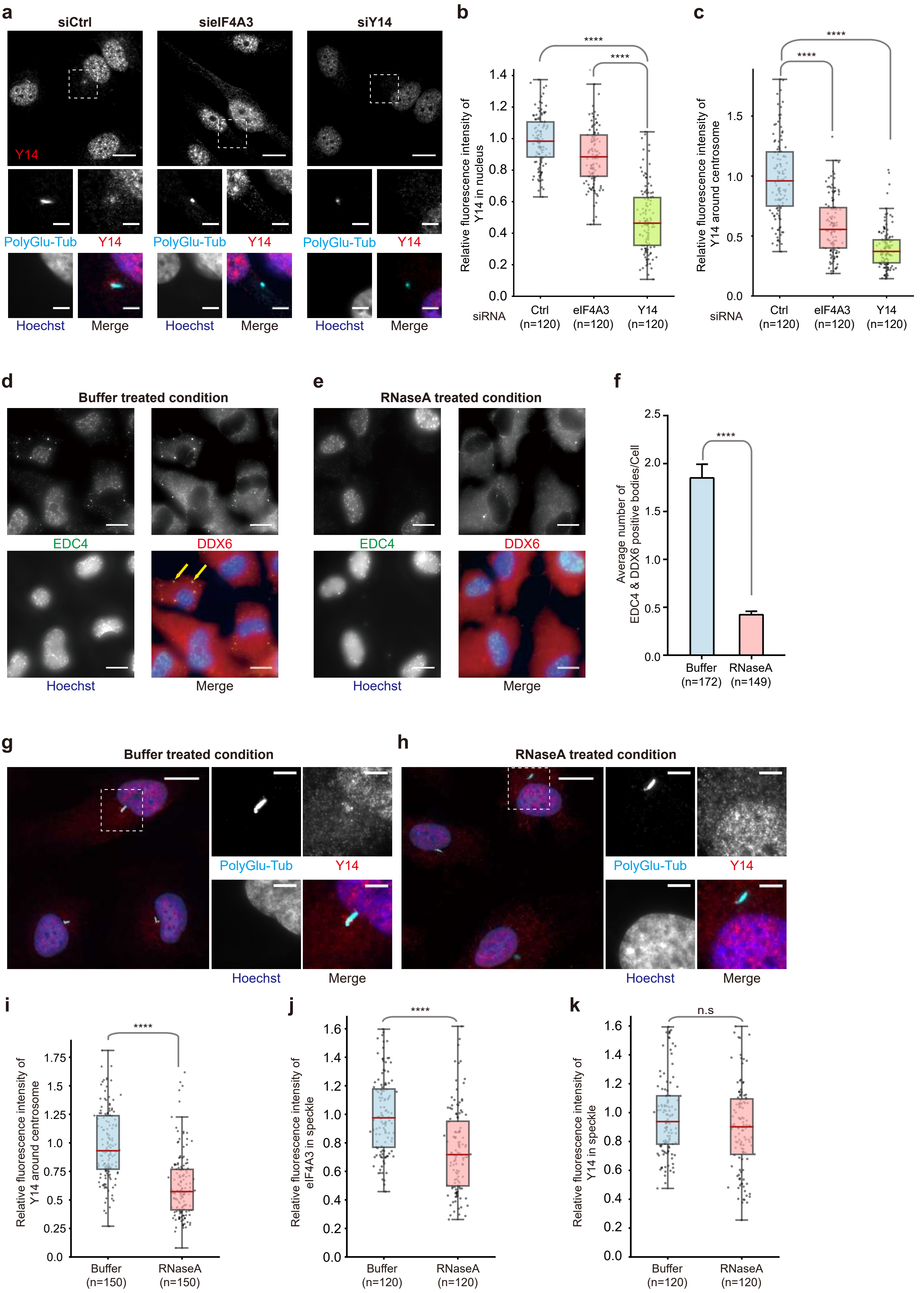
a
siCtrl
sieIF4A3
siY14
Y14
PolyGlu-Tub
Y14
Hoechst
Merge
b
Relative fluorescence intensity of Y14 in nucleus
****
****
siRNA
Ctrl (n=120)
eIF4A3 (n=120)
Y14 (n=120)
c
Relative fluorescence intensity of Y14 around centrosome
****
****
siRNA
Ctrl (n=120)
eIF4A3 (n=120)
Y14 (n=120)
d
Buffer treated condition
EDC4
DDX6
Hoechst
Merge
e
RNaseA treated condition
EDC4
DDX6
Hoechst
Merge
f
Average number of EDC4 & DDX6 positive bodies/Cell
****
Buffer (n=172)
RNaseA (n=149)
g
Buffer treated condition
PolyGlu-Tub
Y14
Hoechst
Merge
h
RNaseA treated condition
PolyGlu-Tub
Y14
Hoechst
Merge
i
Relative fluorescence intensity of Y14 around centrosome
****
Buffer (n=150)
RNaseA (n=150)
j
Relative fluorescence intensity of eIF4A3 in speckle
****
Buffer (n=120)
RNaseA (n=120)
k
Relative fluorescence intensity of Y14 in speckle
n.s
Buffer (n=120)
RNaseA (n=120)

Fig.S6

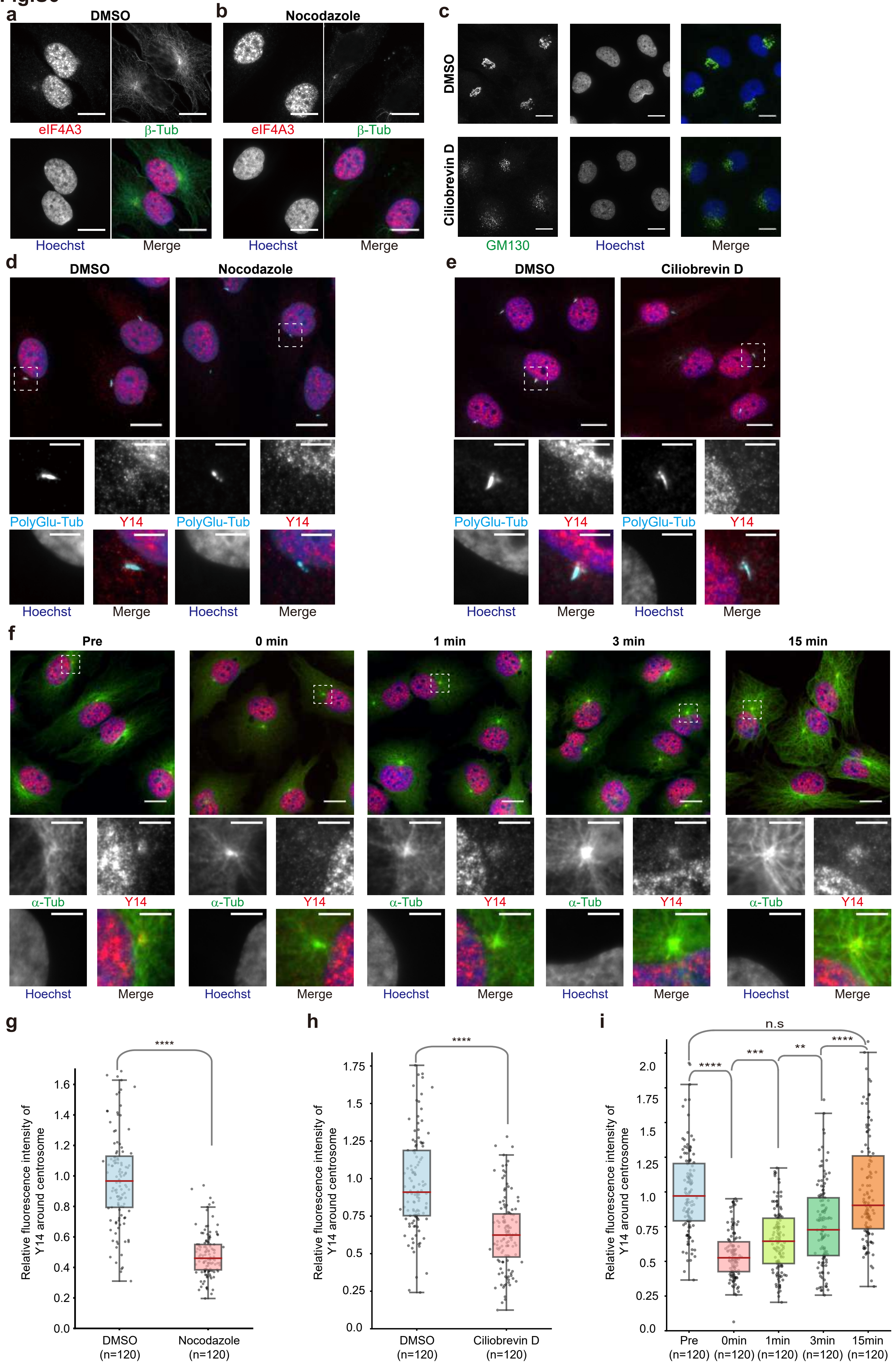

a
DMSO
eIF4A3
β-Tub
Hoechst
Merge
b
Nocodazole
eIF4A3
β-Tub
Hoechst
Merge
c
DMSO
Ciliobrevin D
GM130
Hoechst
Merge
d
DMSO
Nocodazole
PolyGlu-Tub
Y14
Hoechst
Merge
e
DMSO
Ciliobrevin D
PolyGlu-Tub
Y14
Hoechst
Merge
f
Pre
0 min
1 min
3 min
15 min
α-Tub
Y14
Hoechst
Merge
g
Relative fluorescence intensity of Y14 around centrosome
****
DMSO (n=120)
Nocodazole (n=120)
h
Relative fluorescence intensity of Y14 around centrosome
****
DMSO (n=120)
Ciliobrevin D (n=120)
i
Relative fluorescence intensity of Y14 around centrosome
n.s
****
***
**
****
Pre (n=120)
0min (n=120)
1min (n=120)
3min (n=120)
15min (n=120)

Fig.S7

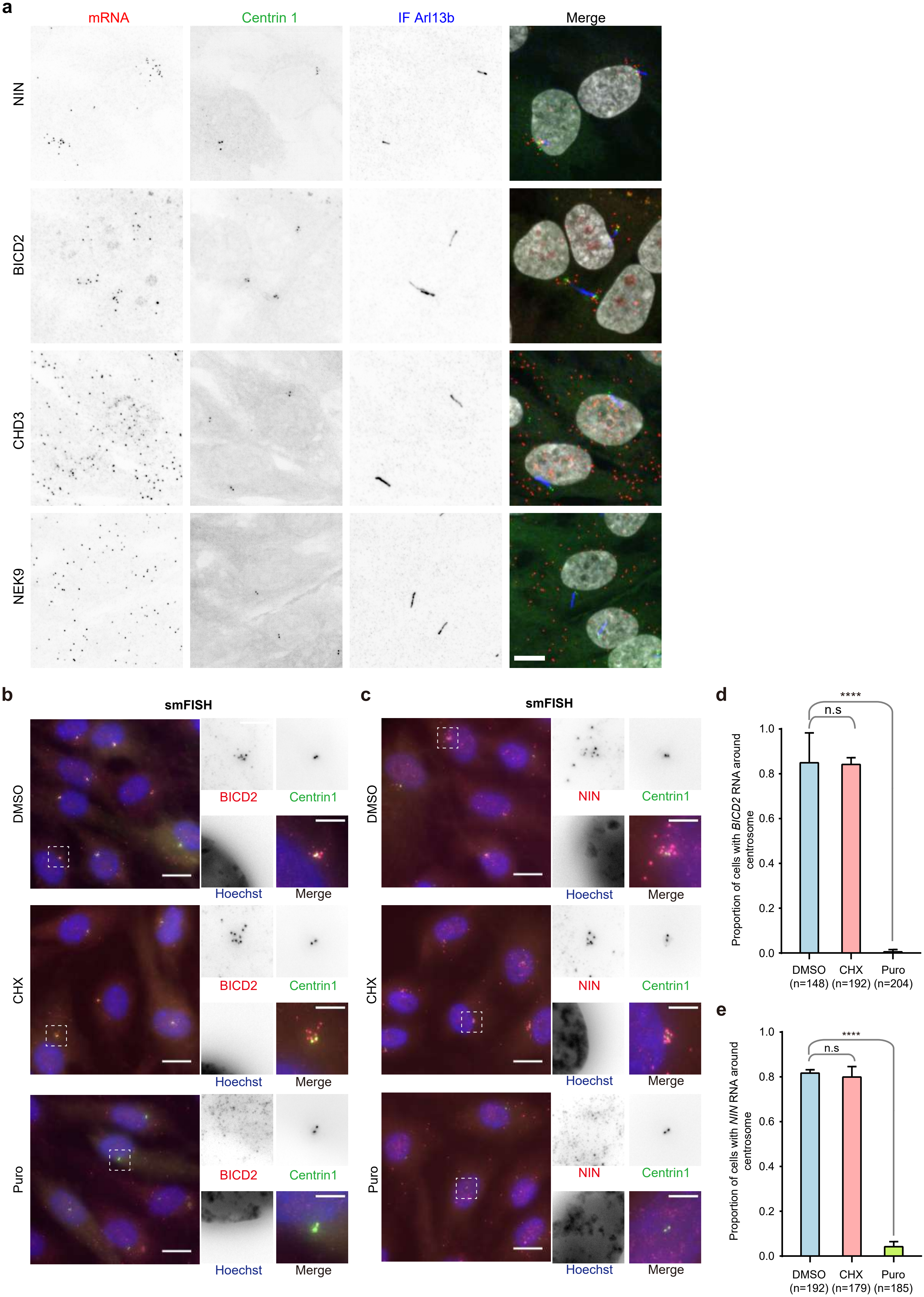

a
mRNA
Centrin 1
IF Arl13b
Merge
NIN
BICD2
CHD3
NEK9
b
smFISH
DMSO
CHX
Puro
BICD2
Centrin1
Hoechst
Merge
c
smFISH
NIN
d
Proportion of cells with BICD2 RNA around centrosome
n.s
****
1.0
0.8
0.6
0.4
0.2
0.0
DMSO (n=148)
CHX (n=192)
Puro (n=204)
e
Proportion of cells with NIN RNA around centrosome
DMSO (n=192)
CHX (n=179)
Puro (n=185)

Fig.S8

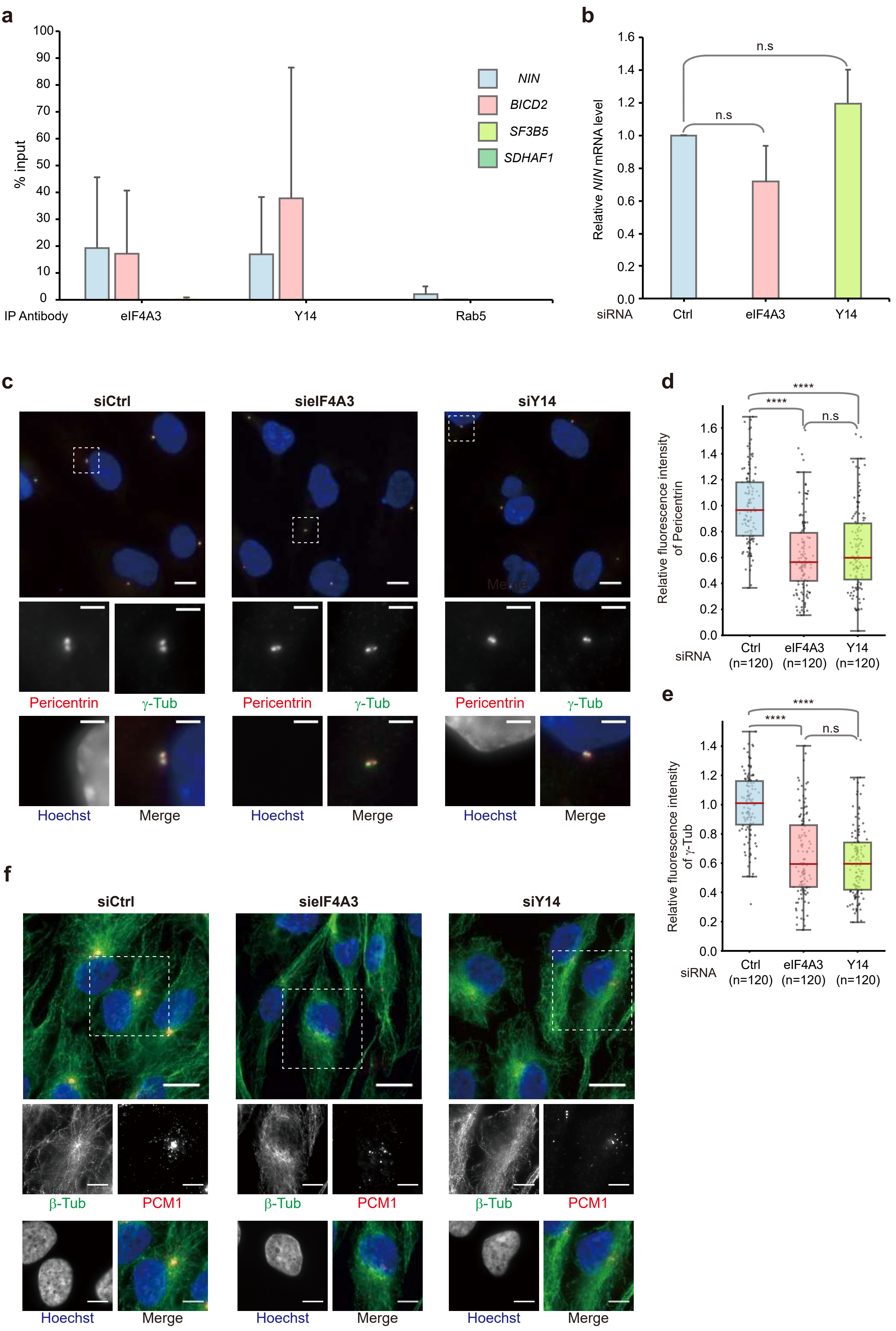
a
% input
IP Antibody
eIF4A3
Y14
Rab5
NIN
BICD2
SF3B5
SDHAF1
b
Relative NIN mRNA level
siRNA
Ctrl
eIF4A3
Y14
n.s
n.s
c
siCtrl
sieIF4A3
siY14
Pericentrin
γ-Tub
Hoechst
Merge
d
Relative fluorescence intensity of Pericentrin
siRNA
Ctrl (n=120)
eIF4A3 (n=120)
Y14 (n=120)
****
****
n.s
e
Relative fluorescence intensity of γ-Tub
siRNA
Ctrl (n=120)
eIF4A3 (n=120)
Y14 (n=120)
****
****
n.s
f
siCtrl
sieIF4A3
siY14
β-Tub
PCM1
Hoechst
Merge

Fig.S9

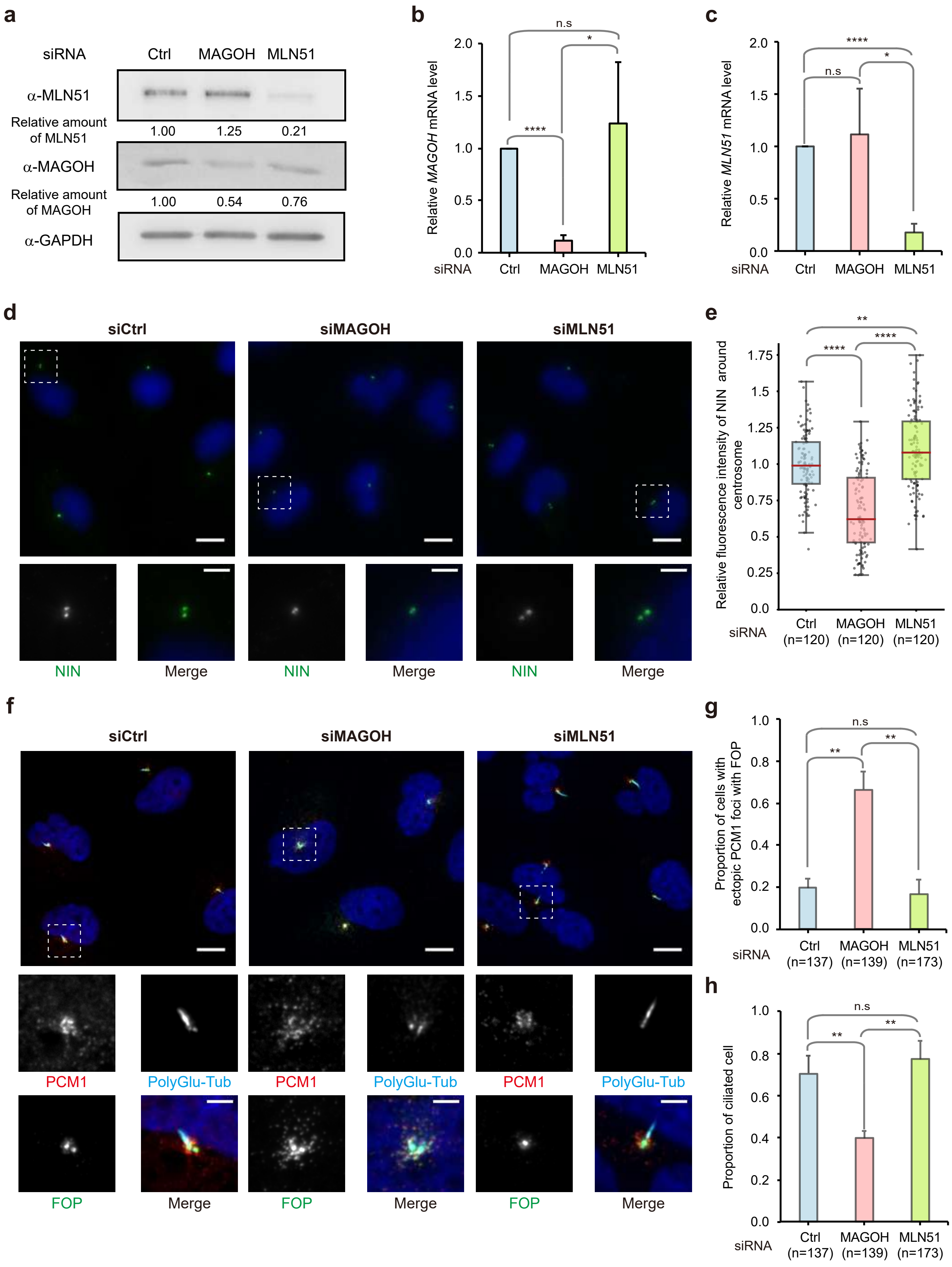
a
siRNA
Ctrl
MAGOH
MLN51
α-MLN51
Relative amount of MLN51
1.00
1.25
0.21
α-MAGOH
Relative amount of MAGOH
1.00
0.54
0.76
α-GAPDH
b
Relative MAGOH mRNA level
n.s
*
****
c
Relative MLN51 mRNA level
d
siCtrl
siMAGOH
siMLN51
NIN
Merge
e
Relative fluorescence intensity of NIN around centrosome
**
****
Ctrl (n=120)
MAGOH (n=120)
MLN51 (n=120)
f
PCM1
PolyGlu-Tub
FOP
g
Proportion of cells with ectopic PCM1 foci with FOP
Ctrl (n=137)
MAGOH (n=139)
MLN51 (n=173)
h
Proportion of ciliated cell

**Supplementary table S1**

| ENSG | ENST | GeneName | Cell line | **Localization** |
|---|---|---|---|---|
| ENSG00000185963 | ENST00000375512 | BICD2 | Diff. RPE1 GFP-Centrin | Cilium base |
| ENSG00000100503 | ENST00000389868 | NIN | Diff. RPE1 GFP-Centrin | Cilium base |
| ENSG00000122545 | ENST00000399035 | SEPT7 | Diff. RPE1 GFP-Centrin | Clusters |
| ENSG00000138160 | ENST00000260731 | KIF11 | Diff. RPE1 GFP-Centrin | Clusters |
| ENSG00000197102 | ENST00000555062 | DYNC1H1 | Diff. RPE1 GFP-Centrin | Foci |
| ENSG00000107581 | ENST00000369144 | EIF3A | Diff. RPE1 GFP-Centrin | Foci |
| ENSG00000163346 | ENST00000368463 | PBXIP1 | Diff. RPE1 GFP-Centrin | Foci |
| ENSG00000273217 | ENST00000514667 | AC008695.1 | Diff. RPE1 GFP-Centrin | no signal |
| ENSG00000115073 | ENST00000289228 | ACTR1B | Diff. RPE1 GFP-Centrin | no signal |
| ENSG00000165923 | ENST00000525123 | AGBL2 | Diff. RPE1 GFP-Centrin | no signal |
| ENSG00000186094 | ENST00000371839 | AGBL4 | Diff. RPE1 GFP-Centrin | no signal |
| ENSG00000169126 | ENST00000305242 | ARMC4 | Diff. RPE1 GFP-Centrin | no signal |
| ENSG00000039987 | ENST00000042931 | BEST2 | Diff. RPE1 GFP-Centrin | no signal |
| ENSG00000135127 | ENST00000397558 | BICDL1 | Diff. RPE1 GFP-Centrin | no signal |
| ENSG00000160469 | ENST00000309383 | BRSK1 | Diff. RPE1 GFP-Centrin | no signal |
| ENSG00000154493 | ENST00000284694 | C10ORF90 | Diff. RPE1 GFP-Centrin | no signal |
| ENSG00000205129 | ENST00000378850 | C4ORF47 | Diff. RPE1 GFP-Centrin | no signal |
| ENSG00000181751 | ENST00000510890 | C5ORF30 | Diff. RPE1 GFP-Centrin | no signal |
| ENSG00000004948 | ENST00000360249 | CALCR | Diff. RPE1 GFP-Centrin | no signal |
| ENSG00000076826 | ENST00000446248 | CAMSAP3 | Diff. RPE1 GFP-Centrin | no signal |
| ENSG00000133962 | ENST00000256343 | CATSPERB | Diff. RPE1 GFP-Centrin | no signal |
| ENSG00000174898 | ENST00000381624 | CATSPERD | Diff. RPE1 GFP-Centrin | no signal |
| ENSG00000161180 | ENST00000292779 | CCDC116 | Diff. RPE1 GFP-Centrin | no signal |
| ENSG00000244607 | ENST00000310232 | CCDC13 | Diff. RPE1 GFP-Centrin | no signal |
| ENSG00000135205 | ENST00000285871 | CCDC146 | Diff. RPE1 GFP-Centrin | no signal |
| ENSG00000182645 | ENST00000333254 | CCDC172 | Diff. RPE1 GFP-Centrin | no signal |
| ENSG00000160050 | ENST00000373602 | CCDC28B | Diff. RPE1 GFP-Centrin | no signal |
| ENSG00000145075 | ENST00000273654 | CCDC39 | Diff. RPE1 GFP-Centrin | no signal |
| ENSG00000166510 | ENST00000591504 | CCDC68 | Diff. RPE1 GFP-Centrin | no signal |
| ENSG00000015133 | ENST00000389857 | CCDC88C | Diff. RPE1 GFP-Centrin | no signal |
| ENSG00000079335 | ENST00000361544 | CDC14A | Diff. RPE1 GFP-Centrin | no signal |
| ENSG00000170312 | ENST00000395284 | CDK1 | Diff. RPE1 GFP-Centrin | no signal |
| ENSG00000100629 | ENST00000281129 | CEP128 | Diff. RPE1 GFP-Centrin | no signal |
| ENSG00000154608 | ENST00000502249 | CEP170P1 | Diff. RPE1 GFP-Centrin | no signal |
| ENSG00000112877 | ENST00000264935 | CEP72 | Diff. RPE1 GFP-Centrin | no signal |
| ENSG00000101624 | ENST00000262127 | CEP76 | Diff. RPE1 GFP-Centrin | no signal |
| ENSG00000111860 | ENST00000368488 | CEP85L | Diff. RPE1 GFP-Centrin | no signal |
| ENSG00000121289 | ENST00000305768 | CEP89 | Diff. RPE1 GFP-Centrin | no signal |
| ENSG00000163885 | ENST00000352312 | CFAP100 | Diff. RPE1 GFP-Centrin | no signal |
| ENSG00000188931 | ENST00000367974 | CFAP126 | Diff. RPE1 GFP-Centrin | no signal |
| ENSG00000160401 | ENST00000373295 | CFAP157 | Diff. RPE1 GFP-Centrin | no signal |
| ENSG00000070761 | ENST00000262498 | CFAP20 | Diff. RPE1 GFP-Centrin | no signal |
| ENSG00000272514 | ENST00000369562 | CFAP206 | Diff. RPE1 GFP-Centrin | no signal |
| ENSG00000163075 | ENST00000413057 | CFAP221 | Diff. RPE1 GFP-Centrin | no signal |
| ENSG00000213085 | ENST00000368099 | CFAP45 | Diff. RPE1 GFP-Centrin | no signal |
| ENSG00000171811 | ENST00000368586 | CFAP46 | Diff. RPE1 GFP-Centrin | no signal |
| ENSG00000172361 | ENST00000398545 | CFAP53 | Diff. RPE1 GFP-Centrin | no signal |
| ENSG00000156042 | ENST00000310715 | CFAP70 | Diff. RPE1 GFP-Centrin | no signal |
| ENSG00000186710 | ENST00000335621 | CFAP73 | Diff. RPE1 GFP-Centrin | no signal |
| ENSG00000169607 | ENST00000541405 | CKAP2L | Diff. RPE1 GFP-Centrin | no signal |

| | | | | |
|---|---|---|---|---|
| ENSG00000126890 | ENST00000247306 | CTAG2 | Diff. RPE1 GFP-Centrin | no signal |
| ENSG00000165325 | ENST00000298050 | DEUP1 | Diff. RPE1 GFP-Centrin | no signal |
| ENSG00000166938 | ENST00000319194 | DIS3L | Diff. RPE1 GFP-Centrin | no signal |
| ENSG00000166938 | ENST00000319194 | DIS3L | Diff. RPE1 GFP-Centrin | no signal |
| ENSG00000105877 | ENST00000409508 | DNAH11 | Diff. RPE1 GFP-Centrin | no signal |
| ENSG00000174844 | ENST00000311202 | DNAH12 | Diff. RPE1 GFP-Centrin | no signal |
| ENSG00000158486 | ENST00000261383 | DNAH3 | Diff. RPE1 GFP-Centrin | no signal |
| ENSG00000039139 | ENST00000265104 | DNAH5 | Diff. RPE1 GFP-Centrin | no signal |
| ENSG00000115423 | ENST00000389394 | DNAH6 | Diff. RPE1 GFP-Centrin | no signal |
| ENSG00000118997 | ENST00000312428 | DNAH7 | Diff. RPE1 GFP-Centrin | no signal |
| ENSG00000124721 | ENST00000327475 | DNAH8 | Diff. RPE1 GFP-Centrin | no signal |
| ENSG00000197959 | ENST00000367731 | DNM3 | Diff. RPE1 GFP-Centrin | no signal |
| ENSG00000144635 | ENST00000273130 | DYNC1LI1 | Diff. RPE1 GFP-Centrin | no signal |
| ENSG00000096093 | ENST00000371068 | EFHC1 | Diff. RPE1 GFP-Centrin | no signal |
| ENSG00000013016 | ENST00000322054 | EHD3 | Diff. RPE1 GFP-Centrin | no signal |
| ENSG00000151023 | ENST00000376363 | ENKUR | Diff. RPE1 GFP-Centrin | no signal |
| ENSG00000165689 | ENST00000371725 | ENTR1 | Diff. RPE1 GFP-Centrin | no signal |
| ENSG00000173040 | ENST00000344408 | EVC2 | Diff. RPE1 GFP-Centrin | no signal |
| ENSG00000182263 | ENST00000333129 | FIGN | Diff. RPE1 GFP-Centrin | no signal |
| ENSG00000189139 | ENST00000340446 | FSCB | Diff. RPE1 GFP-Centrin | no signal |
| ENSG00000065135 | ENST00000369851 | GNAI3 | Diff. RPE1 GFP-Centrin | no signal |
| ENSG00000215203 | ENST00000399770 | GRXCR1 | Diff. RPE1 GFP-Centrin | no signal |
| ENSG00000177602 | ENST00000325418 | GSG2 | Diff. RPE1 GFP-Centrin | no signal |
| ENSG00000092036 | ENST00000342454 | HAUS4 | Diff. RPE1 GFP-Centrin | no signal |
| ENSG00000131351 | ENST00000253669 | HAUS8 | Diff. RPE1 GFP-Centrin | no signal |
| ENSG00000131351 | ENST00000253669 | HAUS8 | Diff. RPE1 GFP-Centrin | no signal |
| ENSG00000188175 | ENST00000394468 | HEPACAM2 | Diff. RPE1 GFP-Centrin | no signal |
| ENSG00000188175 | ENST00000341723 | HEPACAM2 | Diff. RPE1 GFP-Centrin | no signal |
| ENSG00000134709 | ENST00000371208 | HOOK1 | Diff. RPE1 GFP-Centrin | no signal |
| ENSG00000158748 | ENST00000289753 | HTR6 | Diff. RPE1 GFP-Centrin | no signal |
| ENSG00000157423 | ENST00000393567 | HYDIN | Diff. RPE1 GFP-Centrin | no signal |
| ENSG00000130294 | ENST00000648680 | KIF1A | Diff. RPE1 GFP-Centrin | no signal |
| ENSG00000125337 | ENST00000351261 | KIF25 | Diff. RPE1 GFP-Centrin | no signal |
| ENSG00000182866 | ENST00000336890 | LCK | Diff. RPE1 GFP-Centrin | no signal |
| ENSG00000172264 | ENST00000402914 | MACROD2 | Diff. RPE1 GFP-Centrin | no signal |
| ENSG00000111837 | ENST00000313243 | MAK | Diff. RPE1 GFP-Centrin | no signal |
| ENSG00000111837 | ENST00000313243 | MAK | Diff. RPE1 GFP-Centrin | no signal |
| ENSG00000212916 | ENST00000418460 | MAP10 | Diff. RPE1 GFP-Centrin | no signal |
| ENSG00000212916 | ENST00000418460 | MAP10 | Diff. RPE1 GFP-Centrin | no signal |
| ENSG00000173327 | ENST00000309100 | MAP3K11 | Diff. RPE1 GFP-Centrin | no signal |
| ENSG00000180834 | ENST00000318631 | MAP6D1 | Diff. RPE1 GFP-Centrin | no signal |
| ENSG00000163875 | ENST00000373075 | MEAF6 | Diff. RPE1 GFP-Centrin | no signal |
| ENSG00000163875 | ENST00000373075 | MEAF6 | Diff. RPE1 GFP-Centrin | no signal |
| ENSG00000243156 | ENST00000441493 | MICAL3 | Diff. RPE1 GFP-Centrin | no signal |
| ENSG00000158411 | ENST00000289359 | MITD1 | Diff. RPE1 GFP-Centrin | no signal |
| ENSG00000034971 | ENST00000037502 | MYOC | Diff. RPE1 GFP-Centrin | no signal |
| ENSG00000166579 | ENST00000402554 | NDEL1 | Diff. RPE1 GFP-Centrin | no signal |
| ENSG00000136098 | ENST00000610828 | NEK3 | Diff. RPE1 GFP-Centrin | no signal |
| ENSG00000101004 | ENST00000278886 | NINL | Diff. RPE1 GFP-Centrin | no signal |
| ENSG00000144061 | ENST00000445609 | NPHP1 | Diff. RPE1 GFP-Centrin | no signal |
| ENSG00000112530 | ENST00000366888 | PACRG | Diff. RPE1 GFP-Centrin | no signal |
| ENSG00000075891 | ENST00000355243 | PAX2 | Diff. RPE1 GFP-Centrin | no signal |
| ENSG00000075891 | ENST00000355243 | PAX2 | Diff. RPE1 GFP-Centrin | no signal |

| | | | | |
|---|---|---|---|---|
| ENSG00000116703 | ENST00000340129 | PDC | Diff. RPE1 GFP-Centrin | no signal |
| ENSG00000184588 | ENST00000371045 | PDE4B | Diff. RPE1 GFP-Centrin | no signal |
| ENSG00000158683 | ENST00000289672 | PKD1L1 | Diff. RPE1 GFP-Centrin | no signal |
| ENSG00000170927 | ENST00000340994 | PKHD1 | Diff. RPE1 GFP-Centrin | no signal |
| ENSG00000170927 | ENST00000371117 | PKHD1 | Diff. RPE1 GFP-Centrin | no signal |
| ENSG00000170927 | ENST00000371117 | PKHD1 | Diff. RPE1 GFP-Centrin | no signal |
| ENSG00000205038 | ENST00000378402 | PKHD1L1 | Diff. RPE1 GFP-Centrin | no signal |
| ENSG00000100078 | ENST00000215885 | PLA2G3 | Diff. RPE1 GFP-Centrin | no signal |
| ENSG00000178125 | ENST00000324682 | PPP1R42 | Diff. RPE1 GFP-Centrin | no signal |
| ENSG00000070950 | ENST00000264926 | RAD18 | Diff. RPE1 GFP-Centrin | no signal |
| ENSG00000164188 | ENST00000296604 | RANBP3L | Diff. RPE1 GFP-Centrin | no signal |
| ENSG00000165917 | ENST00000298854 | RAPSN | Diff. RPE1 GFP-Centrin | no signal |
| ENSG00000102760 | ENST00000379359 | RGCC | Diff. RPE1 GFP-Centrin | no signal |
| ENSG00000169220 | ENST00000408923 | RGS14 | Diff. RPE1 GFP-Centrin | no signal |
| ENSG00000104237 | ENST00000220676 | RP1 | Diff. RPE1 GFP-Centrin | no signal |
| ENSG00000102218 | ENST00000218340 | RP2 | Diff. RPE1 GFP-Centrin | no signal |
| ENSG00000092200 | ENST00000400017 | RPGRIP1 | Diff. RPE1 GFP-Centrin | no signal |
| ENSG00000025039 | ENST00000369415 | RRAGD | Diff. RPE1 GFP-Centrin | no signal |
| ENSG00000160188 | ENST00000291536 | RSPH1 | Diff. RPE1 GFP-Centrin | no signal |
| ENSG00000111834 | ENST00000229554 | RSPH4A | Diff. RPE1 GFP-Centrin | no signal |
| ENSG00000165480 | ENST00000462482 | SKA3 | Diff. RPE1 GFP-Centrin | no signal |
| ENSG00000188817 | ENST00000343837 | SNTN | Diff. RPE1 GFP-Centrin | no signal |
| ENSG00000086300 | ENST00000338523 | SNX10 | Diff. RPE1 GFP-Centrin | no signal |
| ENSG00000095637 | ENST00000306402 | SORBS1 | Diff. RPE1 GFP-Centrin | no signal |
| ENSG00000155761 | ENST00000336338 | SPAG17 | Diff. RPE1 GFP-Centrin | no signal |
| ENSG00000123473 | ENST00000371877 | STIL | Diff. RPE1 GFP-Centrin | no signal |
| ENSG00000165730 | ENST00000421961 | STOX1 | Diff. RPE1 GFP-Centrin | no signal |
| ENSG00000147642 | ENST00000276646 | SYBU | Diff. RPE1 GFP-Centrin | no signal |
| ENSG00000111490 | ENST00000229088 | TBC1D30 | Diff. RPE1 GFP-Centrin | no signal |
| ENSG00000184786 | ENST00000366774 | TCTE3 | Diff. RPE1 GFP-Centrin | no signal |
| ENSG00000167858 | ENST00000338694 | TEKT1 | Diff. RPE1 GFP-Centrin | no signal |
| ENSG00000092850 | ENST00000207457 | TEKT2 | Diff. RPE1 GFP-Centrin | no signal |
| ENSG00000125409 | ENST00000395930 | TEKT3 | Diff. RPE1 GFP-Centrin | no signal |
| ENSG00000153060 | ENST00000283025 | TEKT5 | Diff. RPE1 GFP-Centrin | no signal |
| ENSG00000187049 | ENST00000334888 | TMEM216 | Diff. RPE1 GFP-Centrin | no signal |
| ENSG00000138100 | ENST00000380075 | TRIM54 | Diff. RPE1 GFP-Centrin | no signal |
| ENSG00000111199 | ENST00000261740 | TRPV4 | Diff. RPE1 GFP-Centrin | no signal |
| ENSG00000126467 | ENST00000246801 | TSKS | Diff. RPE1 GFP-Centrin | no signal |
| ENSG00000131044 | ENST00000375921 | TTLL9 | Diff. RPE1 GFP-Centrin | no signal |
| ENSG00000178462 | ENST00000380419 | TUBAL3 | Diff. RPE1 GFP-Centrin | no signal |
| ENSG00000104804 | ENST00000221399 | TULP2 | Diff. RPE1 GFP-Centrin | no signal |
| ENSG00000162543 | ENST00000375099 | UBXN10 | Diff. RPE1 GFP-Centrin | no signal |
| ENSG00000075702 | ENST00000401500 | WDR62 | Diff. RPE1 GFP-Centrin | no signal |
| ENSG00000143156 | ENST00000367811 | NME7 | Diff. RPE1 GFP-Centrin | nuclear |
| ENSG00000151475 | ENST00000281154 | SLC25A31 | Diff. RPE1 GFP-Centrin | nuclear |
| ENSG00000214021 | ENST00000383827 | TTLL3 | Diff. RPE1 GFP-Centrin | nuclear |
| ENSG00000122257 | ENST00000319715 | RBBP6 | Diff. RPE1 GFP-Centrin | perinuclear |
| ENSG00000117394 | ENST00000426263 | SLC2A1 | Diff. RPE1 GFP-Centrin | Perinuclear |
| ENSG00000169504 | ENST00000374379 | CLIC4 | Diff. RPE1 GFP-Centrin | Polarized |
| ENSG00000141367 | ENST00000621829 | CLTC | Diff. RPE1 GFP-Centrin | Polarized |
| ENSG00000109861 | ENST00000227266 | CTSC | Diff. RPE1 GFP-Centrin | Polarized |
| ENSG00000044574 | ENST00000324460 | HSPA5 | Diff. RPE1 GFP-Centrin | Polarized |
| ENSG00000111057 | ENST00000388835 | KRT18 | Diff. RPE1 GFP-Centrin | Polarized |

| | | | | |
|---|---|---|---|---|
| ENSG00000168036 | ENST00000349496 | CTNNB1 | Diff. RPE1 GFP-Centrin | Polarized, Foci |
| ENSG00000170004 | ENST00000330494 | CHD3 | Diff. RPE1 GFP-Centrin | Protrusion |
| ENSG00000129250 | ENST00000320785 | KIF1C | Diff. RPE1 GFP-Centrin | Protrusion |
| ENSG00000170759 | ENST00000302418 | KIF5B | Diff. RPE1 GFP-Centrin | Protrusion |
| ENSG00000094914 | ENST00000209873 | AAAS | Diff. RPE1 GFP-Centrin | Random |
| ENSG00000275700 | ENST00000619387 | AATF | Diff. RPE1 GFP-Centrin | Random |
| ENSG00000165660 | ENST00000298492 | ABRAXAS2 | Diff. RPE1 GFP-Centrin | Random |
| ENSG00000165660 | ENST00000298492 | ABRAXAS2 | Diff. RPE1 GFP-Centrin | Random |
| ENSG00000113812 | ENST00000335754 | ACTR8 | Diff. RPE1 GFP-Centrin | Random |
| ENSG00000113812 | ENST00000335754 | ACTR8 | Diff. RPE1 GFP-Centrin | Random |
| ENSG00000084693 | ENST00000323064 | AGBL5 | Diff. RPE1 GFP-Centrin | Random |
| ENSG00000135541 | ENST00000367800 | AHI1 | Diff. RPE1 GFP-Centrin | Random |
| ENSG00000105127 | ENST00000269701 | AKAP8 | Diff. RPE1 GFP-Centrin | Random |
| ENSG00000106948 | ENST00000374088 | AKNA | Diff. RPE1 GFP-Centrin | Random |
| ENSG00000106524 | ENST00000306999 | ANKMY2 | Diff. RPE1 GFP-Centrin | Random |
| ENSG00000107890 | ENST00000376070 | ANKRD26 | Diff. RPE1 GFP-Centrin | Random |
| ENSG00000165138 | ENST00000353234 | ANKS6 | Diff. RPE1 GFP-Centrin | Random |
| ENSG00000135046 | ENST00000257497 | ANXA1 | Diff. RPE1 GFP-Centrin | Random |
| ENSG00000122359 | ENST00000422982 | ANXA11 | Diff. RPE1 GFP-Centrin | Random |
| ENSG00000182718 | ENST00000451270 | ANXA2 | Diff. RPE1 GFP-Centrin | Random |
| ENSG00000134982 | ENST00000257430 | APC | Diff. RPE1 GFP-Centrin | Random |
| ENSG00000134982 | ENST00000257430 | APC | Diff. RPE1 GFP-Centrin | Random |
| ENSG00000100823 | ENST00000216714 | APEX1 | Diff. RPE1 GFP-Centrin | Random |
| ENSG00000062725 | ENST00000083182 | APPBP2 | Diff. RPE1 GFP-Centrin | Random |
| ENSG00000104728 | ENST00000518288 | ARHGEF10 | Diff. RPE1 GFP-Centrin | Random |
| ENSG00000169379 | ENST00000335438 | ARL13B | Diff. RPE1 GFP-Centrin | Random |
| ENSG00000213465 | ENST00000246747 | ARL2 | Diff. RPE1 GFP-Centrin | Random |
| ENSG00000102931 | ENST00000219204 | ARL2BP | Diff. RPE1 GFP-Centrin | Random |
| ENSG00000138175 | ENST00000260746 | ARL3 | Diff. RPE1 GFP-Centrin | Random |
| ENSG00000113966 | ENST00000463745 | ARL6 | Diff. RPE1 GFP-Centrin | Random |
| ENSG00000143862 | ENST00000272217 | ARL8A | Diff. RPE1 GFP-Centrin | Random |
| ENSG00000128272 | ENST00000396680 | ATF4 | Diff. RPE1 GFP-Centrin | Random |
| ENSG00000169136 | ENST00000423777 | ATF5 | Diff. RPE1 GFP-Centrin | Random |
| ENSG00000169136 | ENST00000423777 | ATF5 | Diff. RPE1 GFP-Centrin | Random |
| ENSG00000159720 | ENST00000290949 | ATP6V0D1 | Diff. RPE1 GFP-Centrin | Random |
| ENSG00000100554 | ENST00000216442 | ATP6V1D | Diff. RPE1 GFP-Centrin | Random |
| ENSG00000127423 | ENST00000374298 | AUNIP | Diff. RPE1 GFP-Centrin | Random |
| ENSG00000178999 | ENST00000534871 | AURKB | Diff. RPE1 GFP-Centrin | Random |
| ENSG00000168646 | ENST00000307078 | AXIN2 | Diff. RPE1 GFP-Centrin | Random |
| ENSG00000108641 | ENST00000261499 | B9D1 | Diff. RPE1 GFP-Centrin | Random |
| ENSG00000123810 | ENST00000243578 | B9D2 | Diff. RPE1 GFP-Centrin | Random |
| ENSG00000175866 | ENST00000428708 | BAIAP2 | Diff. RPE1 GFP-Centrin | Random |
| ENSG00000009954 | ENST00000339594 | BAZ1B | Diff. RPE1 GFP-Centrin | Random |
| ENSG00000119636 | ENST00000394009 | BBOF1 | Diff. RPE1 GFP-Centrin | Random |
| ENSG00000174483 | ENST00000318312 | BBS1 | Diff. RPE1 GFP-Centrin | Random |
| ENSG00000179941 | ENST00000393262 | BBS10 | Diff. RPE1 GFP-Centrin | Random |
| ENSG00000181004 | ENST00000314218 | BBS12 | Diff. RPE1 GFP-Centrin | Random |
| ENSG00000125124 | ENST00000245157 | BBS2 | Diff. RPE1 GFP-Centrin | Random |
| ENSG00000140463 | ENST00000268057 | BBS4 | Diff. RPE1 GFP-Centrin | Random |
| ENSG00000140463 | ENST00000268057 | BBS4 | Diff. RPE1 GFP-Centrin | Random |
| ENSG00000163093 | ENST00000295240 | BBS5 | Diff. RPE1 GFP-Centrin | Random |
| ENSG00000138686 | ENST00000264499 | BBS7 | Diff. RPE1 GFP-Centrin | Random |
| ENSG00000122507 | ENST00000242067 | BBS9 | Diff. RPE1 GFP-Centrin | Random |

| | | | | |
|---|---|---|---|---|
| ENSG00000116752 | ENST00000369541 | BCAS2 | Diff. RPE1 GFP-Centrin | Random |
| ENSG00000116752 | ENST00000369541 | BCAS2 | Diff. RPE1 GFP-Centrin | Random |
| ENSG00000107949 | ENST00000278100 | BCCIP | Diff. RPE1 GFP-Centrin | Random |
| ENSG00000069399 | ENST00000164227 | BCL3 | Diff. RPE1 GFP-Centrin | Random |
| ENSG00000115760 | ENST00000421745 | BIRC6 | Diff. RPE1 GFP-Centrin | Random |
| ENSG00000145919 | ENST00000311086 | BOD1 | Diff. RPE1 GFP-Centrin | Random |
| ENSG00000169679 | ENST00000535254 | BUB1 | Diff. RPE1 GFP-Centrin | Random |
| ENSG00000156970 | ENST00000287598 | BUB1B | Diff. RPE1 GFP-Centrin | Random |
| ENSG00000154473 | ENST00000368858 | BUB3 | Diff. RPE1 GFP-Centrin | Random |
| ENSG00000265590 | ENST00000673807 | C21ORF59 | Diff. RPE1 GFP-Centrin | Random |
| ENSG00000198663 | ENST00000480824 | C6ORF89 | Diff. RPE1 GFP-Centrin | Random |
| ENSG00000153790 | ENST00000283905 | C7ORF31 | Diff. RPE1 GFP-Centrin | Random |
| ENSG00000183346 | ENST00000330194 | CABCOCO1 | Diff. RPE1 GFP-Centrin | Random |
| ENSG00000154040 | ENST00000327201 | CABYR | Diff. RPE1 GFP-Centrin | Random |
| ENSG00000163888 | ENST00000296238 | CAMK2N2 | Diff. RPE1 GFP-Centrin | Random |
| ENSG00000130559 | ENST00000389532 | CAMSAP1 | Diff. RPE1 GFP-Centrin | Random |
| ENSG00000118200 | ENST00000358823 | CAMSAP2 | Diff. RPE1 GFP-Centrin | Random |
| ENSG00000175294 | ENST00000312106 | CATSPER1 | Diff. RPE1 GFP-Centrin | Random |
| ENSG00000105974 | ENST00000405348 | CAV1 | Diff. RPE1 GFP-Centrin | Random |
| ENSG00000007080 | ENST00000445755 | CCDC124 | Diff. RPE1 GFP-Centrin | Random |
| ENSG00000175455 | ENST00000310351 | CCDC14 | Diff. RPE1 GFP-Centrin | Random |
| ENSG00000149548 | ENST00000344762 | CCDC15 | Diff. RPE1 GFP-Centrin | Random |
| ENSG00000119242 | ENST00000238156 | CCDC92 | Diff. RPE1 GFP-Centrin | Random |
| ENSG00000204536 | ENST00000376266 | CCHCR1 | Diff. RPE1 GFP-Centrin | Random |
| ENSG00000162063 | ENST00000397066 | CCNF | Diff. RPE1 GFP-Centrin | Random |
| ENSG00000103540 | ENST00000396208 | CCP110 | Diff. RPE1 GFP-Centrin | Random |
| ENSG00000103540 | ENST00000396208 | CCP110 | Diff. RPE1 GFP-Centrin | Random |
| ENSG00000154429 | ENST00000284617 | CCSAP | Diff. RPE1 GFP-Centrin | Random |
| ENSG00000163468 | ENST00000295688 | CCT3 | Diff. RPE1 GFP-Centrin | Random |
| ENSG00000115484 | ENST00000544079 | CCT4 | Diff. RPE1 GFP-Centrin | Random |
| ENSG00000150753 | ENST00000280326 | CCT5 | Diff. RPE1 GFP-Centrin | Random |
| ENSG00000150753 | ENST00000503026 | CCT5 | Diff. RPE1 GFP-Centrin | Random |
| ENSG00000146731 | ENST00000275603 | CCT6A | Diff. RPE1 GFP-Centrin | Random |
| ENSG00000135624 | ENST00000258091 | CCT7 | Diff. RPE1 GFP-Centrin | Random |
| ENSG00000156261 | ENST00000286788 | CCT8 | Diff. RPE1 GFP-Centrin | Random |
| ENSG00000079335 | ENST00000336454 | CDC14A | Diff. RPE1 GFP-Centrin | Random |
| ENSG00000079335 | ENST00000361544 | CDC14A | Diff. RPE1 GFP-Centrin | Random |
| ENSG00000081377 | ENST00000412285 | CDC14B | Diff. RPE1 GFP-Centrin | Random |
| ENSG00000130177 | ENST00000356221 | CDC16 | Diff. RPE1 GFP-Centrin | Random |
| ENSG00000130177 | ENST00000360383 | CDC16 | Diff. RPE1 GFP-Centrin | Random |
| ENSG00000130177 | ENST00000360383 | CDC16 | Diff. RPE1 GFP-Centrin | Random |
| ENSG00000117399 | ENST00000310955 | CDC20 | Diff. RPE1 GFP-Centrin | Random |
| ENSG00000117399 | ENST00000310955 | CDC20 | Diff. RPE1 GFP-Centrin | Random |
| ENSG00000004897 | ENST00000531206 | CDC27 | Diff. RPE1 GFP-Centrin | Random |
| ENSG00000070831 | ENST00000400259 | CDC42 | Diff. RPE1 GFP-Centrin | Random |
| ENSG00000097046 | ENST00000234626 | CDC7 | Diff. RPE1 GFP-Centrin | Random |
| ENSG00000134690 | ENST00000327331 | CDCA8 | Diff. RPE1 GFP-Centrin | Random |
| ENSG00000156345 | ENST00000375871 | CDK20 | Diff. RPE1 GFP-Centrin | Random |
| ENSG00000167797 | ENST00000301488 | CDK2AP2 | Diff. RPE1 GFP-Centrin | Random |
| ENSG00000164885 | ENST00000485972 | CDK5 | Diff. RPE1 GFP-Centrin | Random |
| ENSG00000108465 | ENST00000338399 | CDK5RAP3 | Diff. RPE1 GFP-Centrin | Random |
| ENSG00000105810 | ENST00000265734 | CDK6 | Diff. RPE1 GFP-Centrin | Random |
| ENSG00000115163 | ENST00000335756 | CENPA | Diff. RPE1 GFP-Centrin | Random |

| | | | | |
|---|---|---|---|---|
| ENSG00000138778 | ENST00000380026 | CENPE | Diff. RPE1 GFP-Centrin | Random |
| ENSG00000151849 | ENST00000381884 | CENPJ | Diff. RPE1 GFP-Centrin | Random |
| ENSG00000138092 | ENST00000473706 | CENPO | Diff. RPE1 GFP-Centrin | Random |
| ENSG00000175279 | ENST00000477755 | CENPS | Diff. RPE1 GFP-Centrin | Random |
| ENSG00000166582 | ENST00000299736 | CENPV | Diff. RPE1 GFP-Centrin | Random |
| ENSG00000203760 | ENST00000368328 | CENPW | Diff. RPE1 GFP-Centrin | Random |
| ENSG00000169689 | ENST00000392359 | CENPX | Diff. RPE1 GFP-Centrin | Random |
| ENSG00000116198 | ENST00000378230 | CEP104 | Diff. RPE1 GFP-Centrin | Random |
| ENSG00000154240 | ENST00000535342 | CEP112 | Diff. RPE1 GFP-Centrin | Random |
| ENSG00000168944 | ENST00000306481 | CEP120 | Diff. RPE1 GFP-Centrin | Random |
| ENSG00000135315 | ENST00000403245 | CEP162 | Diff. RPE1 GFP-Centrin | Random |
| ENSG00000135315 | ENST00000403245 | CEP162 | Diff. RPE1 GFP-Centrin | Random |
| ENSG00000143702 | ENST00000366542 | CEP170 | Diff. RPE1 GFP-Centrin | Random |
| ENSG00000099814 | ENST00000556508 | CEP170B | Diff. RPE1 GFP-Centrin | Random |
| ENSG00000174007 | ENST00000409690 | CEP19 | Diff. RPE1 GFP-Centrin | Random |
| ENSG00000164118 | ENST00000296519 | CEP44 | Diff. RPE1 GFP-Centrin | Random |
| ENSG00000182923 | ENST00000332047 | CEP63 | Diff. RPE1 GFP-Centrin | Random |
| ENSG00000011523 | ENST00000377990 | CEP68 | Diff. RPE1 GFP-Centrin | Random |
| ENSG00000173588 | ENST00000339839 | CEP83 | Diff. RPE1 GFP-Centrin | Random |
| ENSG00000130695 | ENST00000252992 | CEP85 | Diff. RPE1 GFP-Centrin | Random |
| ENSG00000121289 | ENST00000305768 | CEP89 | Diff. RPE1 GFP-Centrin | Random |
| ENSG00000177143 | ENST00000327228 | CETN1 | Diff. RPE1 GFP-Centrin | Random |
| ENSG00000147400 | ENST00000370277 | CETN2 | Diff. RPE1 GFP-Centrin | Random |
| ENSG00000188596 | ENST00000524981 | CFAP54 | Diff. RPE1 GFP-Centrin | Random |
| ENSG00000120051 | ENST00000369704 | CFAP58 | Diff. RPE1 GFP-Centrin | Random |
| ENSG00000153774 | ENST00000283882 | CFDP1 | Diff. RPE1 GFP-Centrin | Random |
| ENSG00000254505 | ENST00000347519 | CHMP4A | Diff. RPE1 GFP-Centrin | Random |
| ENSG00000101421 | ENST00000217402 | CHMP4B | Diff. RPE1 GFP-Centrin | Random |
| ENSG00000110172 | ENST00000320585 | CHORDC1 | Diff. RPE1 GFP-Centrin | Random |
| ENSG00000110172 | ENST00000457199 | CHORDC1 | Diff. RPE1 GFP-Centrin | Random |
| ENSG00000138433 | ENST00000342016 | CIR1 | Diff. RPE1 GFP-Centrin | Random |
| ENSG00000136108 | ENST00000378037 | CKAP2 | Diff. RPE1 GFP-Centrin | Random |
| ENSG00000130779 | ENST00000620786 | CLIP1 | Diff. RPE1 GFP-Centrin | Random |
| ENSG00000106665 | ENST00000361545 | CLIP2 | Diff. RPE1 GFP-Centrin | Random |
| ENSG00000070371 | ENST00000442042 | CLTCL1 | Diff. RPE1 GFP-Centrin | Random |
| ENSG00000103351 | ENST00000417763 | CLUAP1 | Diff. RPE1 GFP-Centrin | Random |
| ENSG00000044459 | ENST00000380647 | CNTLN | Diff. RPE1 GFP-Centrin | Random |
| ENSG00000170037 | ENST00000563694 | CNTROB | Diff. RPE1 GFP-Centrin | Random |
| ENSG00000099942 | ENST00000354336 | CRKL | Diff. RPE1 GFP-Centrin | Random |
| ENSG00000113712 | ENST00000261798 | CSNK1A1 | Diff. RPE1 GFP-Centrin | Random |
| ENSG00000113712 | ENST00000377843 | CSNK1A1 | Diff. RPE1 GFP-Centrin | Random |
| ENSG00000204435 | ENST00000375882 | CSNK2B | Diff. RPE1 GFP-Centrin | Random |
| ENSG00000104218 | ENST00000262210 | CSPP1 | Diff. RPE1 GFP-Centrin | Random |
| ENSG00000060069 | ENST00000613122 | CTDP1 | Diff. RPE1 GFP-Centrin | Random |
| ENSG00000198561 | ENST00000526772 | CTNND1 | Diff. RPE1 GFP-Centrin | Random |
| ENSG00000044090 | ENST00000265348 | CUL7 | Diff. RPE1 GFP-Centrin | Random |
| ENSG00000044090 | ENST00000535468 | CUL7 | Diff. RPE1 GFP-Centrin | Random |
| ENSG00000205795 | ENST00000381813 | CYS1 | Diff. RPE1 GFP-Centrin | Random |
| ENSG00000123977 | ENST00000309931 | DAW1 | Diff. RPE1 GFP-Centrin | Random |
| ENSG00000198876 | ENST00000361264 | DCAF12 | Diff. RPE1 GFP-Centrin | Random |
| ENSG00000164934 | ENST00000297579 | DCAF13 | Diff. RPE1 GFP-Centrin | Random |
| ENSG00000164934 | ENST00000297579 | DCAF13 | Diff. RPE1 GFP-Centrin | Random |
| ENSG00000118655 | ENST00000369563 | DCLRE1B | Diff. RPE1 GFP-Centrin | Random |
| ENSG00000137100 | ENST00000259632 | DCTN3 | Diff. RPE1 GFP-Centrin | Random |

| | | | | |
|---|---|---|---|---|
| ENSG00000132912 | ENST00000447998 | DCTN4 | Diff. RPE1 GFP-Centrin | Random |
| ENSG00000104671 | ENST00000221114 | DCTN6 | Diff. RPE1 GFP-Centrin | Random |
| ENSG00000135829 | ENST00000367549 | DHX9 | Diff. RPE1 GFP-Centrin | Random |
| ENSG00000091140 | ENST00000440410 | DLD | Diff. RPE1 GFP-Centrin | Random |
| ENSG00000075711 | ENST00000357674 | DLG1 | Diff. RPE1 GFP-Centrin | Random |
| ENSG00000256061 | ENST00000457155 | DNAAF4 | Diff. RPE1 GFP-Centrin | Random |
| ENSG00000100246 | ENST00000216068 | DNAL4 | Diff. RPE1 GFP-Centrin | Random |
| ENSG00000106976 | ENST00000372923 | DNM1 | Diff. RPE1 GFP-Centrin | Random |
| ENSG00000087470 | ENST00000452533 | DNM1L | Diff. RPE1 GFP-Centrin | Random |
| ENSG00000133884 | ENST00000252268 | DPF2 | Diff. RPE1 GFP-Centrin | Random |
| ENSG00000092964 | ENST00000311151 | DPYSL2 | Diff. RPE1 GFP-Centrin | Random |
| ENSG00000185721 | ENST00000331457 | DRG1 | Diff. RPE1 GFP-Centrin | Random |
| ENSG00000143476 | ENST00000366991 | DTL | Diff. RPE1 GFP-Centrin | Random |
| ENSG00000107404 | ENST00000378891 | DVL1 | Diff. RPE1 GFP-Centrin | Random |
| ENSG00000144635 | ENST00000273130 | DYNC1LI1 | Diff. RPE1 GFP-Centrin | Random |
| ENSG00000135720 | ENST00000258198 | DYNC1LI2 | Diff. RPE1 GFP-Centrin | Random |
| ENSG00000138036 | ENST00000260605 | DYNC2LI1 | Diff. RPE1 GFP-Centrin | Random |
| ENSG00000264364 | ENST00000579991 | DYNLL2 | Diff. RPE1 GFP-Centrin | Random |
| ENSG00000125971 | ENST00000357156 | DYNLRB1 | Diff. RPE1 GFP-Centrin | Random |
| ENSG00000165169 | ENST00000378578 | DYNLT3 | Diff. RPE1 GFP-Centrin | Random |
| ENSG00000134874 | ENST00000361396 | DZIP1 | Diff. RPE1 GFP-Centrin | Random |
| ENSG00000158163 | ENST00000327532 | DZIP1L | Diff. RPE1 GFP-Centrin | Random |
| ENSG00000101412 | ENST00000343380 | E2F1 | Diff. RPE1 GFP-Centrin | Random |
| ENSG00000136813 | ENST00000259335 | ECPAS | Diff. RPE1 GFP-Centrin | Random |
| ENSG00000203965 | ENST00000371088 | EFCAB7 | Diff. RPE1 GFP-Centrin | Random |
| ENSG00000096093 | ENST00000371068 | EFHC1 | Diff. RPE1 GFP-Centrin | Random |
| ENSG00000013016 | ENST00000322054 | EHD3 | Diff. RPE1 GFP-Centrin | Random |
| ENSG00000102119 | ENST00000369842 | EMD | Diff. RPE1 GFP-Centrin | Random |
| ENSG00000125746 | ENST00000245925 | EML2 | Diff. RPE1 GFP-Centrin | Random |
| ENSG00000186871 | ENST00000334463 | ERCC6L | Diff. RPE1 GFP-Centrin | Random |
| ENSG00000186871 | ENST00000334463 | ERCC6L | Diff. RPE1 GFP-Centrin | Random |
| ENSG00000100632 | ENST00000557016 | ERH | Diff. RPE1 GFP-Centrin | Random |
| ENSG00000135476 | ENST00000257934 | ESPL1 | Diff. RPE1 GFP-Centrin | Random |
| ENSG00000072840 | ENST00000382674 | EVC | Diff. RPE1 GFP-Centrin | Random |
| ENSG00000180104 | ENST00000512944 | EXOC3 | Diff. RPE1 GFP-Centrin | Random |
| ENSG00000158161 | ENST00000436342 | EYA3 | Diff. RPE1 GFP-Centrin | Random |
| ENSG00000158161 | ENST00000436342 | EYA3 | Diff. RPE1 GFP-Centrin | Random |
| ENSG00000170264 | ENST00000405894 | FAM161A | Diff. RPE1 GFP-Centrin | Random |
| ENSG00000153310 | ENST00000517654 | FAM49B | Diff. RPE1 GFP-Centrin | Random |
| ENSG00000143756 | ENST00000366862 | FBXO28 | Diff. RPE1 GFP-Centrin | Random |
| ENSG00000103264 | ENST00000565593 | FBXO31 | Diff. RPE1 GFP-Centrin | Random |
| ENSG00000149557 | ENST00000278919 | FEZ1 | Diff. RPE1 GFP-Centrin | Random |
| ENSG00000004478 | ENST00000001008 | FKBP4 | Diff. RPE1 GFP-Centrin | Random |
| ENSG00000154803 | ENST00000285071 | FLCN | Diff. RPE1 GFP-Centrin | Random |
| ENSG00000154803 | ENST00000285071 | FLCN | Diff. RPE1 GFP-Centrin | Random |
| ENSG00000133393 | ENST00000255759 | FOPNL | Diff. RPE1 GFP-Centrin | Random |
| ENSG00000163820 | ENST00000296137 | FYCO1 | Diff. RPE1 GFP-Centrin | Random |
| ENSG00000163820 | ENST00000433878 | FYCO1 | Diff. RPE1 GFP-Centrin | Random |
| ENSG00000139112 | ENST00000266458 | GABARAPL1 | Diff. RPE1 GFP-Centrin | Random |
| ENSG00000141013 | ENST00000268699 | GAS8 | Diff. RPE1 GFP-Centrin | Random |
| ENSG00000178295 | ENST00000381254 | GEN1 | Diff. RPE1 GFP-Centrin | Random |
| ENSG00000074047 | ENST00000361492 | GLI2 | Diff. RPE1 GFP-Centrin | Random |
| ENSG00000106571 | ENST00000395925 | GLI3 | Diff. RPE1 GFP-Centrin | Random |

| | | | | |
|---|---|---|---|---|
| ENSG00000127955 | ENST00000351004 | GNAI1 | Diff. RPE1 GFP-Centrin | Random |
| ENSG00000127955 | ENST00000351004 | GNAI1 | Diff. RPE1 GFP-Centrin | Random |
| ENSG00000114353 | ENST00000266027 | GNAI2 | Diff. RPE1 GFP-Centrin | Random |
| ENSG00000167110 | ENST00000421699 | GOLGA2 | Diff. RPE1 GFP-Centrin | Random |
| ENSG00000143147 | ENST00000271357 | GPR161 | Diff. RPE1 GFP-Centrin | Random |
| ENSG00000143147 | ENST00000271357 | GPR161 | Diff. RPE1 GFP-Centrin | Random |
| ENSG00000173020 | ENST00000308595 | GRK2 | Diff. RPE1 GFP-Centrin | Random |
| ENSG00000082701 | ENST00000264235 | GSK3B | Diff. RPE1 GFP-Centrin | Random |
| ENSG00000188486 | ENST00000530167 | H2AFX | Diff. RPE1 GFP-Centrin | Random |
| ENSG00000113648 | ENST00000312469 | H2AFY | Diff. RPE1 GFP-Centrin | Random |
| ENSG00000177602 | ENST00000325418 | HASPIN | Diff. RPE1 GFP-Centrin | Random |
| ENSG00000152240 | ENST00000282058 | HAUS1 | Diff. RPE1 GFP-Centrin | Random |
| ENSG00000214367 | ENST00000443786 | HAUS3 | Diff. RPE1 GFP-Centrin | Random |
| ENSG00000214367 | ENST00000443786 | HAUS3 | Diff. RPE1 GFP-Centrin | Random |
| ENSG00000092036 | ENST00000541587 | HAUS4 | Diff. RPE1 GFP-Centrin | Random |
| ENSG00000249115 | ENST00000203166 | HAUS5 | Diff. RPE1 GFP-Centrin | Random |
| ENSG00000147874 | ENST00000380502 | HAUS6 | Diff. RPE1 GFP-Centrin | Random |
| ENSG00000147874 | ENST00000380502 | HAUS6 | Diff. RPE1 GFP-Centrin | Random |
| ENSG00000213397 | ENST00000370211 | HAUS7 | Diff. RPE1 GFP-Centrin | Random |
| ENSG00000213397 | ENST00000370211 | HAUS7 | Diff. RPE1 GFP-Centrin | Random |
| ENSG00000128731 | ENST00000261609 | HERC2 | Diff. RPE1 GFP-Centrin | Random |
| ENSG00000156515 | ENST00000359426 | HK1 | Diff. RPE1 GFP-Centrin | Random |
| ENSG00000095066 | ENST00000264827 | HOOK2 | Diff. RPE1 GFP-Centrin | Random |
| ENSG00000168172 | ENST00000307602 | HOOK3 | Diff. RPE1 GFP-Centrin | Random |
| ENSG00000185122 | ENST00000528838 | HSF1 | Diff. RPE1 GFP-Centrin | Random |
| ENSG00000204389 | ENST00000375651 | HSPA1A | Diff. RPE1 GFP-Centrin | Random |
| ENSG00000204388 | ENST00000375650 | HSPA1B | Diff. RPE1 GFP-Centrin | Random |
| ENSG00000173110 | ENST00000309758 | HSPA6 | Diff. RPE1 GFP-Centrin | Random |
| ENSG00000081870 | ENST00000194214 | HSPB11 | Diff. RPE1 GFP-Centrin | Random |
| ENSG00000197386 | ENST00000355072 | HTT | Diff. RPE1 GFP-Centrin | Random |
| ENSG00000112144 | ENST00000350082 | ICK | Diff. RPE1 GFP-Centrin | Random |
| ENSG00000187535 | ENST00000426508 | IFT140 | Diff. RPE1 GFP-Centrin | Random |
| ENSG00000138002 | ENST00000260570 | IFT172 | Diff. RPE1 GFP-Centrin | Random |
| ENSG00000109083 | ENST00000395418 | IFT20 | Diff. RPE1 GFP-Centrin | Random |
| ENSG00000128581 | ENST00000315322 | IFT22 | Diff. RPE1 GFP-Centrin | Random |
| ENSG00000100360 | ENST00000433985 | IFT27 | Diff. RPE1 GFP-Centrin | Random |
| ENSG00000100360 | ENST00000433985 | IFT27 | Diff. RPE1 GFP-Centrin | Random |
| ENSG00000118096 | ENST00000264021 | IFT46 | Diff. RPE1 GFP-Centrin | Random |
| ENSG00000101052 | ENST00000373039 | IFT52 | Diff. RPE1 GFP-Centrin | Random |
| ENSG00000114446 | ENST00000264538 | IFT57 | Diff. RPE1 GFP-Centrin | Random |
| ENSG00000096872 | ENST00000380062 | IFT74 | Diff. RPE1 GFP-Centrin | Random |
| ENSG00000122970 | ENST00000552912 | IFT81 | Diff. RPE1 GFP-Centrin | Random |
| ENSG00000032742 | ENST00000351808 | IFT88 | Diff. RPE1 GFP-Centrin | Random |
| ENSG00000128908 | ENST00000361937 | INO80 | Diff. RPE1 GFP-Centrin | Random |
| ENSG00000148384 | ENST00000371712 | INPP5E | Diff. RPE1 GFP-Centrin | Random |
| ENSG00000173226 | ENST00000310864 | IQCB1 | Diff. RPE1 GFP-Centrin | Random |
| ENSG00000106012 | ENST00000402050 | IQCE | Diff. RPE1 GFP-Centrin | Random |
| ENSG00000106012 | ENST00000402050 | IQCE | Diff. RPE1 GFP-Centrin | Random |
| ENSG00000114473 | ENST00000265239 | IQCG | Diff. RPE1 GFP-Centrin | Random |
| ENSG00000140575 | ENST00000268182 | IQGAP1 | Diff. RPE1 GFP-Centrin | Random |
| ENSG00000142856 | ENST00000271002 | ITGB3BP | Diff. RPE1 GFP-Centrin | Random |
| ENSG00000143543 | ENST00000271843 | JTB | Diff. RPE1 GFP-Centrin | Random |
| ENSG00000108773 | ENST00000225916 | KAT2A | Diff. RPE1 GFP-Centrin | Random |

| | | | | |
|---|---|---|---|---|
| ENSG00000103510 | ENST00000219797 | KAT8 | Diff. RPE1 GFP-Centrin | Random |
| ENSG00000186625 | ENST00000367411 | KATNA1 | Diff. RPE1 GFP-Centrin | Random |
| ENSG00000186625 | ENST00000367411 | KATNA1 | Diff. RPE1 GFP-Centrin | Random |
| ENSG00000102781 | ENST00000380615 | KATNAL1 | Diff. RPE1 GFP-Centrin | Random |
| ENSG00000140854 | ENST00000379661 | KATNB1 | Diff. RPE1 GFP-Centrin | Random |
| ENSG00000198920 | ENST00000361413 | KIAA0753 | Diff. RPE1 GFP-Centrin | Random |
| ENSG00000197892 | ENST00000524189 | KIF13B | Diff. RPE1 GFP-Centrin | Random |
| ENSG00000197892 | ENST00000524189 | KIF13B | Diff. RPE1 GFP-Centrin | Random |
| ENSG00000163808 | ENST00000326047 | KIF15 | Diff. RPE1 GFP-Centrin | Random |
| ENSG00000137807 | ENST00000260363 | KIF23 | Diff. RPE1 GFP-Centrin | Random |
| ENSG00000068796 | ENST00000381103 | KIF2A | Diff. RPE1 GFP-Centrin | Random |
| ENSG00000068796 | ENST00000401507 | KIF2A | Diff. RPE1 GFP-Centrin | Random |
| ENSG00000101350 | ENST00000375712 | KIF3B | Diff. RPE1 GFP-Centrin | Random |
| ENSG00000101350 | ENST00000375712 | KIF3B | Diff. RPE1 GFP-Centrin | Random |
| ENSG00000090889 | ENST00000374403 | KIF4A | Diff. RPE1 GFP-Centrin | Random |
| ENSG00000166813 | ENST00000394412 | KIF7 | Diff. RPE1 GFP-Centrin | Random |
| ENSG00000075945 | ENST00000361580 | KIFAP3 | Diff. RPE1 GFP-Centrin | Random |
| ENSG00000137171 | ENST00000347162 | KLC4 | Diff. RPE1 GFP-Centrin | Random |
| ENSG00000003096 | ENST00000371882 | KLHL13 | Diff. RPE1 GFP-Centrin | Random |
| ENSG00000162413 | ENST00000377658 | KLHL21 | Diff. RPE1 GFP-Centrin | Random |
| ENSG00000162413 | ENST00000377658 | KLHL21 | Diff. RPE1 GFP-Centrin | Random |
| ENSG00000099910 | ENST00000328879 | KLHL22 | Diff. RPE1 GFP-Centrin | Random |
| ENSG00000198642 | ENST00000359039 | KLHL9 | Diff. RPE1 GFP-Centrin | Random |
| ENSG00000128944 | ENST00000608100 | KNSTRN | Diff. RPE1 GFP-Centrin | Random |
| ENSG00000184445 | ENST00000333479 | KNTC1 | Diff. RPE1 GFP-Centrin | Random |
| ENSG00000135338 | ENST00000369846 | LCA5 | Diff. RPE1 GFP-Centrin | Random |
| ENSG00000166477 | ENST00000299601 | LEO1 | Diff. RPE1 GFP-Centrin | Random |
| ENSG00000169683 | ENST00000306688 | LRRC45 | Diff. RPE1 GFP-Centrin | Random |
| ENSG00000133739 | ENST00000360375 | LRRCC1 | Diff. RPE1 GFP-Centrin | Random |
| ENSG00000133739 | ENST00000360375 | LRRCC1 | Diff. RPE1 GFP-Centrin | Random |
| ENSG00000107816 | ENST00000370223 | LZTS2 | Diff. RPE1 GFP-Centrin | Random |
| ENSG00000107816 | ENST00000370223 | LZTS2 | Diff. RPE1 GFP-Centrin | Random |
| ENSG00000002822 | ENST00000399654 | MAD1L1 | Diff. RPE1 GFP-Centrin | Random |
| ENSG00000002822 | ENST00000399654 | MAD1L1 | Diff. RPE1 GFP-Centrin | Random |
| ENSG00000179632 | ENST00000322428 | MAF1 | Diff. RPE1 GFP-Centrin | Random |
| ENSG00000130479 | ENST00000324096 | MAP1S | Diff. RPE1 GFP-Centrin | Random |
| ENSG00000126934 | ENST00000262948 | MAP2K2 | Diff. RPE1 GFP-Centrin | Random |
| ENSG00000047849 | ENST00000429422 | MAP4 | Diff. RPE1 GFP-Centrin | Random |
| ENSG00000101367 | ENST00000375571 | MAPRE1 | Diff. RPE1 GFP-Centrin | Random |
| ENSG00000166974 | ENST00000300249 | MAPRE2 | Diff. RPE1 GFP-Centrin | Random |
| ENSG00000084764 | ENST00000233121 | MAPRE3 | Diff. RPE1 GFP-Centrin | Random |
| ENSG00000007047 | ENST00000262891 | MARK4 | Diff. RPE1 GFP-Centrin | Random |
| ENSG00000152601 | ENST00000463374 | MBNL1 | Diff. RPE1 GFP-Centrin | Random |
| ENSG00000163875 | ENST00000296214 | MEAF6 | Diff. RPE1 GFP-Centrin | Random |
| ENSG00000169057 | ENST00000303391 | MECP2 | Diff. RPE1 GFP-Centrin | Random |
| ENSG00000184634 | ENST00000374080 | MED12 | Diff. RPE1 GFP-Centrin | Random |
| ENSG00000101752 | ENST00000261537 | MIB1 | Diff. RPE1 GFP-Centrin | Random |
| ENSG00000101871 | ENST00000317552 | MID1 | Diff. RPE1 GFP-Centrin | Random |
| ENSG00000125863 | ENST00000399054 | MKKS | Diff. RPE1 GFP-Centrin | Random |
| ENSG00000011143 | ENST00000393119 | MKS1 | Diff. RPE1 GFP-Centrin | Random |
| ENSG00000011143 | ENST00000393119 | MKS1 | Diff. RPE1 GFP-Centrin | Random |
| ENSG00000076242 | ENST00000231790 | MLH1 | Diff. RPE1 GFP-Centrin | Random |
| ENSG00000168303 | ENST00000306984 | MPLKIP | Diff. RPE1 GFP-Centrin | Random |

| | | | | |
|---|---|---|---|---|
| ENSG00000172167 | ENST00000305949 | MTBP | Diff. RPE1 GFP-Centrin | Random |
| ENSG00000133026 | ENST00000269243 | MYH10 | Diff. RPE1 GFP-Centrin | Random |
| ENSG00000204899 | ENST00000377818 | MZT1 | Diff. RPE1 GFP-Centrin | Random |
| ENSG00000135372 | ENST00000257829 | NAT10 | Diff. RPE1 GFP-Centrin | Random |
| ENSG00000163382 | ENST00000368235 | NAXE | Diff. RPE1 GFP-Centrin | Random |
| ENSG00000010292 | ENST00000315579 | NCAPD2 | Diff. RPE1 GFP-Centrin | Random |
| ENSG00000109805 | ENST00000251496 | NCAPG | Diff. RPE1 GFP-Centrin | Random |
| ENSG00000167566 | ENST00000335999 | NCKAP5L | Diff. RPE1 GFP-Centrin | Random |
| ENSG00000166579 | ENST00000334527 | NDEL1 | Diff. RPE1 GFP-Centrin | Random |
| ENSG00000166579 | ENST00000402554 | NDEL1 | Diff. RPE1 GFP-Centrin | Random |
| ENSG00000104419 | ENST00000323851 | NDRG1 | Diff. RPE1 GFP-Centrin | Random |
| ENSG00000137601 | ENST00000507142 | NEK1 | Diff. RPE1 GFP-Centrin | Random |
| ENSG00000137601 | ENST00000511633 | NEK1 | Diff. RPE1 GFP-Centrin | Random |
| ENSG00000117650 | ENST00000366999 | NEK2 | Diff. RPE1 GFP-Centrin | Random |
| ENSG00000136098 | ENST00000610828 | NEK3 | Diff. RPE1 GFP-Centrin | Random |
| ENSG00000114904 | ENST00000233027 | NEK4 | Diff. RPE1 GFP-Centrin | Random |
| ENSG00000119408 | ENST00000320246 | NEK6 | Diff. RPE1 GFP-Centrin | Random |
| ENSG00000151414 | ENST00000367385 | NEK7 | Diff. RPE1 GFP-Centrin | Random |
| ENSG00000119638 | ENST00000238616 | NEK9 | Diff. RPE1 GFP-Centrin | Random |
| ENSG00000119638 | ENST00000238616 | NEK9 | Diff. RPE1 GFP-Centrin | Random |
| ENSG00000196712 | ENST00000358273 | NF1 | Diff. RPE1 GFP-Centrin | Random |
| ENSG00000116044 | ENST00000397063 | NFE2L2 | Diff. RPE1 GFP-Centrin | Random |
| ENSG00000100906 | ENST00000216797 | NFKBIA | Diff. RPE1 GFP-Centrin | Random |
| ENSG00000145029 | ENST00000273598 | NICN1 | Diff. RPE1 GFP-Centrin | Random |
| ENSG00000239672 | ENST00000393196 | NME1 | Diff. RPE1 GFP-Centrin | Random |
| ENSG00000144061 | ENST00000445609 | NPHP1 | Diff. RPE1 GFP-Centrin | Random |
| ENSG00000113971 | ENST00000337331 | NPHP3 | Diff. RPE1 GFP-Centrin | Random |
| ENSG00000131697 | ENST00000378156 | NPHP4 | Diff. RPE1 GFP-Centrin | Random |
| ENSG00000181163 | ENST00000296930 | NPM1 | Diff. RPE1 GFP-Centrin | Random |
| ENSG00000181163 | ENST00000517671 | NPM1 | Diff. RPE1 GFP-Centrin | Random |
| ENSG00000117697 | ENST00000422588 | NSL1 | Diff. RPE1 GFP-Centrin | Random |
| ENSG00000103274 | ENST00000283027 | NUBP1 | Diff. RPE1 GFP-Centrin | Random |
| ENSG00000095906 | ENST00000262302 | NUBP2 | Diff. RPE1 GFP-Centrin | Random |
| ENSG00000090273 | ENST00000321265 | NUDC | Diff. RPE1 GFP-Centrin | Random |
| ENSG00000167005 | ENST00000300291 | NUDT21 | Diff. RPE1 GFP-Centrin | Random |
| ENSG00000111581 | ENST00000229179 | NUP107 | Diff. RPE1 GFP-Centrin | Random |
| ENSG00000069248 | ENST00000261396 | NUP133 | Diff. RPE1 GFP-Centrin | Random |
| ENSG00000125450 | ENST00000245544 | NUP85 | Diff. RPE1 GFP-Centrin | Random |
| ENSG00000137804 | ENST00000414849 | NUSAP1 | Diff. RPE1 GFP-Centrin | Random |
| ENSG00000124006 | ENST00000404537 | OBSL1 | Diff. RPE1 GFP-Centrin | Random |
| ENSG00000122126 | ENST00000357121 | OCRL | Diff. RPE1 GFP-Centrin | Random |
| ENSG00000138430 | ENST00000344357 | OLA1 | Diff. RPE1 GFP-Centrin | Random |
| ENSG00000115942 | ENST00000234296 | ORC2 | Diff. RPE1 GFP-Centrin | Random |
| ENSG00000102981 | ENST00000458121 | PARD6A | Diff. RPE1 GFP-Centrin | Random |
| ENSG00000102981 | ENST00000458121 | PARD6A | Diff. RPE1 GFP-Centrin | Random |
| ENSG00000041880 | ENST00000398755 | PARP3 | Diff. RPE1 GFP-Centrin | Random |
| ENSG00000132849 | ENST00000371158 | PATJ | Diff. RPE1 GFP-Centrin | Random |
| ENSG00000166803 | ENST00000559519 | PCLAF | Diff. RPE1 GFP-Centrin | Random |
| ENSG00000078674 | ENST00000325083 | PCM1 | Diff. RPE1 GFP-Centrin | Random |
| ENSG00000132646 | ENST00000379143 | PCNA | Diff. RPE1 GFP-Centrin | Random |
| ENSG00000154678 | ENST00000396193 | PDE1C | Diff. RPE1 GFP-Centrin | Random |
| ENSG00000113448 | ENST00000340635 | PDE4D | Diff. RPE1 GFP-Centrin | Random |
| ENSG00000156973 | ENST00000287600 | PDE6D | Diff. RPE1 GFP-Centrin | Random |

| | | | | |
|---|---|---|---|---|
| ENSG00000241360 | ENST00000215904 | PDXP | Diff. RPE1 GFP-Centrin | Random |
| ENSG00000100029 | ENST00000354694 | PES1 | Diff. RPE1 GFP-Centrin | Random |
| ENSG00000170950 | ENST00000304801 | PGK2 | Diff. RPE1 GFP-Centrin | Random |
| ENSG00000164902 | ENST00000297540 | PHAX | Diff. RPE1 GFP-Centrin | Random |
| ENSG00000197724 | ENST00000359246 | PHF2 | Diff. RPE1 GFP-Centrin | Random |
| ENSG00000083535 | ENST00000326291 | PIBF1 | Diff. RPE1 GFP-Centrin | Random |
| ENSG00000051382 | ENST00000477593 | PIK3CB | Diff. RPE1 GFP-Centrin | Random |
| ENSG00000127445 | ENST00000247970 | PIN1 | Diff. RPE1 GFP-Centrin | Random |
| ENSG00000254093 | ENST00000519088 | PINX1 | Diff. RPE1 GFP-Centrin | Random |
| ENSG00000110697 | ENST00000534749 | PITPNM1 | Diff. RPE1 GFP-Centrin | Random |
| ENSG00000118762 | ENST00000237596 | PKD2 | Diff. RPE1 GFP-Centrin | Random |
| ENSG00000067225 | ENST00000335181 | PKM | Diff. RPE1 GFP-Centrin | Random |
| ENSG00000145632 | ENST00000274289 | PLK2 | Diff. RPE1 GFP-Centrin | Random |
| ENSG00000173846 | ENST00000372201 | PLK3 | Diff. RPE1 GFP-Centrin | Random |
| ENSG00000164087 | ENST00000296484 | POC1A | Diff. RPE1 GFP-Centrin | Random |
| ENSG00000139323 | ENST00000313546 | POC1B | Diff. RPE1 GFP-Centrin | Random |
| ENSG00000152359 | ENST00000428202 | POC5 | Diff. RPE1 GFP-Centrin | Random |
| ENSG00000070501 | ENST00000265421 | POLB | Diff. RPE1 GFP-Centrin | Random |
| ENSG00000100413 | ENST00000355209 | POLR3H | Diff. RPE1 GFP-Centrin | Random |
| ENSG00000066027 | ENST00000261461 | PPP2R5A | Diff. RPE1 GFP-Centrin | Random |
| ENSG00000163605 | ENST00000356692 | PPP4R2 | Diff. RPE1 GFP-Centrin | Random |
| ENSG00000198901 | ENST00000556972 | PRC1 | Diff. RPE1 GFP-Centrin | Random |
| ENSG00000142875 | ENST00000370689 | PRKACB | Diff. RPE1 GFP-Centrin | Random |
| ENSG00000114302 | ENST00000265563 | PRKAR2A | Diff. RPE1 GFP-Centrin | Random |
| ENSG00000005249 | ENST00000265717 | PRKAR2B | Diff. RPE1 GFP-Centrin | Random |
| ENSG00000101000 | ENST00000216968 | PROCR | Diff. RPE1 GFP-Centrin | Random |
| ENSG00000080815 | ENST00000357710 | PSEN1 | Diff. RPE1 GFP-Centrin | Random |
| ENSG00000143801 | ENST00000366783 | PSEN2 | Diff. RPE1 GFP-Centrin | Random |
| ENSG00000129084 | ENST00000396393 | PSMA1 | Diff. RPE1 GFP-Centrin | Random |
| ENSG00000100804 | ENST00000361611 | PSMB5 | Diff. RPE1 GFP-Centrin | Random |
| ENSG00000134222 | ENST00000369909 | PSRC1 | Diff. RPE1 GFP-Centrin | Random |
| ENSG00000185920 | ENST00000546744 | PTCH1 | Diff. RPE1 GFP-Centrin | Random |
| ENSG00000076201 | ENST00000265562 | PTPN23 | Diff. RPE1 GFP-Centrin | Random |
| ENSG00000084733 | ENST00000264710 | RAB10 | Diff. RPE1 GFP-Centrin | Random |
| ENSG00000103769 | ENST00000564910 | RAB11A | Diff. RPE1 GFP-Centrin | Random |
| ENSG00000112210 | ENST00000317483 | RAB23 | Diff. RPE1 GFP-Centrin | Random |
| ENSG00000222014 | ENST00000410061 | RAB6C | Diff. RPE1 GFP-Centrin | Random |
| ENSG00000167461 | ENST00000300935 | RAB8A | Diff. RPE1 GFP-Centrin | Random |
| ENSG00000011454 | ENST00000373647 | RABGAP1 | Diff. RPE1 GFP-Centrin | Random |
| ENSG00000011454 | ENST00000373647 | RABGAP1 | Diff. RPE1 GFP-Centrin | Random |
| ENSG00000152061 | ENST00000251507 | RABGAP1L | Diff. RPE1 GFP-Centrin | Random |
| ENSG00000161800 | ENST00000312377 | RACGAP1 | Diff. RPE1 GFP-Centrin | Random |
| ENSG00000204628 | ENST00000512805 | RACK1 | Diff. RPE1 GFP-Centrin | Random |
| ENSG00000051180 | ENST00000267868 | RAD51 | Diff. RPE1 GFP-Centrin | Random |
| ENSG00000185379 | ENST00000335858 | RAD51D | Diff. RPE1 GFP-Centrin | Random |
| ENSG00000132341 | ENST00000254675 | RAN | Diff. RPE1 GFP-Centrin | Random |
| ENSG00000099901 | ENST00000331821 | RANBP1 | Diff. RPE1 GFP-Centrin | Random |
| ENSG00000153201 | ENST00000283195 | RANBP2 | Diff. RPE1 GFP-Centrin | Random |
| ENSG00000031823 | ENST00000439268 | RANBP3 | Diff. RPE1 GFP-Centrin | Random |
| ENSG00000100401 | ENST00000356244 | RANGAP1 | Diff. RPE1 GFP-Centrin | Random |
| ENSG00000125249 | ENST00000245304 | RAP2A | Diff. RPE1 GFP-Centrin | Random |
| ENSG00000158987 | ENST00000509018 | RAPGEF6 | Diff. RPE1 GFP-Centrin | Random |
| ENSG00000179051 | ENST00000375436 | RCC2 | Diff. RPE1 GFP-Centrin | Random |

| | | | | |
|---|---|---|---|---|
| ENSG00000137710 | ENST00000343115 | RDX | Diff. RPE1 GFP-Centrin | Random |
| ENSG00000165476 | ENST00000373758 | REEP3 | Diff. RPE1 GFP-Centrin | Random |
| ENSG00000168476 | ENST00000306306 | REEP4 | Diff. RPE1 GFP-Centrin | Random |
| ENSG00000104856 | ENST00000221452 | RELB | Diff. RPE1 GFP-Centrin | Random |
| ENSG00000196862 | ENST00000408999 | RGPD4 | Diff. RPE1 GFP-Centrin | Random |
| ENSG00000111785 | ENST00000392837 | RIC8B | Diff. RPE1 GFP-Centrin | Random |
| ENSG00000080345 | ENST00000243326 | RIF1 | Diff. RPE1 GFP-Centrin | Random |
| ENSG00000188026 | ENST00000376874 | RILPL1 | Diff. RPE1 GFP-Centrin | Random |
| ENSG00000139405 | ENST00000548278 | RITA1 | Diff. RPE1 GFP-Centrin | Random |
| ENSG00000176623 | ENST00000406452 | RMDN1 | Diff. RPE1 GFP-Centrin | Random |
| ENSG00000137824 | ENST00000338376 | RMDN3 | Diff. RPE1 GFP-Centrin | Random |
| ENSG00000112130 | ENST00000229866 | RNF8 | Diff. RPE1 GFP-Centrin | Random |
| ENSG00000134318 | ENST00000315872 | ROCK2 | Diff. RPE1 GFP-Centrin | Random |
| ENSG00000101413 | ENST00000373433 | RPRD1B | Diff. RPE1 GFP-Centrin | Random |
| ENSG00000149273 | ENST00000524851 | RPS3 | Diff. RPE1 GFP-Centrin | Random |
| ENSG00000171863 | ENST00000304921 | RPS7 | Diff. RPE1 GFP-Centrin | Random |
| ENSG00000172426 | ENST00000372163 | RSPH9 | Diff. RPE1 GFP-Centrin | Random |
| ENSG00000087302 | ENST00000261700 | RTRAF | Diff. RPE1 GFP-Centrin | Random |
| ENSG00000183207 | ENST00000596247 | RUVBL2 | Diff. RPE1 GFP-Centrin | Random |
| ENSG00000156876 | ENST00000287482 | SASS6 | Diff. RPE1 GFP-Centrin | Random |
| ENSG00000143653 | ENST00000366510 | SCCPDH | Diff. RPE1 GFP-Centrin | Random |
| ENSG00000151466 | ENST00000281142 | SCLT1 | Diff. RPE1 GFP-Centrin | Random |
| ENSG00000111319 | ENST00000228916 | SCNN1A | Diff. RPE1 GFP-Centrin | Random |
| ENSG00000157020 | ENST00000350697 | SEC13 | Diff. RPE1 GFP-Centrin | Random |
| ENSG00000085415 | ENST00000262124 | SEH1L | Diff. RPE1 GFP-Centrin | Random |
| ENSG00000129810 | ENST00000442720 | SGO1 | Diff. RPE1 GFP-Centrin | Random |
| ENSG00000138771 | ENST00000296043 | SHROOM3 | Diff. RPE1 GFP-Centrin | Random |
| ENSG00000068903 | ENST00000249396 | SIRT2 | Diff. RPE1 GFP-Centrin | Random |
| ENSG00000182628 | ENST00000330137 | SKA2 | Diff. RPE1 GFP-Centrin | Random |
| ENSG00000157933 | ENST00000378536 | SKI | Diff. RPE1 GFP-Centrin | Random |
| ENSG00000155380 | ENST00000369626 | SLC16A1 | Diff. RPE1 GFP-Centrin | Random |
| ENSG00000133302 | ENST00000265140 | SLF1 | Diff. RPE1 GFP-Centrin | Random |
| ENSG00000141646 | ENST00000342988 | SMAD4 | Diff. RPE1 GFP-Centrin | Random |
| ENSG00000101665 | ENST00000262158 | SMAD7 | Diff. RPE1 GFP-Centrin | Random |
| ENSG00000153147 | ENST00000283131 | SMARCA5 | Diff. RPE1 GFP-Centrin | Random |
| ENSG00000128602 | ENST00000249373 | SMO | Diff. RPE1 GFP-Centrin | Random |
| ENSG00000099940 | ENST00000215730 | SNAP29 | Diff. RPE1 GFP-Centrin | Random |
| ENSG00000064199 | ENST00000227135 | SPA17 | Diff. RPE1 GFP-Centrin | Random |
| ENSG00000144451 | ENST00000432529 | SPAG16 | Diff. RPE1 GFP-Centrin | Random |
| ENSG00000061656 | ENST00000374273 | SPAG4 | Diff. RPE1 GFP-Centrin | Random |
| ENSG00000076382 | ENST00000321765 | SPAG5 | Diff. RPE1 GFP-Centrin | Random |
| ENSG00000133104 | ENST00000494062 | SPART | Diff. RPE1 GFP-Centrin | Random |
| ENSG00000021574 | ENST00000615843 | SPAST | Diff. RPE1 GFP-Centrin | Random |
| ENSG00000163611 | ENST00000295872 | SPICE1 | Diff. RPE1 GFP-Centrin | Random |
| ENSG00000198917 | ENST00000361256 | SPOUT1 | Diff. RPE1 GFP-Centrin | Random |
| ENSG00000084112 | ENST00000360239 | SSH1 | Diff. RPE1 GFP-Centrin | Random |
| ENSG00000176101 | ENST00000322310 | SSNA1 | Diff. RPE1 GFP-Centrin | Random |
| ENSG00000159433 | ENST00000290607 | STARD9 | Diff. RPE1 GFP-Centrin | Random |
| ENSG00000040341 | ENST00000355780 | STAU2 | Diff. RPE1 GFP-Centrin | Random |
| ENSG00000117632 | ENST00000374291 | STMN1 | Diff. RPE1 GFP-Centrin | Random |
| ENSG00000159082 | ENST00000438952 | SYNJ1 | Diff. RPE1 GFP-Centrin | Random |
| ENSG00000147526 | ENST00000348567 | TACC1 | Diff. RPE1 GFP-Centrin | Random |
| ENSG00000171148 | ENST00000301964 | TADA3 | Diff. RPE1 GFP-Centrin | Random |

| | | | | |
|---|---|---|---|---|
| ENSG00000169762 | ENST00000405303 | TAPT1 | Diff. RPE1 GFP-Centrin | Random |
| ENSG00000127364 | ENST00000247881 | TAS2R4 | Diff. RPE1 GFP-Centrin | Random |
| ENSG00000156787 | ENST00000518805 | TBC1D31 | Diff. RPE1 GFP-Centrin | Random |
| ENSG00000146350 | ENST00000398197 | TBC1D32 | Diff. RPE1 GFP-Centrin | Random |
| ENSG00000141556 | ENST00000355528 | TBCD | Diff. RPE1 GFP-Centrin | Random |
| ENSG00000284770 | ENST00000366601 | TBCE | Diff. RPE1 GFP-Centrin | Random |
| ENSG00000139437 | ENST00000405876 | TCHP | Diff. RPE1 GFP-Centrin | Random |
| ENSG00000120438 | ENST00000321394 | TCP1 | Diff. RPE1 GFP-Centrin | Random |
| ENSG00000179029 | ENST00000437139 | TMEM107 | Diff. RPE1 GFP-Centrin | Random |
| ENSG00000186889 | ENST00000335390 | TMEM17 | Diff. RPE1 GFP-Centrin | Random |
| ENSG00000150433 | ENST00000531262 | TMEM218 | Diff. RPE1 GFP-Centrin | Random |
| ENSG00000155755 | ENST00000286196 | TMEM237 | Diff. RPE1 GFP-Centrin | Random |
| ENSG00000164953 | ENST00000453321 | TMEM67 | Diff. RPE1 GFP-Centrin | Random |
| ENSG00000173273 | ENST00000310430 | TNKS | Diff. RPE1 GFP-Centrin | Random |
| ENSG00000107854 | ENST00000371627 | TNKS2 | Diff. RPE1 GFP-Centrin | Random |
| ENSG00000083312 | ENST00000454282 | TNPO1 | Diff. RPE1 GFP-Centrin | Random |
| ENSG00000198718 | ENST00000361462 | TOGARAM1 | Diff. RPE1 GFP-Centrin | Random |
| ENSG00000197579 | ENST00000360538 | TOPORS | Diff. RPE1 GFP-Centrin | Random |
| ENSG00000134779 | ENST00000383056 | TPGS2 | Diff. RPE1 GFP-Centrin | Random |
| ENSG00000204104 | ENST00000391993 | TRAF3IP1 | Diff. RPE1 GFP-Centrin | Random |
| ENSG00000213186 | ENST00000309784 | TRIM59 | Diff. RPE1 GFP-Centrin | Random |
| ENSG00000100106 | ENST00000403663 | TRIOBP | Diff. RPE1 GFP-Centrin | Random |
| ENSG00000100815 | ENST00000267622 | TRIP11 | Diff. RPE1 GFP-Centrin | Random |
| ENSG00000103671 | ENST00000261884 | TRIP4 | Diff. RPE1 GFP-Centrin | Random |
| ENSG00000128881 | ENST00000267890 | TTBK2 | Diff. RPE1 GFP-Centrin | Random |
| ENSG00000149292 | ENST00000529221 | TTC12 | Diff. RPE1 GFP-Centrin | Random |
| ENSG00000011295 | ENST00000475723 | TTC19 | Diff. RPE1 GFP-Centrin | Random |
| ENSG00000123607 | ENST00000243344 | TTC21B | Diff. RPE1 GFP-Centrin | Random |
| ENSG00000105948 | ENST00000464848 | TTC26 | Diff. RPE1 GFP-Centrin | Random |
| ENSG00000100154 | ENST00000397906 | TTC28 | Diff. RPE1 GFP-Centrin | Random |
| ENSG00000197557 | ENST00000355689 | TTC30A | Diff. RPE1 GFP-Centrin | Random |
| ENSG00000165533 | ENST00000345383 | TTC8 | Diff. RPE1 GFP-Centrin | Random |
| ENSG00000135912 | ENST00000392102 | TTLL4 | Diff. RPE1 GFP-Centrin | Random |
| ENSG00000119685 | ENST00000298832 | TTLL5 | Diff. RPE1 GFP-Centrin | Random |
| ENSG00000123416 | ENST00000336023 | TUBA1B | Diff. RPE1 GFP-Centrin | Random |
| ENSG00000127824 | ENST00000248437 | TUBA4A | Diff. RPE1 GFP-Centrin | Random |
| ENSG00000137267 | ENST00000333628 | TUBB2A | Diff. RPE1 GFP-Centrin | Random |
| ENSG00000258947 | ENST00000315491 | TUBB3 | Diff. RPE1 GFP-Centrin | Random |
| ENSG00000188229 | ENST00000340384 | TUBB4B | Diff. RPE1 GFP-Centrin | Random |
| ENSG00000176014 | ENST00000317702 | TUBB6 | Diff. RPE1 GFP-Centrin | Random |
| ENSG00000074935 | ENST00000368662 | TUBE1 | Diff. RPE1 GFP-Centrin | Random |
| ENSG00000137822 | ENST00000564079 | TUBGCP4 | Diff. RPE1 GFP-Centrin | Random |
| ENSG00000112041 | ENST00000322263 | TULP1 | Diff. RPE1 GFP-Centrin | Random |
| ENSG00000078246 | ENST00000448120 | TULP3 | Diff. RPE1 GFP-Centrin | Random |
| ENSG00000130338 | ENST00000367097 | TULP4 | Diff. RPE1 GFP-Centrin | Random |
| ENSG00000115514 | ENST00000264255 | TXNDC9 | Diff. RPE1 GFP-Centrin | Random |
| ENSG00000127481 | ENST00000375254 | UBR4 | Diff. RPE1 GFP-Centrin | Random |
| ENSG00000109103 | ENST00000335765 | UNC119 | Diff. RPE1 GFP-Centrin | Random |
| ENSG00000175970 | ENST00000344651 | UNC119B | Diff. RPE1 GFP-Centrin | Random |
| ENSG00000135763 | ENST00000258243 | URB2 | Diff. RPE1 GFP-Centrin | Random |
| ENSG00000124486 | ENST00000324545 | USP9X | Diff. RPE1 GFP-Centrin | Random |
| ENSG00000198382 | ENST00000356136 | UVRAG | Diff. RPE1 GFP-Centrin | Random |
| ENSG00000126756 | ENST00000333119 | UXT | Diff. RPE1 GFP-Centrin | Random |

| | | | | |
|---|---|---|---|---|
| ENSG00000139722 | ENST00000267202 | VPS37B | Diff. RPE1 GFP-Centrin | Random |
| ENSG00000132612 | ENST00000254950 | VPS4A | Diff. RPE1 GFP-Centrin | Random |
| ENSG00000119541 | ENST00000238497 | VPS4B | Diff. RPE1 GFP-Centrin | Random |
| ENSG00000143951 | ENST00000409354 | WDPCP | Diff. RPE1 GFP-Centrin | Random |
| ENSG00000119333 | ENST00000372715 | WDR34 | Diff. RPE1 GFP-Centrin | Random |
| ENSG00000118965 | ENST00000281405 | WDR35 | Diff. RPE1 GFP-Centrin | Random |
| ENSG00000085433 | ENST00000369962 | WDR47 | Diff. RPE1 GFP-Centrin | Random |
| ENSG00000126870 | ENST00000407559 | WDR60 | Diff. RPE1 GFP-Centrin | Random |
| ENSG00000075702 | ENST00000401500 | WDR62 | Diff. RPE1 GFP-Centrin | Random |
| ENSG00000095397 | ENST00000265134 | WHRN | Diff. RPE1 GFP-Centrin | Random |
| ENSG00000011451 | ENST00000599686 | WIZ | Diff. RPE1 GFP-Centrin | Random |
| ENSG00000116213 | ENST00000270708 | WRAP73 | Diff. RPE1 GFP-Centrin | Random |
| ENSG00000165392 | ENST00000298139 | WRN | Diff. RPE1 GFP-Centrin | Random |
| ENSG00000196584 | ENST00000359321 | XRCC2 | Diff. RPE1 GFP-Centrin | Random |
| ENSG00000164924 | ENST00000395958 | YWHAZ | Diff. RPE1 GFP-Centrin | Random |
| ENSG00000214717 | ENST00000381223 | ZBED1 | Diff. RPE1 GFP-Centrin | Random |
| ENSG00000166140 | ENST00000561768 | ZFYVE19 | Diff. RPE1 GFP-Centrin | Random |
| ENSG00000072121 | ENST00000347230 | ZFYVE26 | Diff. RPE1 GFP-Centrin | Random |
| ENSG00000164631 | ENST00000405858 | ZNF12 | Diff. RPE1 GFP-Centrin | Random |
| ENSG00000010244 | ENST00000394673 | ZNF207 | Diff. RPE1 GFP-Centrin | Random |
| ENSG00000181315 | ENST00000415922 | ZNF322 | Diff. RPE1 GFP-Centrin | Random |
| ENSG00000109445 | ENST00000262990 | ZNF330 | Diff. RPE1 GFP-Centrin | Random |
| ENSG00000138311 | ENST00000395254 | ZNF365 | Diff. RPE1 GFP-Centrin | Random |

Supplementary table S2

**Oligonucleotide sequence (5' end on the left, 3' end on the right ; concatenation of first barcode (25 nucleotides), FLAP Y sequence : TTACACTCGGACCTCGTCGACATGCATT, target-specific hybridization sequence, FLAP X sequence : CCTCCTAAGTTTCGAGCTGGACTCAGTG, second barcode (25 nucleotides))**

**BICD2**

CGAGGGGTACGATCTCCCTGTTCCATTACACTCGGACCTCGTCGACATGCATTACAATGACTGTTTTCTTTCCCTCAATGAAGATGCCCTCCTAAGTTTCGAGCTGGACTCAGTGGGGGTTTGGTTTACTCCTGGTTACC
CGAGGGGTACGATCTCCCTGTTCCATTACACTCGGACCTCGTCGACATGCATTGTGTAAAGGTTCCTTGGCAGGAGAACATGCCCTCCTAAGTTTCGAGCTGGACTCAGTGGGGGTTTGGTTTACTCCTGGTTACC
CGAGGGGTACGATCTCCCTGTTCCATTACACTCGGACCTCGTCGACATGCATTGTCCTCCTGGCAGCTACTCTACAGGCTCCTCCTAAGTTTCGAGCTGGACTCAGTGGGGGTTTGGTTTACTCCTGGTTACC
CGAGGGGTACGATCTCCCTGTTCCATTACACTCGGACCTCGTCGACATGCATTCGTTGATGTCCACCTCGTAGTAGTCCCCCCTCCTAAGTTTCGAGCTGGACTCAGTGGGGGTTTGGTTTACTCCTGGTTACC
CGAGGGGTACGATCTCCCTGTTCCATTACACTCGGACCTCGTCGACATGCATTGAGCAAAGGTCAAGTTTTCTTCTTGTTACATTGAACCCTCCTAAGTTTCGAGCTGGACTCAGTGGGGGTTTGGTTTACTCCTGGTTACC
CGAGGGGTACGATCTCCCTGTTCCATTACACTCGGACCTCGTCGACATGCATTTAGACCACGGCACCTGAGACCGTCTCTCCTCCTAAGTTTCGAGCTGGACTCAGTGGGGGTTTGGTTTACTCCTGGTTACC
CGAGGGGTACGATCTCCCTGTTCCATTACACTCGGACCTCGTCGACATGCATTGAACTTGAGGCCATCCAGCGAGACATGCCCTCCTAAGTTTCGAGCTGGACTCAGTGGGGGTTTGGTTTACTCCTGGTTACC
CGAGGGGTACGATCTCCCTGTTCCATTACACTCGGACCTCGTCGACATGCATTTGTAACAAGGAGCCCCCGTGGTGGGCCTCCTAAGTTTCGAGCTGGACTCAGTGGGGGTTTGGTTTACTCCTGGTTACC
CGAGGGGTACGATCTCCCTGTTCCATTACACTCGGACCTCGTCGACATGCATTCAAACATAGCACGCAGCGAGGAGAAGGTGCCTCCTAAGTTTCGAGCTGGACTCAGTGGGGGTTTGGTTTACTCCTGGTTACC
CGAGGGGTACGATCTCCCTGTTCCATTACACTCGGACCTCGTCGACATGCATTTTTTTCCCGCTCCATCTGCATCAGCTGCTCCTCCTAAGTTTCGAGCTGGACTCAGTGGGGGTTTGGTTTACTCCTGGTTACC
CGAGGGGTACGATCTCCCTGTTCCATTACACTCGGACCTCGTCGACATGCATTTTGATCTGGTCACGGATGATAGCGATCAGGCCTCCTAAGTTTCGAGCTGGACTCAGTGGGGGTTTGGTTTACTCCTGGTTACC
CGAGGGGTACGATCTCCCTGTTCCATTACACTCGGACCTCGTCGACATGCATTCCATACAGAGGCCTTTCCACGCAGCAGCCTCCTAAGTTTCGAGCTGGACTCAGTGGGGGTTTGGTTTACTCCTGGTTACC
CGAGGGGTACGATCTCCCTGTTCCATTACACTCGGACCTCGTCGACATGCATTCACTGAGTAGGTCGGAGACGAGGCTGGCCTCCTAAGTTTCGAGCTGGACTCAGTGGGGGTTTGGTTTACTCCTGGTTACC
CGAGGGGTACGATCTCCCTGTTCCATTACACTCGGACCTCGTCGACATGCATTGTCACTCAGGGGTGATGGCAGTGAGGCCTCCTAAGTTTCGAGCTGGACTCAGTGGGGGTTTGGTTTACTCCTGGTTACC
CGAGGGGTACGATCTCCCTGTTCCATTACACTCGGACCTCGTCGACATGCATTTGTAGATGTTCATGGGCTCCCGGCGTGCCTCCTAAGTTTCGAGCTGGACTCAGTGGGGGTTTGGTTTACTCCTGGTTACC
CGAGGGGTACGATCTCCCTGTTCCATTACACTCGGACCTCGTCGACATGCATTTCCTGATACCCTCAGCTCTTCACAAACGTGCCTCCTAAGTTTCGAGCTGGACTCAGTGGGGGTTTGGTTTACTCCTGGTTACC
CGAGGGGTACGATCTCCCTGTTCCATTACACTCGGACCTCGTCGACATGCATTATCCTCCAGCTGGCTGTTGAGGTACTCGCCTCCTAAGTTTCGAGCTGGACTCAGTGGGGGTTTGGTTTACTCCTGGTTACC
CGAGGGGTACGATCTCCCTGTTCCATTACACTCGGACCTCGTCGACATGCATTCTCAGATTCTCTGTGAGGCGGGTCACCCCTCCTAAGTTTCGAGCTGGACTCAGTGGGGGTTTGGTTTACTCCTGGTTACC
CGAGGGGTACGATCTCCCTGTTCCATTACACTCGGACCTCGTCGACATGCATTTCGCAGCATGCTCAACATTAAAGCTTTTTTCCTCCTCCTAAGTTTCGAGCTGGACTCAGTGGGGGTTTGGTTTACTCCTGGTTACC
CGAGGGGTACGATCTCCCTGTTCCATTACACTCGGACCTCGTCGACATGCATTTAGTCCAGCATGACACGGTTGGGTGTCTCCTCCTAAGTTTCGAGCTGGACTCAGTGGGGGTTTGGTTTACTCCTGGTTACC
CGAGGGGTACGATCTCCCTGTTCCATTACACTCGGACCTCGTCGACATGCATTTTGAGGCCCTCAAACTCCACCTGGTTCTCCTCCTAAGTTTCGAGCTGGACTCAGTGGGGGTTTGGTTTACTCCTGGTTACC
CGAGGGGTACGATCTCCCTGTTCCATTACACTCGGACCTCGTCGACATGCATTATTGTTGCACATGCACACGTGGTGGTAGAGCCTCCTAAGTTTCGAGCTGGACTCAGTGGGGGTTTGGTTTACTCCTGGTTACC
CGAGGGGTACGATCTCCCTGTTCCATTACACTCGGACCTCGTCGACATGCATTATGGTACTTGCAGGCCAAGATCTCAGGCCCTCCTAAGTTTCGAGCTGGACTCAGTGGGGGTTTGGTTTACTCCTGGTTACC
CGAGGGGTACGATCTCCCTGTTCCATTACACTCGGACCTCGTCGACATGCATTTCTCCTCCTCCAGACGCTTGATCTCATGCCTCCTAAGTTTCGAGCTGGACTCAGTGGGGGTTTGGTTTACTCCTGGTTACC
CGAGGGGTACGATCTCCCTGTTCCATTACACTCGGACCTCGTCGACATGCATTGTTTTTCTTCCTCCAAGAGTACCAATTTGACCACCTCCTAAGTTTCGAGCTGGACTCAGTGGGGGTTTGGTTTACTCCTGGTTACC
CGAGGGGTACGATCTCCCTGTTCCATTACACTCGGACCTCGTCGACATGCATTTATGGAGCACACAGCTCTGCCTCGTGCCCTCCTAAGTTTCGAGCTGGACTCAGTGGGGGTTTGGTTTACTCCTGGTTACC
CGAGGGGTACGATCTCCCTGTTCCATTACACTCGGACCTCGTCGACATGCATTTGCTTCACAACGTCCTAAAACCCAGAAAATTCTCCTCCTAAGTTTCGAGCTGGACTCAGTGGGGGTTTGGTTTACTCCTGGTTACC
CGAGGGGTACGATCTCCCTGTTCCATTACACTCGGACCTCGTCGACATGCATTCTTCAGCTTCTGGATCTCAGAGATGTTGAGCCCTCCTAAGTTTCGAGCTGGACTCAGTGGGGGTTTGGTTTACTCCTGGTTACC
CGAGGGGTACGATCTCCCTGTTCCATTACACTCGGACCTCGTCGACATGCATTCAAAGCCATTGACCAGGGCCTCGGCACCTCCTAAGTTTCGAGCTGGACTCAGTGGGGGTTTGGTTTACTCCTGGTTACC
CGAGGGGTACGATCTCCCTGTTCCATTACACTCGGACCTCGTCGACATGCATTCTGGCCTTCTCTAGCAGGGAGACCTTCCCTCCTAAGTTTCGAGCTGGACTCAGTGGGGGTTTGGTTTACTCCTGGTTACC
CGAGGGGTACGATCTCCCTGTTCCATTACACTCGGACCTCGTCGACATGCATTTCCTCATGGCTGTCACGGTCCTTCTCGCCTCCTAAGTTTCGAGCTGGACTCAGTGGGGGTTTGGTTTACTCCTGGTTACC
CGAGGGGTACGATCTCCCTGTTCCATTACACTCGGACCTCGTCGACATGCATTTGTCCAGGGCTGTCTGCCGCTCCTTCCTCCTAAGTTTCGAGCTGGACTCAGTGGGGGTTTGGTTTACTCCTGGTTACC
CGAGGGGTACGATCTCCCTGTTCCATTACACTCGGACCTCGTCGACATGCATTTTGGCCAGCTCCTCACTGAAGGTCACCCCTCCTAAGTTTCGAGCTGGACTCAGTGGGGGTTTGGTTTACTCCTGGTTACC
CGAGGGGTACGATCTCCCTGTTCCATTACACTCGGACCTCGTCGACATGCATTTCCTATTCCTTTTCCATGCTCTGCCTCATTTTCCTCCTAAGTTTCGAGCTGGACTCAGTGGGGGTTTGGTTTACTCCTGGTTACC
CGAGGGGTACGATCTCCCTGTTCCATTACACTCGGACCTCGTCGACATGCATTTCTCCTGCTGTTCTGACAGGGAGCCCCCTCCTAAGTTTCGAGCTGGACTCAGTGGGGGTTTGGTTTACTCCTGGTTACC
CGAGGGGTACGATCTCCCTGTTCCATTACACTCGGACCTCGTCGACATGCATTCACCTCGAGCTCCTCGAACTGCAGCTTCCTCCTAAGTTTCGAGCTGGACTCAGTGGGGGTTTGGTTTACTCCTGGTTACC
CGAGGGGTACGATCTCCCTGTTCCATTACACTCGGACCTCGTCGACATGCATTATGTAGACCTGGTTTGCTAAGGAGTTTTAATGAGCCTCCTAAGTTTCGAGCTGGACTCAGTGGGGGTTTGGTTTACTCCTGGTTACC
CGAGGGGTACGATCTCCCTGTTCCATTACACTCGGACCTCGTCGACATGCATTAAAGAATACTATGGCTAACCCTCATCCCCTACCCTCCTAAGTTTCGAGCTGGACTCAGTGGGGGTTTGGTTTACTCCTGGTTACC
CGAGGGGTACGATCTCCCTGTTCCATTACACTCGGACCTCGTCGACATGCATTGACACCAGACAAACACGGTGGGAGCTGCCTCCTAAGTTTCGAGCTGGACTCAGTGGGGGTTTGGTTTACTCCTGGTTACC
CGAGGGGTACGATCTCCCTGTTCCATTACACTCGGACCTCGTCGACATGCATTTACTCCTTGATGTCATCCCGCAGGCGGCCTCCTAAGTTTCGAGCTGGACTCAGTGGGGGTTTGGTTTACTCCTGGTTACC
CGAGGGGTACGATCTCCCTGTTCCATTACACTCGGACCTCGTCGACATGCATTTCCATAAGCGCTTCCTTGTCCTTGTCCACCCTCCTAAGTTTCGAGCTGGACTCAGTGGGGGTTTGGTTTACTCCTGGTTACC
CGAGGGGTACGATCTCCCTGTTCCATTACACTCGGACCTCGTCGACATGCATTGCTGTGTAATGTACTCGTCACACCTGGTGCCTCCTAAGTTTCGAGCTGGACTCAGTGGGGGTTTGGTTTACTCCTGGTTACC
CGAGGGGTACGATCTCCCTGTTCCATTACACTCGGACCTCGTCGACATGCATTCGTTGTTGGGCTCGGCAGCATCGTCACCCTCCTAAGTTTCGAGCTGGACTCAGTGGGGGTTTGGTTTACTCCTGGTTACC
CGAGGGGTACGATCTCCCTGTTCCATTACACTCGGACCTCGTCGACATGCATTTCACCTTCTTTAGCTCCTTCTCCAGCCGCCTCCTAAGTTTCGAGCTGGACTCAGTGGGGGTTTGGTTTACTCCTGGTTACC
CGAGGGGTACGATCTCCCTGTTCCATTACACTCGGACCTCGTCGACATGCATTGCTCATCCTGGGCCACACTCAGGCTCCTCCTAAGTTTCGAGCTGGACTCAGTGGGGGTTTGGTTTACTCCTGGTTACC
CGAGGGGTACGATCTCCCTGTTCCATTACACTCGGACCTCGTCGACATGCATTAAAATAGGAGTCCCTACAACTAGGAATAAGAAATGCCCTCCTAAGTTTCGAGCTGGACTCAGTGGGGGTTTGGTTTACTCCTGGTTACC
CGAGGGGTACGATCTCCCTGTTCCATTACACTCGGACCTCGTCGACATGCATTTGCTTGTCCAAAGGCCTCCTTGAGCTGCCTCCTAAGTTTCGAGCTGGACTCAGTGGGGGTTTGGTTTACTCCTGGTTACC
CGAGGGGTACGATCTCCCTGTTCCATTACACTCGGACCTCGTCGACATGCATTCGTTCATACAGCTTGCTCAGCTTGTCCCACCTCCTAAGTTTCGAGCTGGACTCAGTGGGGGTTTGGTTTACTCCTGGTTACC
CGAGGGGTACGATCTCCCTGTTCCATTACACTCGGACCTCGTCGACATGCATTGGTTTTGTGTTTGCTTTCATTTTCTAAGCCCCTCCTCCTAAGTTTCGAGCTGGACTCAGTGGGGGTTTGGTTTACTCCTGGTTACC
CGAGGGGTACGATCTCCCTGTTCCATTACACTCGGACCTCGTCGACATGCATTTGAGCAGCGACTTCAGCTTGAGGATCTCCCTCCTAAGTTTCGAGCTGGACTCAGTGGGGGTTTGGTTTACTCCTGGTTACC
CGAGGGGTACGATCTCCCTGTTCCATTACACTCGGACCTCGTCGACATGCATTAAAGCACTCATCTCCGAGTCTAAAGCTACACGCCTCCTAAGTTTCGAGCTGGACTCAGTGGGGGTTTGGTTTACTCCTGGTTACC
CGAGGGGTACGATCTCCCTGTTCCATTACACTCGGACCTCGTCGACATGCATTGCGTGTGCTCCAGCTGCTTCTGTGTGTCCTCCTAAGTTTCGAGCTGGACTCAGTGGGGGTTTGGTTTACTCCTGGTTACC
CGAGGGGTACGATCTCCCTGTTCCATTACACTCGGACCTCGTCGACATGCATTATCTGAATACCCTGGATTTCTTAATGAACATGGCCCTCCTAAGTTTCGAGCTGGACTCAGTGGGGGTTTGGTTTACTCCTGGTTACC
CGAGGGGTACGATCTCCCTGTTCCATTACACTCGGACCTCGTCGACATGCATTATGTGACTCTTTACCCAGGCCCACTGTTCCCTCCTAAGTTTCGAGCTGGACTCAGTGGGGGTTTGGTTTACTCCTGGTTACC
CGAGGGGTACGATCTCCCTGTTCCATTACACTCGGACCTCGTCGACATGCATTTATGCGCACTGGCTTTGTAGGCATTCACATCCTCCTAAGTTTCGAGCTGGACTCAGTGGGGGTTTGGTTTACTCCTGGTTACC
CGAGGGGTACGATCTCCCTGTTCCATTACACTCGGACCTCGTCGACATGCATTTCCCACTAACCTCACTCAGCAAACAAAACAGCCTCCTAAGTTTCGAGCTGGACTCAGTGGGGGTTTGGTTTACTCCTGGTTACC
CGAGGGGTACGATCTCCCTGTTCCATTACACTCGGACCTCGTCGACATGCATTGGTGGCTGGTGTAGAAGGAGTCATTGATGCCTCCTAAGTTTCGAGCTGGACTCAGTGGGGGTTTGGTTTACTCCTGGTTACC
CGAGGGGTACGATCTCCCTGTTCCATTACACTCGGACCTCGTCGACATGCATTCATTGCGCAGCTTCATCATGGTCTCGGTAACCTCCTAAGTTTCGAGCTGGACTCAGTGGGGGTTTGGTTTACTCCTGGTTACC
CGAGGGGTACGATCTCCCTGTTCCATTACACTCGGACCTCGTCGACATGCATTGCAAGAGAAGATATAATGTCTCGCTTTCACATTTGCCTCCTAAGTTTCGAGCTGGACTCAGTGGGGGTTTGGTTTACTCCTGGTTACC
CGAGGGGTACGATCTCCCTGTTCCATTACACTCGGACCTCGTCGACATGCATTCATGCCCACTAGCTCTTCCCTGCTCGCCCTCCTAAGTTTCGAGCTGGACTCAGTGGGGGTTTGGTTTACTCCTGGTTACC
CGAGGGGTACGATCTCCCTGTTCCATTACACTCGGACCTCGTCGACATGCATTTCCATCTCGCTGCGGATAGCCTCATAGTCCTCCTAAGTTTCGAGCTGGACTCAGTGGGGGTTTGGTTTACTCCTGGTTACC
CGAGGGGTACGATCTCCCTGTTCCATTACACTCGGACCTCGTCGACATGCATTCCTGCAGCAGACGAGCTTCCCGGAATTTCCTCCTAAGTTTCGAGCTGGACTCAGTGGGGGTTTGGTTTACTCCTGGTTACC
CGAGGGGTACGATCTCCCTGTTCCATTACACTCGGACCTCGTCGACATGCATTGAGGACATTGCGCAACTGCTTCAGCTCCCCTCCTAAGTTTCGAGCTGGACTCAGTGGGGGTTTGGTTTACTCCTGGTTACC
CGAGGGGTACGATCTCCCTGTTCCATTACACTCGGACCTCGTCGACATGCATTGCAGTGGCATATCCAAAAAGTTTCTAAGTGGGCCTCCTAAGTTTCGAGCTGGACTCAGTGGGGGTTTGGTTTACTCCTGGTTACC
CGAGGGGTACGATCTCCCTGTTCCATTACACTCGGACCTCGTCGACATGCATTATGATCTAAAGGCTGCTGAACCACAGCGTGCCTCCTAAGTTTCGAGCTGGACTCAGTGGGGGTTTGGTTTACTCCTGGTTACC
CGAGGGGTACGATCTCCCTGTTCCATTACACTCGGACCTCGTCGACATGCATTTTTTTAGGTTGCCCTTCCTTCCTCCTTGTGGCCTCCTAAGTTTCGAGCTGGACTCAGTGGGGGTTTGGTTTACTCCTGGTTACC
CGAGGGGTACGATCTCCCTGTTCCATTACACTCGGACCTCGTCGACATGCATTTCTGAGCACAGACACTTGCTTCTGCAGGCCTCCTAAGTTTCGAGCTGGACTCAGTGGGGGTTTGGTTTACTCCTGGTTACC
CGAGGGGTACGATCTCCCTGTTCCATTACACTCGGACCTCGTCGACATGCATTTCACCTGCTGTCCACGCTGGCCTTCACCTCCTAAGTTTCGAGCTGGACTCAGTGGGGGTTTGGTTTACTCCTGGTTACC
CGAGGGGTACGATCTCCCTGTTCCATTACACTCGGACCTCGTCGACATGCATTGTTCAGCGTCTTCTTCTCGTCCTCAGCAGCCTCCTAAGTTTCGAGCTGGACTCAGTGGGGGTTTGGTTTACTCCTGGTTACC
CGAGGGGTACGATCTCCCTGTTCCATTACACTCGGACCTCGTCGACATGCATTGCTCTGAGATCTCCTTGAGGCGGATGGCCTCCTAAGTTTCGAGCTGGACTCAGTGGGGGTTTGGTTTACTCCTGGTTACC
CGAGGGGTACGATCTCCCTGTTCCATTACACTCGGACCTCGTCGACATGCATTACGACTGTAGAATTTTCTAAGTGTACCACAATTTGGCCTCCTAAGTTTCGAGCTGGACTCAGTGGGGGTTTGGTTTACTCCTGGTTACC
CGAGGGGTACGATCTCCCTGTTCCATTACACTCGGACCTCGTCGACATGCATTCACGCTGGATCTCCACATTCTGGTTGATCTCCTCCTAAGTTTCGAGCTGGACTCAGTGGGGGTTTGGTTTACTCCTGGTTACC
CGAGGGGTACGATCTCCCTGTTCCATTACACTCGGACCTCGTCGACATGCATTTTTTGAGAATGATCTGTGCTGGGACACCTCCCTCCTAAGTTTCGAGCTGGACTCAGTGGGGGTTTGGTTTACTCCTGGTTACC
CGAGGGGTACGATCTCCCTGTTCCATTACACTCGGACCTCGTCGACATGCATTCTTAACTGATGAAGGCACTACTGGCAATGATTACCTCCTAAGTTTCGAGCTGGACTCAGTGGGGGTTTGGTTTACTCCTGGTTACC
CGAGGGGTACGATCTCCCTGTTCCATTACACTCGGACCTCGTCGACATGCATTCTTGGGCGTGGAGGTCTTGTTGTCCAGTCCTCCTAAGTTTCGAGCTGGACTCAGTGGGGGTTTGGTTTACTCCTGGTTACC
CGAGGGGTACGATCTCCCTGTTCCATTACACTCGGACCTCGTCGACATGCATTGTACTGCTCCTTGGAGGCCGACTCCTCCTCCTAAGTTTCGAGCTGGACTCAGTGGGGGTTTGGTTTACTCCTGGTTACC
CGAGGGGTACGATCTCCCTGTTCCATTACACTCGGACCTCGTCGACATGCATTCTGCTCATGGTCCAGCTCGAGCAGCTCCTCCTAAGTTTCGAGCTGGACTCAGTGGGGGTTTGGTTTACTCCTGGTTACC
CGAGGGGTACGATCTCCCTGTTCCATTACACTCGGACCTCGTCGACATGCATTTCATGTAGTGTGACAGCTCCTTGCGCAGCCTCCTAAGTTTCGAGCTGGACTCAGTGGGGGTTTGGTTTACTCCTGGTTACC
CGAGGGGTACGATCTCCCTGTTCCATTACACTCGGACCTCGTCGACATGCATTGTCAGCAGCCACCTTCTTGTGGTTTGTGTCCTCCTAAGTTTCGAGCTGGACTCAGTGGGGGTTTGGTTTACTCCTGGTTACC
CGAGGGGTACGATCTCCCTGTTCCATTACACTCGGACCTCGTCGACATGCATTTGCTTGTTGGCCTTGAGCACAGTGCGCCCTCCTAAGTTTCGAGCTGGACTCAGTGGGGGTTTGGTTTACTCCTGGTTACC
CGAGGGGTACGATCTCCCTGTTCCATTACACTCGGACCTCGTCGACATGCATTAAGGTCCATTTGGTTCCCAAAGCACACTCACCTCCTAAGTTTCGAGCTGGACTCAGTGGGGGTTTGGTTTACTCCTGGTTACC
CGAGGGGTACGATCTCCCTGTTCCATTACACTCGGACCTCGTCGACATGCATTGGCGCTCATTCTCCGACTGCGTGTTGGCCTCCTAAGTTTCGAGCTGGACTCAGTGGGGGTTTGGTTTACTCCTGGTTACC
CGAGGGGTACGATCTCCCTGTTCCATTACACTCGGACCTCGTCGACATGCATTATACACTTAGCTGAGCTCCTCGCTGGGTCCTCCTAAGTTTCGAGCTGGACTCAGTGGGGGTTTGGTTTACTCCTGGTTACC
CGAGGGGTACGATCTCCCTGTTCCATTACACTCGGACCTCGTCGACATGCATTCTGTTCTTCTGTTCGCGCTCCGTCTTCAGCCTCCTAAGTTTCGAGCTGGACTCAGTGGGGGTTTGGTTTACTCCTGGTTACC
CGAGGGGTACGATCTCCCTGTTCCATTACACTCGGACCTCGTCGACATGCATTTGATGTTCTCCTCCTCCAGTTCCGAGTAGCCTCCTAAGTTTCGAGCTGGACTCAGTGGGGGTTTGGTTTACTCCTGGTTACC
CGAGGGGTACGATCTCCCTGTTCCATTACACTCGGACCTCGTCGACATGCATTCATGGCCTTCTCATTCTCATACTTGCTCTTCACCTCCTAAGTTTCGAGCTGGACTCAGTGGGGGTTTGGTTTACTCCTGGTTACC
CGAGGGGTACGATCTCCCTGTTCCATTACACTCGGACCTCGTCGACATGCATTTCTGCAGCTCTAGCACCTTCCGCACGTCCTCCTAAGTTTCGAGCTGGACTCAGTGGGGGTTTGGTTTACTCCTGGTTACC

**NIN**

ATCCTGGGTTCTGCCTACAGTCAGATTACACTCGGACCTCGTCGACATGCATTACCTCAGGTGCTGCATCTTCACACACTATCCTCCTAAGTTTCGAGCTGGACTCAGTGGGCGTTGAGTTGGTCCCTTCGTTAG
ATCCTGGGTTCTGCCTACAGTCAGATTACACTCGGACCTCGTCGACATGCATTTCATTTTCCAGACGACTGTTTTCCTTTAAAGAGAGCCTCCTAAGTTTCGAGCTGGACTCAGTGGGCGTTGAGTTGGTCCCTTCGTTAG

ATCCTGGGTTCTGCCTACAGTCAGATTACACTCGGACCTCGTCGACATGCATTGCGCTACTCTGAAGGCCTTCCCTCTCCCTCCTAAGTTTCGAGCTGGACTCAGTGGGCGTTGAGTTGGTCCCTTCGTTAG
ATCCTGGGTTCTGCCTACAGTCAGATTACACTCGGACCTCGTCGACATGCATTCGCTCAAAGCTTGCTTGAGCCTGTGTCAGCCTCCTAAGTTTCGAGCTGGACTCAGTGGGCGTTGAGTTGGTCCCTTCGTTAG
ATCCTGGGTTCTGCCTACAGTCAGATTACACTCGGACCTCGTCGACATGCATTCTAGCCTTGAGCTCCATCTCATGTTGCAGCCTCCTAAGTTTCGAGCTGGACTCAGTGGGCGTTGAGTTGGTCCCTTCGTTAG
ATCCTGGGTTCTGCCTACAGTCAGATTACACTCGGACCTCGTCGACATGCATTCATATTCTGCCAACTTCTCTGCATTTTCTAGAAGCCTCCTAAGTTTCGAGCTGGACTCAGTGGGCGTTGAGTTGGTCCCTTCGTTAG
ATCCTGGGTTCTGCCTACAGTCAGATTACACTCGGACCTCGTCGACATGCATTTCTCTTCTTGCAGGAAGAACTCAGCACTGCCCTCCTAAGTTTCGAGCTGGACTCAGTGGGCGTTGAGTTGGTCCCTTCGTTAG
ATCCTGGGTTCTGCCTACAGTCAGATTACACTCGGACCTCGTCGACATGCATTTCTGTTAGACGTTGATTAAGTTCTTGCAGTTTTTCCCCTCCTAAGTTTCGAGCTGGACTCAGTGGGCGTTGAGTTGGTCCCTTCGTTAG
ATCCTGGGTTCTGCCTACAGTCAGATTACACTCGGACCTCGTCGACATGCATTTTCCAAGAGCTCTTTTCTTAACTCCTCTTTCTCACCTCCTAAGTTTCGAGCTGGACTCAGTGGGCGTTGAGTTGGTCCCTTCGTTAG
ATCCTGGGTTCTGCCTACAGTCAGATTACACTCGGACCTCGTCGACATGCATTGTCAAGCCTCTCACCTTCTCTTCTGTCCACCTCCTAAGTTTCGAGCTGGACTCAGTGGGCGTTGAGTTGGTCCCTTCGTTAG
ATCCTGGGTTCTGCCTACAGTCAGATTACACTCGGACCTCGTCGACATGCATTGTTTTTCCTTCACAAACCAATGTTCCCTTAAGTCCCTCCTAAGTTTCGAGCTGGACTCAGTGGGCGTTGAGTTGGTCCCTTCGTTAG
ATCCTGGGTTCTGCCTACAGTCAGATTACACTCGGACCTCGTCGACATGCATTATTAAAGACTTGAGCTTCTCGGCCTTGTCCACCTCCTAAGTTTCGAGCTGGACTCAGTGGGCGTTGAGTTGGTCCCTTCGTTAG
ATCCTGGGTTCTGCCTACAGTCAGATTACACTCGGACCTCGTCGACATGCATTAGCCTTAAAGCTGGCCAGAGCCGCCTCCTCCTAAGTTTCGAGCTGGACTCAGTGGGCGTTGAGTTGGTCCCTTCGTTAG
ATCCTGGGTTCTGCCTACAGTCAGATTACACTCGGACCTCGTCGACATGCATTTGCTCCTGGTGAAACTGCTCTAGTTCCTGCCTCCTAAGTTTCGAGCTGGACTCAGTGGGCGTTGAGTTGGTCCCTTCGTTAG
ATCCTGGGTTCTGCCTACAGTCAGATTACACTCGGACCTCGTCGACATGCATTTCCGTAACGCTTCCCACCTCTAACATATTTGCCTCCTAAGTTTCGAGCTGGACTCAGTGGGCGTTGAGTTGGTCCCTTCGTTAG
ATCCTGGGTTCTGCCTACAGTCAGATTACACTCGGACCTCGTCGACATGCATTGTGCACGATATTCTTCCAGCTCAGACTGGACCTCCTAAGTTTCGAGCTGGACTCAGTGGGCGTTGAGTTGGTCCCTTCGTTAG
ATCCTGGGTTCTGCCTACAGTCAGATTACACTCGGACCTCGTCGACATGCATTAGCTGGCGTTGAAGCTGCAGAAACTGCTCCTCCTAAGTTTCGAGCTGGACTCAGTGGGCGTTGAGTTGGTCCCTTCGTTAG
ATCCTGGGTTCTGCCTACAGTCAGATTACACTCGGACCTCGTCGACATGCATTAGGGCCAGTGTTAATTCTGTCAAATTGATGTTTCCCTCCTAAGTTTCGAGCTGGACTCAGTGGGCGTTGAGTTGGTCCCTTCGTTAG
ATCCTGGGTTCTGCCTACAGTCAGATTACACTCGGACCTCGTCGACATGCATTGAGTGTGTTTCTCCACCAGGCTTGTCAGCCTCCTAAGTTTCGAGCTGGACTCAGTGGGCGTTGAGTTGGTCCCTTCGTTAG
ATCCTGGGTTCTGCCTACAGTCAGATTACACTCGGACCTCGTCGACATGCATTTCCTGGCTCTTTTTCCTTCTGGCATAGCATCCTCCTAAGTTTCGAGCTGGACTCAGTGGGCGTTGAGTTGGTCCCTTCGTTAG
ATCCTGGGTTCTGCCTACAGTCAGATTACACTCGGACCTCGTCGACATGCATTTTAGCCTTGCCTTGACTGGAAATACCTTTAAGACCTCCTAAGTTTCGAGCTGGACTCAGTGGGCGTTGAGTTGGTCCCTTCGTTAG
ATCCTGGGTTCTGCCTACAGTCAGATTACACTCGGACCTCGTCGACATGCATTATTGCTGATGAGGTTGTGGTACGTCGTCCCCTCCTAAGTTTCGAGCTGGACTCAGTGGGCGTTGAGTTGGTCCCTTCGTTAG
ATCCTGGGTTCTGCCTACAGTCAGATTACACTCGGACCTCGTCGACATGCATTAAAGACATCACTTCTTGTTTTAAAGCCTCCTTTTCCCCTCCTAAGTTTCGAGCTGGACTCAGTGGGCGTTGAGTTGGTCCCTTCGTTAG
ATCCTGGGTTCTGCCTACAGTCAGATTACACTCGGACCTCGTCGACATGCATTCTTCTAAATTCTGAACCTTTTCCTGAGAAGCCACCCTCCTAAGTTTCGAGCTGGACTCAGTGGGCGTTGAGTTGGTCCCTTCGTTAG
ATCCTGGGTTCTGCCTACAGTCAGATTACACTCGGACCTCGTCGACATGCATTCTCAGCTTCTCCTGCAGGTGAGTCTTTTCCCTCCTAAGTTTCGAGCTGGACTCAGTGGGCGTTGAGTTGGTCCCTTCGTTAG
ATCCTGGGTTCTGCCTACAGTCAGATTACACTCGGACCTCGTCGACATGCATTTCCACTCTGTGAAGCATTCAAGTCATCTGGCCTCCTAAGTTTCGAGCTGGACTCAGTGGGCGTTGAGTTGGTCCCTTCGTTAG
ATCCTGGGTTCTGCCTACAGTCAGATTACACTCGGACCTCGTCGACATGCATTTCATCTGTTCAATGACCAGCTCTGCCTCAACCTCCTAAGTTTCGAGCTGGACTCAGTGGGCGTTGAGTTGGTCCCTTCGTTAG
ATCCTGGGTTCTGCCTACAGTCAGATTACACTCGGACCTCGTCGACATGCATTGCTCATATTCAATGGATTGCATTCTTCAGAACCGCCTCCTAAGTTTCGAGCTGGACTCAGTGGGCGTTGAGTTGGTCCCTTCGTTAG
ATCCTGGGTTCTGCCTACAGTCAGATTACACTCGGACCTCGTCGACATGCATTAGAAGCCTTTTCACTAGTTTCAGTTTCTGTTCAACCCTCCTAAGTTTCGAGCTGGACTCAGTGGGCGTTGAGTTGGTCCCTTCGTTAG
ATCCTGGGTTCTGCCTACAGTCAGATTACACTCGGACCTCGTCGACATGCATTATCTTTGAGAAGGTGGTTTTCCTGTTGCACAATCCTCCTAAGTTTCGAGCTGGACTCAGTGGGCGTTGAGTTGGTCCCTTCGTTAG
ATCCTGGGTTCTGCCTACAGTCAGATTACACTCGGACCTCGTCGACATGCATTTGGGTCTGTATTTTAACTTCAGAGAGCTCCGCCTCCTAAGTTTCGAGCTGGACTCAGTGGGCGTTGAGTTGGTCCCTTCGTTAG
ATCCTGGGTTCTGCCTACAGTCAGATTACACTCGGACCTCGTCGACATGCATTCTCTTGTTCCCGTTCCTCCAATGCACTGTCCTCCTAAGTTTCGAGCTGGACTCAGTGGGCGTTGAGTTGGTCCCTTCGTTAG
ATCCTGGGTTCTGCCTACAGTCAGATTACACTCGGACCTCGTCGACATGCATTCTGTTAGCCTCAACTTCTTCTGACGGTGAGTCCTCCTAAGTTTCGAGCTGGACTCAGTGGGCGTTGAGTTGGTCCCTTCGTTAG
ATCCTGGGTTCTGCCTACAGTCAGATTACACTCGGACCTCGTCGACATGCATTCTTCAACGGAAGCCTGAGCACTCTGCCTCCTCCTAAGTTTCGAGCTGGACTCAGTGGGCGTTGAGTTGGTCCCTTCGTTAG
ATCCTGGGTTCTGCCTACAGTCAGATTACACTCGGACCTCGTCGACATGCATTTTCATCTACTTGGTCTTGTAGTACCCTGCACCCTCCTAAGTTTCGAGCTGGACTCAGTGGGCGTTGAGTTGGTCCCTTCGTTAG
ATCCTGGGTTCTGCCTACAGTCAGATTACACTCGGACCTCGTCGACATGCATTGCTGACAATAGCAACAATTTCTCCTTTCCTTCACCTCCTAAGTTTCGAGCTGGACTCAGTGGGCGTTGAGTTGGTCCCTTCGTTAG
ATCCTGGGTTCTGCCTACAGTCAGATTACACTCGGACCTCGTCGACATGCATTCTTCTACTCTCCCTTCACAAATCTAAGCTTAAGTCCTCCTAAGTTTCGAGCTGGACTCAGTGGGCGTTGAGTTGGTCCCTTCGTTAG
ATCCTGGGTTCTGCCTACAGTCAGATTACACTCGGACCTCGTCGACATGCATTGACACAGCTATTTAATTCCTCACTCAGAGCTTCCCTCCTAAGTTTCGAGCTGGACTCAGTGGGCGTTGAGTTGGTCCCTTCGTTAG
ATCCTGGGTTCTGCCTACAGTCAGATTACACTCGGACCTCGTCGACATGCATTTAGAGATTTTTTGCTGGAAGTCAGAGAGATCGGCCTCCTAAGTTTCGAGCTGGACTCAGTGGGCGTTGAGTTGGTCCCTTCGTTAG
ATCCTGGGTTCTGCCTACAGTCAGATTACACTCGGACCTCGTCGACATGCATTTCCTGAGATCCATTTAATGTCCCTAATTTCAGGCCTCCTAAGTTTCGAGCTGGACTCAGTGGGCGTTGAGTTGGTCCCTTCGTTAG
ATCCTGGGTTCTGCCTACAGTCAGATTACACTCGGACCTCGTCGACATGCATTCCAGGATTTCTGTTCCTGCTTCATCGTCGCCCTCCTAAGTTTCGAGCTGGACTCAGTGGGCGTTGAGTTGGTCCCTTCGTTAG
ATCCTGGGTTCTGCCTACAGTCAGATTACACTCGGACCTCGTCGACATGCATTACATCTGTGCAGCTGTTTCATTTTCTCCAGTTCCTCCTAAGTTTCGAGCTGGACTCAGTGGGCGTTGAGTTGGTCCCTTCGTTAG
ATCCTGGGTTCTGCCTACAGTCAGATTACACTCGGACCTCGTCGACATGCATTCTGACCACGGTTTCGTCCAGCTGCTTTTCCCTCCTAAGTTTCGAGCTGGACTCAGTGGGCGTTGAGTTGGTCCCTTCGTTAG
ATCCTGGGTTCTGCCTACAGTCAGATTACACTCGGACCTCGTCGACATGCATTTAATGGCGCACTTTATCTTCGAGCTCCAGTCCCTCCTAAGTTTCGAGCTGGACTCAGTGGGCGTTGAGTTGGTCCCTTCGTTAG
ATCCTGGGTTCTGCCTACAGTCAGATTACACTCGGACCTCGTCGACATGCATTGAGGCAACATATGTCCCTGTGATGTTGTTCTCCTCCTAAGTTTCGAGCTGGACTCAGTGGGCGTTGAGTTGGTCCCTTCGTTAG
ATCCTGGGTTCTGCCTACAGTCAGATTACACTCGGACCTCGTCGACATGCATTCCTCGAGCATCTCTCCATCCACATTCTGTCCTCCTAAGTTTCGAGCTGGACTCAGTGGGCGTTGAGTTGGTCCCTTCGTTAG
ATCCTGGGTTCTGCCTACAGTCAGATTACACTCGGACCTCGTCGACATGCATTTCTTCCGGTTCAGGTGACCATCACGGGCCTCCTAAGTTTCGAGCTGGACTCAGTGGGCGTTGAGTTGGTCCCTTCGTTAG
ATCCTGGGTTCTGCCTACAGTCAGATTACACTCGGACCTCGTCGACATGCATTTCTTCTATCCAGTCTTGGGGAGGGGAAGCCTCCTAAGTTTCGAGCTGGACTCAGTGGGCGTTGAGTTGGTCCCTTCGTTAG
ATCCTGGGTTCTGCCTACAGTCAGATTACACTCGGACCTCGTCGACATGCATTTTCATCAGATGATCCAACTTGTCCCAGGACCCTCCTAAGTTTCGAGCTGGACTCAGTGGGCGTTGAGTTGGTCCCTTCGTTAG
ATCCTGGGTTCTGCCTACAGTCAGATTACACTCGGACCTCGTCGACATGCATTTTCTCTTGCCAAAGCAGCTGCTGTTCCTCCCTCCTAAGTTTCGAGCTGGACTCAGTGGGCGTTGAGTTGGTCCCTTCGTTAG
ATCCTGGGTTCTGCCTACAGTCAGATTACACTCGGACCTCGTCGACATGCATTGCCTTTTCTGCTGGTTATGGAGCCCTGACCTCCTAAGTTTCGAGCTGGACTCAGTGGGCGTTGAGTTGGTCCCTTCGTTAG
ATCCTGGGTTCTGCCTACAGTCAGATTACACTCGGACCTCGTCGACATGCATTCCAAGCTCAACATGTGGCAAAGGTCGGTGCCTCCTAAGTTTCGAGCTGGACTCAGTGGGCGTTGAGTTGGTCCCTTCGTTAG
ATCCTGGGTTCTGCCTACAGTCAGATTACACTCGGACCTCGTCGACATGCATTTCCTCAAGCTTCACTTGCAGTTGTTTTTTCTCCCCTCCTAAGTTTCGAGCTGGACTCAGTGGGCGTTGAGTTGGTCCCTTCGTTAG
ATCCTGGGTTCTGCCTACAGTCAGATTACACTCGGACCTCGTCGACATGCATTTTTTTTCTAAGGTGTGCGTTTCGTTCTCATGCCCCTCCTAAGTTTCGAGCTGGACTCAGTGGGCGTTGAGTTGGTCCCTTCGTTAG
ATCCTGGGTTCTGCCTACAGTCAGATTACACTCGGACCTCGTCGACATGCATTTTGCTTCATGTTCTCCTGTGCCTTCTTGCACCTCCTAAGTTTCGAGCTGGACTCAGTGGGCGTTGAGTTGGTCCCTTCGTTAG
ATCCTGGGTTCTGCCTACAGTCAGATTACACTCGGACCTCGTCGACATGCATTGTGAATGCTGTTCTTGGTAACCAAAAGTTCATTTTCCCTCCTAAGTTTCGAGCTGGACTCAGTGGGCGTTGAGTTGGTCCCTTCGTTAG
ATCCTGGGTTCTGCCTACAGTCAGATTACACTCGGACCTCGTCGACATGCATTCTCCTGGCTGTTCTCAATGCCCTCTTCCCCTCCTAAGTTTCGAGCTGGACTCAGTGGGCGTTGAGTTGGTCCCTTCGTTAG
ATCCTGGGTTCTGCCTACAGTCAGATTACACTCGGACCTCGTCGACATGCATTGCCAGGTGTCCAGTATTCTCTCCACAGACCTCCTAAGTTTCGAGCTGGACTCAGTGGGCGTTGAGTTGGTCCCTTCGTTAG
ATCCTGGGTTCTGCCTACAGTCAGATTACACTCGGACCTCGTCGACATGCATTTATTTTTGTCCTTAGCCTAGGCCAATCCACAGCCTCCTAAGTTTCGAGCTGGACTCAGTGGGCGTTGAGTTGGTCCCTTCGTTAG
ATCCTGGGTTCTGCCTACAGTCAGATTACACTCGGACCTCGTCGACATGCATTTTTCTCTTGTGTTCCTGCCCATAAAGCAGTGCCTCCTAAGTTTCGAGCTGGACTCAGTGGGCGTTGAGTTGGTCCCTTCGTTAG
ATCCTGGGTTCTGCCTACAGTCAGATTACACTCGGACCTCGTCGACATGCATTCTCCTTGACCTCAGGTGGCATGCCAATGCCTCCTAAGTTTCGAGCTGGACTCAGTGGGCGTTGAGTTGGTCCCTTCGTTAG
ATCCTGGGTTCTGCCTACAGTCAGATTACACTCGGACCTCGTCGACATGCATTGTCTGCTTATCCCTCTTTTCTAAGAGTAGCTATTCCTCCTAAGTTTCGAGCTGGACTCAGTGGGCGTTGAGTTGGTCCCTTCGTTAG
ATCCTGGGTTCTGCCTACAGTCAGATTACACTCGGACCTCGTCGACATGCATTACTCCAAAGTGGTTTGTTCAGTGCTTTCCCTCCTCCTAAGTTTCGAGCTGGACTCAGTGGGCGTTGAGTTGGTCCCTTCGTTAG
ATCCTGGGTTCTGCCTACAGTCAGATTACACTCGGACCTCGTCGACATGCATTGCAAGGCGGTCCCGGATATAGTTCTCTTCCCTCCTAAGTTTCGAGCTGGACTCAGTGGGCGTTGAGTTGGTCCCTTCGTTAG
ATCCTGGGTTCTGCCTACAGTCAGATTACACTCGGACCTCGTCGACATGCATTGGATGAGTAGCTATCTCTGGGGCCCAGCCTCCTAAGTTTCGAGCTGGACTCAGTGGGCGTTGAGTTGGTCCCTTCGTTAG
ATCCTGGGTTCTGCCTACAGTCAGATTACACTCGGACCTCGTCGACATGCATTGCTGAGTCACTCGTAGGTCAGATTTCATCCCCTCCTAAGTTTCGAGCTGGACTCAGTGGGCGTTGAGTTGGTCCCTTCGTTAG
ATCCTGGGTTCTGCCTACAGTCAGATTACACTCGGACCTCGTCGACATGCATTGGACATTTGCAGGTTTACATTCTGCACGGTGCCTCCTAAGTTTCGAGCTGGACTCAGTGGGCGTTGAGTTGGTCCCTTCGTTAG
ATCCTGGGTTCTGCCTACAGTCAGATTACACTCGGACCTCGTCGACATGCATTAGCTGTGACTTTAAGCTCGCACTCTGATGTTCCTCCTAAGTTTCGAGCTGGACTCAGTGGGCGTTGAGTTGGTCCCTTCGTTAG
ATCCTGGGTTCTGCCTACAGTCAGATTACACTCGGACCTCGTCGACATGCATTGCAAGATCTCTTCCTATGCCTTGGAAGCCCCTCCTAAGTTTCGAGCTGGACTCAGTGGGCGTTGAGTTGGTCCCTTCGTTAG
ATCCTGGGTTCTGCCTACAGTCAGATTACACTCGGACCTCGTCGACATGCATTTGAGAGAGGTGGGTTTTACTATCCACATTTTCCCTCCTAAGTTTCGAGCTGGACTCAGTGGGCGTTGAGTTGGTCCCTTCGTTAG
ATCCTGGGTTCTGCCTACAGTCAGATTACACTCGGACCTCGTCGACATGCATTTTCCCTTTTCTTGAAACGAAGATCCCATCTTCTCCTCCTAAGTTTCGAGCTGGACTCAGTGGGCGTTGAGTTGGTCCCTTCGTTAG
ATCCTGGGTTCTGCCTACAGTCAGATTACACTCGGACCTCGTCGACATGCATTGCTCAATTTTTCTTGCTCTGTGGGATTCATGGCCTCCTAAGTTTCGAGCTGGACTCAGTGGGCGTTGAGTTGGTCCCTTCGTTAG
ATCCTGGGTTCTGCCTACAGTCAGATTACACTCGGACCTCGTCGACATGCATTGCATGGCCCATCCCATCATCCAGGCCCTCCTAAGTTTCGAGCTGGACTCAGTGGGCGTTGAGTTGGTCCCTTCGTTAG
ATCCTGGGTTCTGCCTACAGTCAGATTACACTCGGACCTCGTCGACATGCATTTCAGGCAGAGAACTGCACCCTTCTCTGCCTCCTAAGTTTCGAGCTGGACTCAGTGGGCGTTGAGTTGGTCCCTTCGTTAG
ATCCTGGGTTCTGCCTACAGTCAGATTACACTCGGACCTCGTCGACATGCATTATTCATCTGACTGACCTTCCTGTTAGCAGGACCTCCTAAGTTTCGAGCTGGACTCAGTGGGCGTTGAGTTGGTCCCTTCGTTAG
ATCCTGGGTTCTGCCTACAGTCAGATTACACTCGGACCTCGTCGACATGCATTTGTTCTGTACCATGGTCTGGAGCCTCTCCCTCCTAAGTTTCGAGCTGGACTCAGTGGGCGTTGAGTTGGTCCCTTCGTTAG
ATCCTGGGTTCTGCCTACAGTCAGATTACACTCGGACCTCGTCGACATGCATTGTTTCTGACATGATGACTTCTTGGGCAGTTTGCCTCCTAAGTTTCGAGCTGGACTCAGTGGGCGTTGAGTTGGTCCCTTCGTTAG
ATCCTGGGTTCTGCCTACAGTCAGATTACACTCGGACCTCGTCGACATGCATTGGAACTCAGGGCTTTCTGACAAGAGCTGTCCTCCTAAGTTTCGAGCTGGACTCAGTGGGCGTTGAGTTGGTCCCTTCGTTAG
ATCCTGGGTTCTGCCTACAGTCAGATTACACTCGGACCTCGTCGACATGCATTTGTGATGAACTTGGAAAGTGCCCTGCACCCCTCCTAAGTTTCGAGCTGGACTCAGTGGGCGTTGAGTTGGTCCCTTCGTTAG
ATCCTGGGTTCTGCCTACAGTCAGATTACACTCGGACCTCGTCGACATGCATTAGTAAGTCTTCCACCAACTGCTGGCGGACCTCCTAAGTTTCGAGCTGGACTCAGTGGGCGTTGAGTTGGTCCCTTCGTTAG
ATCCTGGGTTCTGCCTACAGTCAGATTACACTCGGACCTCGTCGACATGCATTCCTCATGCCTGCAAGTGGCCTCATGATGCCTCCTAAGTTTCGAGCTGGACTCAGTGGGCGTTGAGTTGGTCCCTTCGTTAG
ATCCTGGGTTCTGCCTACAGTCAGATTACACTCGGACCTCGTCGACATGCATTGTCAGATTCCTAACTATATTACTGAGGCTGGCCCTCCTAAGTTTCGAGCTGGACTCAGTGGGCGTTGAGTTGGTCCCTTCGTTAG
ATCCTGGGTTCTGCCTACAGTCAGATTACACTCGGACCTCGTCGACATGCATTTACTTCCAACCACTGAGTTCTCCATGAGCTCCTCCTAAGTTTCGAGCTGGACTCAGTGGGCGTTGAGTTGGTCCCTTCGTTAG
ATCCTGGGTTCTGCCTACAGTCAGATTACACTCGGACCTCGTCGACATGCATTCTCATCCAGTTTTACTTGTTTCTTTTTCAGGCCTTCCTCCTAAGTTTCGAGCTGGACTCAGTGGGCGTTGAGTTGGTCCCTTCGTTAG
ATCCTGGGTTCTGCCTACAGTCAGATTACACTCGGACCTCGTCGACATGCATTTTCGGAGGTCCTGTTTTCAAGTTCCTCCTGCCTCCTAAGTTTCGAGCTGGACTCAGTGGGCGTTGAGTTGGTCCCTTCGTTAG
ATCCTGGGTTCTGCCTACAGTCAGATTACACTCGGACCTCGTCGACATGCATTACCTGATGGGTTTAGATGTTTTTGGTGCTTGGCCTCCTAAGTTTCGAGCTGGACTCAGTGGGCGTTGAGTTGGTCCCTTCGTTAG
ATCCTGGGTTCTGCCTACAGTCAGATTACACTCGGACCTCGTCGACATGCATTTATGAGTGTACCCTTTGGTTTGGCTATGACCCCTCCTAAGTTTCGAGCTGGACTCAGTGGGCGTTGAGTTGGTCCCTTCGTTAG
ATCCTGGGTTCTGCCTACAGTCAGATTACACTCGGACCTCGTCGACATGCATTCTGCTTGCCTGCCTGCTGCAGGATCTCCTCCTAAGTTTCGAGCTGGACTCAGTGGGCGTTGAGTTGGTCCCTTCGTTAG
ATCCTGGGTTCTGCCTACAGTCAGATTACACTCGGACCTCGTCGACATGCATTTCTTGTAGTGAGGCCTCCTGGGTTGACCCTCCTAAGTTTCGAGCTGGACTCAGTGGGCGTTGAGTTGGTCCCTTCGTTAG
ATCCTGGGTTCTGCCTACAGTCAGATTACACTCGGACCTCGTCGACATGCATTTCTTGAGCATGAGTGGAAAGTCCTATTTGGCCCTCCTAAGTTTCGAGCTGGACTCAGTGGGCGTTGAGTTGGTCCCTTCGTTAG
ATCCTGGGTTCTGCCTACAGTCAGATTACACTCGGACCTCGTCGACATGCATTCGCATGGTGATCATCCACCTCCGAGGCCCTCCTAAGTTTCGAGCTGGACTCAGTGGGCGTTGAGTTGGTCCCTTCGTTAG
ATCCTGGGTTCTGCCTACAGTCAGATTACACTCGGACCTCGTCGACATGCATTATCTGACCGTAGCTTCTCTTTTTCTCTGACCACCTCCTAAGTTTCGAGCTGGACTCAGTGGGCGTTGAGTTGGTCCCTTCGTTAG
ATCCTGGGTTCTGCCTACAGTCAGATTACACTCGGACCTCGTCGACATGCATTCTCTCTCTCTTTTCGGAGTTCATTTTTTAAGGCTGCCTCCTAAGTTTCGAGCTGGACTCAGTGGGCGTTGAGTTGGTCCCTTCGTTAG
ATCCTGGGTTCTGCCTACAGTCAGATTACACTCGGACCTCGTCGACATGCATTTGTTCCTGGTTTCCCTGTGGTGTGTTGGCCTCCTAAGTTTCGAGCTGGACTCAGTGGGCGTTGAGTTGGTCCCTTCGTTAG
ATCCTGGGTTCTGCCTACAGTCAGATTACACTCGGACCTCGTCGACATGCATTGAGGTTGTACTCATTCCGCCGCTCTATGGCCTCCTAAGTTTCGAGCTGGACTCAGTGGGCGTTGAGTTGGTCCCTTCGTTAG
ATCCTGGGTTCTGCCTACAGTCAGATTACACTCGGACCTCGTCGACATGCATTCTTCACCATTGCGTCTGCCTTAGTGTTCTTGCCTCCTAAGTTTCGAGCTGGACTCAGTGGGCGTTGAGTTGGTCCCTTCGTTAG
ATCCTGGGTTCTGCCTACAGTCAGATTACACTCGGACCTCGTCGACATGCATTAGGTGCTGGTTTATCCTTTCTGCCTGCAGCCTCCTAAGTTTCGAGCTGGACTCAGTGGGCGTTGAGTTGGTCCCTTCGTTAG

ATCCTGGGTTCTGCCTACAGTCAGATTACACTCGGACCTCGTCGACATGCATTTCTATCATTCGTTCCTCCATGACAGTTACCAGCCTCCTAAGTTTCGAGCTGGACTCAGTGGGCGTTGAGTTGGTCCCTTCGTTAG
ATCCTGGGTTCTGCCTACAGTCAGATTACACTCGGACCTCGTCGACATGCATTCTGTTTTCTGTCGCTGTTCAGTCACTTCCAGCCTCCTAAGTTTCGAGCTGGACTCAGTGGGCGTTGAGTTGGTCCCTTCGTTAG
ATCCTGGGTTCTGCCTACAGTCAGATTACACTCGGACCTCGTCGACATGCATTAGAGTTGTTCTTTGAGCTGATTCACTTTCTCTTGCCTCCTAAGTTTCGAGCTGGACTCAGTGGGCGTTGAGTTGGTCCCTTCGTTAG

**NEK9**
GTCCCTAATTGCCTCTCGGATTCCT TTACACTCGGACCTCGTCGACATGGTTACTGCAGGATTCAGCCGAGGTGACGACCTCCTAAGTTTCGAGCTGGACTCAGTGGATCTGGCCTGTGAGTGTTGGTGTG
GTCCCTAATTGCCTCTCGGATTCCT TTACACTCGGACCTCGTCGACATGAGACCCTGATCACTTGCTTCCCACCAAGGCCTCCTAAGTTTCGAGCTGGACTCAGTGGATCTGGCCTGTGAGTGTTGGTGTG
GTCCCTAATTGCCTCTCGGATTCCT TTACACTCGGACCTCGTCGACATGTTGGCCAAGGTAAAGGACGTTGTGTAGGGACCTCCTAAGTTTCGAGCTGGACTCAGTGGATCTGGCCTGTGAGTGTTGGTGTG
GTCCCTAATTGCCTCTCGGATTCCT TTACACTCGGACCTCGTCGACATGATTCATTGAGTCCACAGGCCAGCACTTTGCCCTCCTAAGTTTCGAGCTGGACTCAGTGGATCTGGCCTGTGAGTGTTGGTGTG
GTCCCTAATTGCCTCTCGGATTCCT TTACACTCGGACCTCGTCGACATGTCCTCTTCTGAATCCAAACCCAGTCGTCCACCTCCTAAGTTTCGAGCTGGACTCAGTGGATCTGGCCTGTGAGTGTTGGTGTG
GTCCCTAATTGCCTCTCGGATTCCT TTACACTCGGACCTCGTCGACATGTCCACTGGATTGCTGAGGAAGAAGTTCAGCCCTCCTAAGTTTCGAGCTGGACTCAGTGGATCTGGCCTGTGAGTGTTGGTGTG
GTCCCTAATTGCCTCTCGGATTCCT TTACACTCGGACCTCGTCGACATGCATGGGTTCTAGCACTTCAGGGCCAGCAACCTCCTAAGTTTCGAGCTGGACTCAGTGGATCTGGCCTGTGAGTGTTGGTGTG
GTCCCTAATTGCCTCTCGGATTCCT TTACACTCGGACCTCGTCGACATGCTACAGCAATGGGTGCTTCAGTCACAGTGCCCTCCTAAGTTTCGAGCTGGACTCAGTGGATCTGGCCTGTGAGTGTTGGTGTG
GTCCCTAATTGCCTCTCGGATTCCT TTACACTCGGACCTCGTCGACATGAGAGTACTGGCTAGAGTCAACTTCCATGGCCCTCCTAAGTTTCGAGCTGGACTCAGTGGATCTGGCCTGTGAGTGTTGGTGTG
GTCCCTAATTGCCTCTCGGATTCCT TTACACTCGGACCTCGTCGACATGAGCTCTGGAGACATGTAATATGGGGTTCCCCCTCCTAAGTTTCGAGCTGGACTCAGTGGATCTGGCCTGTGAGTGTTGGTGTG
GTCCCTAATTGCCTCTCGGATTCCT TTACACTCGGACCTCGTCGACATGTCAAGGCATCACGACGTTCCTTCTCAGACACCTCCTAAGTTTCGAGCTGGACTCAGTGGATCTGGCCTGTGAGTGTTGGTGTG
GTCCCTAATTGCCTCTCGGATTCCT TTACACTCGGACCTCGTCGACATGTCGACTTCCTTCCACACAACCAGTGAGTCACCTCCTAAGTTTCGAGCTGGACTCAGTGGATCTGGCCTGTGAGTGTTGGTGTG
GTCCCTAATTGCCTCTCGGATTCCT TTACACTCGGACCTCGTCGACATGCTAGAGGCTGGGTCTACAGGAGTCTGTTCCCTCCTAAGTTTCGAGCTGGACTCAGTGGATCTGGCCTGTGAGTGTTGGTGTG
GTCCCTAATTGCCTCTCGGATTCCT TTACACTCGGACCTCGTCGACATGTGCTGTCTGAGTTCCTTTGGAATGCATCCCCCTCCTAAGTTTCGAGCTGGACTCAGTGGATCTGGCCTGTGAGTGTTGGTGTG
GTCCCTAATTGCCTCTCGGATTCCT TTACACTCGGACCTCGTCGACATGTGCTGTAGCTTCTGTTGTTCAGCCAGACACCCTCCTAAGTTTCGAGCTGGACTCAGTGGATCTGGCCTGTGAGTGTTGGTGTG
GTCCCTAATTGCCTCTCGGATTCCT TTACACTCGGACCTCGTCGACATGGGGGTTTGTGTTCCAAAGGAGCTTCAGAGGCCTCCTAAGTTTCGAGCTGGACTCAGTGGATCTGGCCTGTGAGTGTTGGTGTG
GTCCCTAATTGCCTCTCGGATTCCT TTACACTCGGACCTCGTCGACATGACTTTGAGTCCTTGCAGCTCTTCATAGGGCCCTCCTAAGTTTCGAGCTGGACTCAGTGGATCTGGCCTGTGAGTGTTGGTGTG
GTCCCTAATTGCCTCTCGGATTCCT TTACACTCGGACCTCGTCGACATGCTGCATTTTCCAGCTCCTTTCGAAGCCAGCCCTCCTAAGTTTCGAGCTGGACTCAGTGGATCTGGCCTGTGAGTGTTGGTGTG
GTCCCTAATTGCCTCTCGGATTCCT TTACACTCGGACCTCGTCGACATGTGGCCTCTGTGGGACTGATTAAACCTTCCACCTCCTAAGTTTCGAGCTGGACTCAGTGGATCTGGCCTGTGAGTGTTGGTGTG
GTCCCTAATTGCCTCTCGGATTCCT TTACACTCGGACCTCGTCGACATGCCTCGGTCTGCTTCCATTGTTCCTCGGAAGCCTCCTAAGTTTCGAGCTGGACTCAGTGGATCTGGCCTGTGAGTGTTGGTGTG
GTCCCTAATTGCCTCTCGGATTCCT TTACACTCGGACCTCGTCGACATGCCACTTGGGTCAGGAGTTTCAGATTCCTGCCCTCCTAAGTTTCGAGCTGGACTCAGTGGATCTGGCCTGTGAGTGTTGGTGTG
GTCCCTAATTGCCTCTCGGATTCCT TTACACTCGGACCTCGTCGACATGGCCATGTGGTCTCTCTGTGGGGGTCATTCCTCCTAAGTTTCGAGCTGGACTCAGTGGATCTGGCCTGTGAGTGTTGGTGTG
GTCCCTAATTGCCTCTCGGATTCCT TTACACTCGGACCTCGTCGACATGTCAATAGCAGCTGTGTGAGTCTTGCCTGGGCCTCCTAAGTTTCGAGCTGGACTCAGTGGATCTGGCCTGTGAGTGTTGGTGTG
GTCCCTAATTGCCTCTCGGATTCCT TTACACTCGGACCTCGTCGACATGATTCCCGACATGCACTGATTCAGACCCAGCCCTCCTAAGTTTCGAGCTGGACTCAGTGGATCTGGCCTGTGAGTGTTGGTGTG
GTCCCTAATTGCCTCTCGGATTCCT TTACACTCGGACCTCGTCGACATGAAGGCCTTGGGAACATCCACCTTTTGTGGTCCTCCTAAGTTTCGAGCTGGACTCAGTGGATCTGGCCTGTGAGTGTTGGTGTG
GTCCCTAATTGCCTCTCGGATTCCT TTACACTCGGACCTCGTCGACATGTTCCACATGCTTTGGCTGTCGATAGGAGGCCCTCCTAAGTTTCGAGCTGGACTCAGTGGATCTGGCCTGTGAGTGTTGGTGTG
GTCCCTAATTGCCTCTCGGATTCCT TTACACTCGGACCTCGTCGACATGTGTCTCCATGGCCCAGCTGACCATGGACCTCCTAAGTTTCGAGCTGGACTCAGTGGATCTGGCCTGTGAGTGTTGGTGTG
GTCCCTAATTGCCTCTCGGATTCCT TTACACTCGGACCTCGTCGACATGTCCTTCTCCACTGTGACCACAGCAAAGTGGCCTCCTAAGTTTCGAGCTGGACTCAGTGGATCTGGCCTGTGAGTGTTGGTGTG
GTCCCTAATTGCCTCTCGGATTCCT TTACACTCGGACCTCGTCGACATGACATCCAGTTTCTGGGGGGTGGATTTTCCACCTCCTAAGTTTCGAGCTGGACTCAGTGGATCTGGCCTGTGAGTGTTGGTGTG
GTCCCTAATTGCCTCTCGGATTCCT TTACACTCGGACCTCGTCGACATGAGTTCATCTGCAGTAGGTCTCTGCTCAGGACCTCCTAAGTTTCGAGCTGGACTCAGTGGATCTGGCCTGTGAGTGTTGGTGTG
GTCCCTAATTGCCTCTCGGATTCCT TTACACTCGGACCTCGTCGACATGTCCAGCTTTATGGATGCAGCTCACTGCTGACCTCCTAAGTTTCGAGCTGGACTCAGTGGATCTGGCCTGTGAGTGTTGGTGTG
GTCCCTAATTGCCTCTCGGATTCCT TTACACTCGGACCTCGTCGACATGTCCAGCTCAATCAGCAGCGTGGTATTGTCCCCTCCTAAGTTTCGAGCTGGACTCAGTGGATCTGGCCTGTGAGTGTTGGTGTG
GTCCCTAATTGCCTCTCGGATTCCT TTACACTCGGACCTCGTCGACATGGGCAATAATGTTGTCGTGCTGCAGCAGTGCCCTCCTAAGTTTCGAGCTGGACTCAGTGGATCTGGCCTGTGAGTGTTGGTGTG
GTCCCTAATTGCCTCTCGGATTCCT TTACACTCGGACCTCGTCGACATGCGAGTTGATGGAATCGCAGTGTCGCTCGTACCTCCTAAGTTTCGAGCTGGACTCAGTGGATCTGGCCTGTGAGTGTTGGTGTG
GTCCCTAATTGCCTCTCGGATTCCT TTACACTCGGACCTCGTCGACATGAGGCACCAGGAATCTGAATCTAAGTCAGGCCCTCCTAAGTTTCGAGCTGGACTCAGTGGATCTGGCCTGTGAGTGTTGGTGTG
GTCCCTAATTGCCTCTCGGATTCCT TTACACTCGGACCTCGTCGACATGTAACACCAGACCCTGCAATCTCTCAACCTCCCTCCTAAGTTTCGAGCTGGACTCAGTGGATCTGGCCTGTGAGTGTTGGTGTG
GTCCCTAATTGCCTCTCGGATTCCT TTACACTCGGACCTCGTCGACATGGACAGGAGCTGCTGGCCCCATTACTGTTCCTCCTAAGTTTCGAGCTGGACTCAGTGGATCTGGCCTGTGAGTGTTGGTGTG
GTCCCTAATTGCCTCTCGGATTCCT TTACACTCGGACCTCGTCGACATGCAACGATGAGAATGGTATGCCATCCACGGCCCTCCTAAGTTTCGAGCTGGACTCAGTGGATCTGGCCTGTGAGTGTTGGTGTG
GTCCCTAATTGCCTCTCGGATTCCT TTACACTCGGACCTCGTCGACATGACAGGTCCGGGACATGATGCAGAGATCCACCTCCTAAGTTTCGAGCTGGACTCAGTGGATCTGGCCTGTGAGTGTTGGTGTG
GTCCCTAATTGCCTCTCGGATTCCT TTACACTCGGACCTCGTCGACATGATAGGCCGAGGCCATGAGGTACAGATATCACCTCCTAAGTTTCGAGCTGGACTCAGTGGATCTGGCCTGTGAGTGTTGGTGTG
GTCCCTAATTGCCTCTCGGATTCCT TTACACTCGGACCTCGTCGACATGCAGTGGCAGCAATGGTAAACTCATCACCGCCCTCCTAAGTTTCGAGCTGGACTCAGTGGATCTGGCCTGTGAGTGTTGGTGTG
GTCCCTAATTGCCTCTCGGATTCCT TTACACTCGGACCTCGTCGACATGACTGGGTCAACAGAAATGTCCCATCACAGCCCTCCTAAGTTTCGAGCTGGACTCAGTGGATCTGGCCTGTGAGTGTTGGTGTG
GTCCCTAATTGCCTCTCGGATTCCT TTACACTCGGACCTCGTCGACATGTTCGCCACAGCCCCAAGAATAGACTTCCTTCCTCCTAAGTTTCGAGCTGGACTCAGTGGATCTGGCCTGTGAGTGTTGGTGTG
GTCCCTAATTGCCTCTCGGATTCCT TTACACTCGGACCTCGTCGACATGCGAAGGCATAGAGCTGACCCTCATCAGTCACCTCCTAAGTTTCGAGCTGGACTCAGTGGATCTGGCCTGTGAGTGTTGGTGTG
GTCCCTAATTGCCTCTCGGATTCCT TTACACTCGGACCTCGTCGACATGTAGTGCCTCCTTGCATGTTCACCCAAGTGTCCTCCTAAGTTTCGAGCTGGACTCAGTGGATCTGGCCTGTGAGTGTTGGTGTG
GTCCCTAATTGCCTCTCGGATTCCT TTACACTCGGACCTCGTCGACATGTTTCCTCCATCTCTCTCCTGCGTTTCCTGACCTCCTAAGTTTCGAGCTGGACTCAGTGGATCTGGCCTGTGAGTGTTGGTGTG
GTCCCTAATTGCCTCTCGGATTCCT TTACACTCGGACCTCGTCGACATGTCTGAAAACGCTGCACTGAGAGGAGATGGGCCTCCTAAGTTTCGAGCTGGACTCAGTGGATCTGGCCTGTGAGTGTTGGTGTG
GTCCCTAATTGCCTCTCGGATTCCT TTACACTCGGACCTCGTCGACATGACCACCACATGATTATCTCCACAGGAGACCCCTCCTAAGTTTCGAGCTGGACTCAGTGGATCTGGCCTGTGAGTGTTGGTGTG
GTCCCTAATTGCCTCTCGGATTCCT TTACACTCGGACCTCGTCGACATGTCATCACCACATGACACCTGACGGATAGCTCCTCCTAAGTTTCGAGCTGGACTCAGTGGATCTGGCCTGTGAGTGTTGGTGTG
GTCCCTAATTGCCTCTCGGATTCCT TTACACTCGGACCTCGTCGACATGGCAGTTCAAAAATGACGCAGCCAACTGCCCCCTCCTAAGTTTCGAGCTGGACTCAGTGGATCTGGCCTGTGAGTGTTGGTGTG
GTCCCTAATTGCCTCTCGGATTCCT TTACACTCGGACCTCGTCGACATGGGTACCACACCACCATCTCTTCCTCAAACACCTCCTAAGTTTCGAGCTGGACTCAGTGGATCTGGCCTGTGAGTGTTGGTGTG
GTCCCTAATTGCCTCTCGGATTCCT TTACACTCGGACCTCGTCGACATGGTCCACCCCCATGCAGCCATAATAATCTGACCTCCTAAGTTTCGAGCTGGACTCAGTGGATCTGGCCTGTGAGTGTTGGTGTG

**CHD3**
GTCTCCCGCCGCACAGTCTCTAATG TTACACTCGGACCTCGTCGACATGGTGAAGCTCTGTGAAGCCACCATCGGCGACCTCCTAAGTTTCGAGCTGGACTCAGTGCAGAGACTAGCCTCCAGTACCCACC
GTCTCCCGCCGCACAGTCTCTAATG TTACACTCGGACCTCGTCGACATGCAGAAGCCTCAGGAGTGGTGGGAGACGTTCCTCCTAAGTTTCGAGCTGGACTCAGTGCAGAGACTAGCCTCCAGTACCCACC
GTCTCCCGCCGCACAGTCTCTAATG TTACACTCGGACCTCGTCGACATGTTGACGAGAGACATGACTCCAATGCGGGTCCCTCCTAAGTTTCGAGCTGGACTCAGTGCAGAGACTAGCCTCCAGTACCCACC
GTCTCCCGCCGCACAGTCTCTAATG TTACACTCGGACCTCGTCGACATGACAGCATTGAGGAAAGCCTTCCGCTGACGGCCTCCTAAGTTTCGAGCTGGACTCAGTGCAGAGACTAGCCTCCAGTACCCACC
GTCTCCCGCCGCACAGTCTCTAATG TTACACTCGGACCTCGTCGACATGTTGGTCATCTGCGAGAAGATGAGCACTCGGCCTCCTAAGTTTCGAGCTGGACTCAGTGCAGAGACTAGCCTCCAGTACCCACC
GTCTCCCGCCGCACAGTCTCTAATG TTACACTCGGACCTCGTCGACATGTATTAAGCAGCGACACCTGGTTCCCACCACCCTCCTAAGTTTCGAGCTGGACTCAGTGCAGAGACTAGCCTCCAGTACCCACC
GTCTCCCGCCGCACAGTCTCTAATG TTACACTCGGACCTCGTCGACATGAAGACATCTGCCTTGAGTCTCCGCAGCATGCCTCCTAAGTTTCGAGCTGGACTCAGTGCAGAGACTAGCCTCCAGTACCCACC
GTCTCCCGCCGCACAGTCTCTAATG TTACACTCGGACCTCGTCGACATGAATCTCTCTGGGGTGAGGAAGTTCAGGAGACCTCCTAAGTTTCGAGCTGGACTCAGTGCAGAGACTAGCCTCCAGTACCCACC
GTCTCCCGCCGCACAGTCTCTAATG TTACACTCGGACCTCGTCGACATGCCAAGTGCTGCCTGATCAATGGTGATCAGCCCTCCTAAGTTTCGAGCTGGACTCAGTGCAGAGACTAGCCTCCAGTACCCACC
GTCTCCCGCCGCACAGTCTCTAATG TTACACTCGGACCTCGTCGACATGGTCAGGAGAACATGGAACTTCACCTGTGCCCCTCCTAAGTTTCGAGCTGGACTCAGTGCAGAGACTAGCCTCCAGTACCCACC
GTCTCCCGCCGCACAGTCTCTAATG TTACACTCGGACCTCGTCGACATGAGAATTTGGGTGCCCACATCTGGAACTCCCCCTCCTAAGTTTCGAGCTGGACTCAGTGCAGAGACTAGCCTCCAGTACCCACC
GTCTCCCGCCGCACAGTCTCTAATG TTACACTCGGACCTCGTCGACATGTCCCAGTTAATGATGGTAGAGAGTGGGGCACCTCCTAAGTTTCGAGCTGGACTCAGTGCAGAGACTAGCCTCCAGTACCCACC
GTCTCCCGCCGCACAGTCTCTAATG TTACACTCGGACCTCGTCGACATGCCAGGAGAAGCGTAGCCAGTTCAGCCCTTCCTCCTAAGTTTCGAGCTGGACTCAGTGCAGAGACTAGCCTCCAGTACCCACC
GTCTCCCGCCGCACAGTCTCTAATG TTACACTCGGACCTCGTCGACATGAACTGATACATGTGCAGGGTGCCTCCAGTGCCTCCTAAGTTTCGAGCTGGACTCAGTGCAGAGACTAGCCTCCAGTACCCACC
GTCTCCCGCCGCACAGTCTCTAATG TTACACTCGGACCTCGTCGACATGGTAGGATCATTAGTGGGAGAACTGGGAGGCCCTCCTAAGTTTCGAGCTGGACTCAGTGCAGAGACTAGCCTCCAGTACCCACC
GTCTCCCGCCGCACAGTCTCTAATG TTACACTCGGACCTCGTCGACATGGGAGCAGTGCCAGTAGGATAGTCCTACCCCCTCCTAAGTTTCGAGCTGGACTCAGTGCAGAGACTAGCCTCCAGTACCCACC
GTCTCCCGCCGCACAGTCTCTAATG TTACACTCGGACCTCGTCGACATGAGCCATTCACCATTGGGAATGTCAGGCAGGCCTCCTAAGTTTCGAGCTGGACTCAGTGCAGAGACTAGCCTCCAGTACCCACC
GTCTCCCGCCGCACAGTCTCTAATG TTACACTCGGACCTCGTCGACATGATGAATGTGGTAGGAGGAGATGCACGCGTCCCTCCTAAGTTTCGAGCTGGACTCAGTGCAGAGACTAGCCTCCAGTACCCACC
GTCTCCCGCCGCACAGTCTCTAATG TTACACTCGGACCTCGTCGACATGAGCAGCTGACACCTGCTCAGCTACAGCTCCTCCTAAGTTTCGAGCTGGACTCAGTGCAGAGACTAGCCTCCAGTACCCACC
GTCTCCCGCCGCACAGTCTCTAATG TTACACTCGGACCTCGTCGACATGTAGCAATTAGGGGCCTCATGAACTGGCTGACCTCCTAAGTTTCGAGCTGGACTCAGTGCAGAGACTAGCCTCCAGTACCCACC
GTCTCCCGCCGCACAGTCTCTAATG TTACACTCGGACCTCGTCGACATGCTGTGATGAAGGACCCTGCAGGCATCTGGCCTCCTAAGTTTCGAGCTGGACTCAGTGCAGAGACTAGCCTCCAGTACCCACC
GTCTCCCGCCGCACAGTCTCTAATG TTACACTCGGACCTCGTCGACATGAGCAACTCCTCCAGCTGGTTCAGAACCTTGCCTCCTAAGTTTCGAGCTGGACTCAGTGCAGAGACTAGCCTCCAGTACCCACC
GTCTCCCGCCGCACAGTCTCTAATG TTACACTCGGACCTCGTCGACATGTTGAGCATCATTCTGGATGTCCTGCCACCGCCTCCTAAGTTTCGAGCTGGACTCAGTGCAGAGACTAGCCTCCAGTACCCACC
GTCTCCCGCCGCACAGTCTCTAATG TTACACTCGGACCTCGTCGACATGTGAACCGGGGCTTCTCTGTCTTTTCCTCTCCCTCCTAAGTTTCGAGCTGGACTCAGTGCAGAGACTAGCCTCCAGTACCCACC
GTCTCCCGCCGCACAGTCTCTAATG TTACACTCGGACCTCGTCGACATGCCTGGTCGAAGCTCCCGGTCACCTTTTACACCTCCTAAGTTTCGAGCTGGACTCAGTGCAGAGACTAGCCTCCAGTACCCACC
GTCTCCCGCCGCACAGTCTCTAATG TTACACTCGGACCTCGTCGACATGTCGACTCTGTGGCTGACTTCTCTCTGTCCCCTCCTAAGTTTCGAGCTGGACTCAGTGCAGAGACTAGCCTCCAGTACCCACC
GTCTCCCGCCGCACAGTCTCTAATG TTACACTCGGACCTCGTCGACATGACGGTACCCAGGTTCAGGCTCCATCTCTCCCTCCTAAGTTTCGAGCTGGACTCAGTGCAGAGACTAGCCTCCAGTACCCACC
GTCTCCCGCCGCACAGTCTCTAATG TTACACTCGGACCTCGTCGACATGATGCCTTCTCCTTTCTCACTTGGAGCTGGACCTCCTAAGTTTCGAGCTGGACTCAGTGCAGAGACTAGCCTCCAGTACCCACC
GTCTCCCGCCGCACAGTCTCTAATG TTACACTCGGACCTCGTCGACATGAGCAGGTTTAGAGGTGCAGGGACTGTTGGTCCTCCTAAGTTTCGAGCTGGACTCAGTGCAGAGACTAGCCTCCAGTACCCACC
GTCTCCCGCCGCACAGTCTCTAATG TTACACTCGGACCTCGTCGACATGCGGCATTGACCAACGCCCATTGATGTGCTCCCTCCTAAGTTTCGAGCTGGACTCAGTGCAGAGACTAGCCTCCAGTACCCACC
GTCTCCCGCCGCACAGTCTCTAATG TTACACTCGGACCTCGTCGACATGTCTCATTCCGGAGCTGCCTCTTTGACTGTCCCTCCTAAGTTTCGAGCTGGACTCAGTGCAGAGACTAGCCTCCAGTACCCACC
GTCTCCCGCCGCACAGTCTCTAATG TTACACTCGGACCTCGTCGACATGTCGAAGTCTTCATCCTCCTCCTCTGAACCCCCTCCTAAGTTTCGAGCTGGACTCAGTGCAGAGACTAGCCTCCAGTACCCACC
GTCTCCCGCCGCACAGTCTCTAATG TTACACTCGGACCTCGTCGACATGCTCTGACTGGTTGTCTTGGTCTTCCTGAGCCCTCCTAAGTTTCGAGCTGGACTCAGTGCAGAGACTAGCCTCCAGTACCCACC
GTCTCCCGCCGCACAGTCTCTAATG TTACACTCGGACCTCGTCGACATGAGTTAACTTGCTTGCGAACCCGCTTGCCCTCCTCCTAAGTTTCGAGCTGGACTCAGTGCAGAGACTAGCCTCCAGTACCCACC
GTCTCCCGCCGCACAGTCTCTAATG TTACACTCGGACCTCGTCGACATGCCTAGATTCCGGGCTAGGTCTTCCTGCTGCCTCCTAAGTTTCGAGCTGGACTCAGTGCAGAGACTAGCCTCCAGTACCCACC
GTCTCCCGCCGCACAGTCTCTAATG TTACACTCGGACCTCGTCGACATGCTCATAGTGATGCCTCAGCAGCTTCTCCCACCTCCTAAGTTTCGAGCTGGACTCAGTGCAGAGACTAGCCTCCAGTACCCACC

GTCTCCCGCCGCACAGTCTCTAATG TTACACTCGGACCTCGTCGACATGGTCAGGGTCCACATTCTCCTCCTGCTTGATCCTCCTAAGTTTCGAGCTGGACTCAGTGCAGAGACTAGCCTCCAGTACCCACC

GTCTCCCGCCGCACAGTCTCTAATG TTACACTCGGACCTCGTCGACATGCTTGTCTTCTTCCCGCACGACGTACTGTGCCCTCCTAAGTTTCGAGCTGGACTCAGTGCAGAGACTAGCCTCCAGTACCCACC

GTCTCCCGCCGCACAGTCTCTAATG TTACACTCGGACCTCGTCGACATGGTTCTGCACGTCAGTGTCCTCAGTTGCATCCCTCCTAAGTTTCGAGCTGGACTCAGTGCAGAGACTAGCCTCCAGTACCCACC

GTCTCCCGCCGCACAGTCTCTAATG TTACACTCGGACCTCGTCGACATGATCACACTGCTGTCCTCCTCCTTGTTCTCCCCTCCTAAGTTTCGAGCTGGACTCAGTGCAGAGACTAGCCTCCAGTACCCACC

GTCTCCCGCCGCACAGTCTCTAATG TTACACTCGGACCTCGTCGACATGATGTCGTCAAGCTCCTGCTTGGACATGGAGCCTCCTAAGTTTCGAGCTGGACTCAGTGCAGAGACTAGCCTCCAGTACCCACC

GTCTCCCGCCGCACAGTCTCTAATG TTACACTCGGACCTCGTCGACATGTGGCCACTTGTGTGATTCGCTCTTCCACTGCCTCCTAAGTTTCGAGCTGGACTCAGTGCAGAGACTAGCCTCCAGTACCCACC

GTCTCCCGCCGCACAGTCTCTAATG TTACACTCGGACCTCGTCGACATGTCACTTTGTTGGCCTGGCCAATCCGATGAGCCTCCTAAGTTTCGAGCTGGACTCAGTGCAGAGACTAGCCTCCAGTACCCACC

GTCTCCCGCCGCACAGTCTCTAATG TTACACTCGGACCTCGTCGACATGCGGCTAAAGGCCTGGATGTCATTATGGGGGCCTCCTAAGTTTCGAGCTGGACTCAGTGCAGAGACTAGCCTCCAGTACCCACC

GTCTCCCGCCGCACAGTCTCTAATG TTACACTCGGACCTCGTCGACATGTTGTTGGGCCCCAGGAGCATTAAACCGATCCCTCCTAAGTTTCGAGCTGGACTCAGTGCAGAGACTAGCCTCCAGTACCCACC

GTCTCCCGCCGCACAGTCTCTAATG TTACACTCGGACCTCGTCGACATGTGCTCCTTCAGCTTTCGCAGCATCTTCTGGCCTCCTAAGTTTCGAGCTGGACTCAGTGCAGAGACTAGCCTCCAGTACCCACC

GTCTCCCGCCGCACAGTCTCTAATG TTACACTCGGACCTCGTCGACATGTTCCCAGACGACTTAATAAGTGCCCCACCCCCTCCTAAGTTTCGAGCTGGACTCAGTGCAGAGACTAGCCTCCAGTACCCACC

GTCTCCCGCCGCACAGTCTCTAATG TTACACTCGGACCTCGTCGACATGATTTCTTCTGCATGGGGCTTAGCTCCACCCCCTCCTAAGTTTCGAGCTGGACTCAGTGCAGAGACTAGCCTCCAGTACCCACC

GTCTCCCGCCGCACAGTCTCTAATG TTACACTCGGACCTCGTCGACATGACGATGAGCTCTGTCTTGGCTGGCATGTTCCCTCCTAAGTTTCGAGCTGGACTCAGTGCAGAGACTAGCCTCCAGTACCCACC

GTCTCCCGCCGCACAGTCTCTAATG TTACACTCGGACCTCGTCGACATGTATTCTGCAATGGGGTTCCTGTCAGCAGCACCTCCTAAGTTTCGAGCTGGACTCAGTGCAGAGACTAGCCTCCAGTACCCACC

GTCTCCCGCCGCACAGTCTCTAATG TTACACTCGGACCTCGTCGACATGTTGGACTGGTTGTTCTTGAGTCGATGGGCCCCTCCTAAGTTTCGAGCTGGACTCAGTGCAGAGACTAGCCTCCAGTACCCACC

GTCTCCCGCCGCACAGTCTCTAATG TTACACTCGGACCTCGTCGACATGTTCATTCTCACGAATGATGGCCCGGCTGTCCCTCCTAAGTTTCGAGCTGGACTCAGTGCAGAGACTAGCCTCCAGTACCCACC

GTCTCCCGCCGCACAGTCTCTAATG TTACACTCGGACCTCGTCGACATGCACCAGGAAGGGACCTTTTGTGTGGCCCTCCTCCTAAGTTTCGAGCTGGACTCAGTGCAGAGACTAGCCTCCAGTACCCACC

GTCTCCCGCCGCACAGTCTCTAATG TTACACTCGGACCTCGTCGACATGTCATCAGCTAGAATGGTGTCAGTGCCCTGGCCTCCTAAGTTTCGAGCTGGACTCAGTGCAGAGACTAGCCTCCAGTACCCACC

GTCTCCCGCCGCACAGTCTCTAATG TTACACTCGGACCTCGTCGACATGTCTTCTTCTTATACTTGCGGGGCTGGGCAGCCTCCTAAGTTTCGAGCTGGACTCAGTGCAGAGACTAGCCTCCAGTACCCACC

GTCTCCCGCCGCACAGTCTCTAATG TTACACTCGGACCTCGTCGACATGTACGCTTGTCGCTCTTCCCATCATCCTCGCCCTCCTAAGTTTCGAGCTGGACTCAGTGCAGAGACTAGCCTCCAGTACCCACC

GTCTCCCGCCGCACAGTCTCTAATG TTACACTCGGACCTCGTCGACATGTGGAAGATTTCCAGCTGAAGCTCCTTGGCCCCTCCTAAGTTTCGAGCTGGACTCAGTGCAGAGACTAGCCTCCAGTACCCACC

GTCTCCCGCCGCACAGTCTCTAATG TTACACTCGGACCTCGTCGACATGACTCTCGCTCTGATCTGCCTTGAAGAGGACCCTCCTAAGTTTCGAGCTGGACTCAGTGCAGAGACTAGCCTCCAGTACCCACC

GTCTCCCGCCGCACAGTCTCTAATG TTACACTCGGACCTCGTCGACATGGGGGGTGGGACATCTGGATTTCCATCTGCCCTCCTAAGTTTCGAGCTGGACTCAGTGCAGAGACTAGCCTCCAGTACCCACC

GTCTCCCGCCGCACAGTCTCTAATG TTACACTCGGACCTCGTCGACATGCGGCAGTACTCCATGTGATCATCCTCCTCCCCTCCTAAGTTTCGAGCTGGACTCAGTGCAGAGACTAGCCTCCAGTACCCACC

GTCTCCCGCCGCACAGTCTCTAATG TTACACTCGGACCTCGTCGACATGTTCTTCTTCTTCCTCCTTGGCCTCCCACTGCCTCCTAAGTTTCGAGCTGGACTCAGTGCAGAGACTAGCCTCCAGTACCCACC

GTCTCCCGCCGCACAGTCTCTAATG TTACACTCGGACCTCGTCGACATGAGAATAATTTCCCCACCCTGCTGGCACACCCCTCCTAAGTTTCGAGCTGGACTCAGTGCAGAGACTAGCCTCCAGTACCCACC

GTCTCCCGCCGCACAGTCTCTAATG TTACACTCGGACCTCGTCGACATGACAGTAATCCTGGTGATCCGTCTCGTAGCCCCTCCTAAGTTTCGAGCTGGACTCAGTGCAGAGACTAGCCTCCAGTACCCACC

GTCTCCCGCCGCACAGTCTCTAATG TTACACTCGGACCTCGTCGACATGCTGCCACTGTCCAGGTCTGATTCCTCAGCCCTCCTAAGTTTCGAGCTGGACTCAGTGCAGAGACTAGCCTCCAGTACCCACC

GTCTCCCGCCGCACAGTCTCTAATG TTACACTCGGACCTCGTCGACATGATACGAGCCTCCTTTCTTCCTCTTGCCACCCCTCCTAAGTTTCGAGCTGGACTCAGTGCAGAGACTAGCCTCCAGTACCCACC

GTCTCCCGCCGCACAGTCTCTAATG TTACACTCGGACCTCGTCGACATGTGTTGGCACTGAACTCTCTCCATTTGGCCCCCTCCTAAGTTTCGAGCTGGACTCAGTGCAGAGACTAGCCTCCAGTACCCACC

GTCTCCCGCCGCACAGTCTCTAATG TTACACTCGGACCTCGTCGACATGAGTTGGTGAGCGTGTGGTAATCCTCCTCAGCCTCCTAAGTTTCGAGCTGGACTCAGTGCAGAGACTAGCCTCCAGTACCCACC

GTCTCCCGCCGCACAGTCTCTAATG TTACACTCGGACCTCGTCGACATGTCTCGGTGCTTCCTTCTTCGTTTCCGACCCCCTCCTAAGTTTCGAGCTGGACTCAGTGCAGAGACTAGCCTCCAGTACCCACC

GTCTCCCGCCGCACAGTCTCTAATG TTACACTCGGACCTCGTCGACATGTCACTGTCACGCTTCTTGCGTTTTCGGGGTCCTCCTAAGTTTCGAGCTGGACTCAGTGCAGAGACTAGCCTCCAGTACCCACC

GTCTCCCGCCGCACAGTCTCTAATG TTACACTCGGACCTCGTCGACATGTCCTGGCTTGTTCTCCTTCTGCTTCTTGGGCCTCCTAAGTTTCGAGCTGGACTCAGTGCAGAGACTAGCCTCCAGTACCCACC

GTCTCCCGCCGCACAGTCTCTAATG TTACACTCGGACCTCGTCGACATGTCTCTTCTTCACACCCAATGCTGACGGCAGCCTCCTAAGTTTCGAGCTGGACTCAGTGCAGAGACTAGCCTCCAGTACCCACC

GTCTCCCGCCGCACAGTCTCTAATG TTACACTCGGACCTCGTCGACATGACTTCTTGCCCACAGGATCACAGTGTCTGCCCTCCTAAGTTTCGAGCTGGACTCAGTGCAGAGACTAGCCTCCAGTACCCACC

GTCTCCCGCCGCACAGTCTCTAATG TTACACTCGGACCTCGTCGACATGCAGTCGTCTATACAGATCACCTCCCCGGCCCTCCTAAGTTTCGAGCTGGACTCAGTGCAGAGACTAGCCTCCAGTACCCACC

GTCTCCCGCCGCACAGTCTCTAATG TTACACTCGGACCTCGTCGACATGCATAGCCATGGAGGACAATCCCAGCCAGAACCTCCTAAGTTTCGAGCTGGACTCAGTGCAGAGACTAGCCTCCAGTACCCACC

GTCTCCCGCCGCACAGTCTCTAATG TTACACTCGGACCTCGTCGACATGTGTGCCAGATCTCATTGAGTTTCCCCGAGGCCTCCTAAGTTTCGAGCTGGACTCAGTGCAGAGACTAGCCTCCAGTACCCACC

GTCTCCCGCCGCACAGTCTCTAATG TTACACTCGGACCTCGTCGACATGATAGCTGCCCGTTCCTCATTCTGCCACAGTCCTCCTAAGTTTCGAGCTGGACTCAGTGCAGAGACTAGCCTCCAGTACCCACC

GTCTCCCGCCGCACAGTCTCTAATG TTACACTCGGACCTCGTCGACATGTTCTCTCTTGCCCAAATCCCCTGTTTCCCCCCTCCTAAGTTTCGAGCTGGACTCAGTGCAGAGACTAGCCTCCAGTACCCACC

GTCTCCCGCCGCACAGTCTCTAATG TTACACTCGGACCTCGTCGACATGTCCTCCTGGTTTTCAGCTTCCTCCTTCTCACCTCCTAAGTTTCGAGCTGGACTCAGTGCAGAGACTAGCCTCCAGTACCCACC

GTCTCCCGCCGCACAGTCTCTAATG TTACACTCGGACCTCGTCGACATGAGGCCTTAAACTCCTTCTCAGTCTTGCCCCCCTCCTAAGTTTCGAGCTGGACTCAGTGCAGAGACTAGCCTCCAGTACCCACC

GTCTCCCGCCGCACAGTCTCTAATG TTACACTCGGACCTCGTCGACATGGTTGAAGCCCAGCACCTCAATGTTGCCCCCCTCCTAAGTTTCGAGCTGGACTCAGTGCAGAGACTAGCCTCCAGTACCCACC

GTCTCCCGCCGCACAGTCTCTAATG TTACACTCGGACCTCGTCGACATGAAGATGATGACAGTGTCAGCAGTGGCCAGACCTCCTAAGTTTCGAGCTGGACTCAGTGCAGAGACTAGCCTCCAGTACCCACC

GTCTCCCGCCGCACAGTCTCTAATG TTACACTCGGACCTCGTCGACATGGGATATGTCAGCAAACTCCTCCAGGAAGCCCCTCCTAAGTTTCGAGCTGGACTCAGTGCAGAGACTAGCCTCCAGTACCCACC

GTCTCCCGCCGCACAGTCTCTAATG TTACACTCGGACCTCGTCGACATGATCTACCACAAGACAGGCCCAGCGGATGGCCTCCTAAGTTTCGAGCTGGACTCAGTGCAGAGACTAGCCTCCAGTACCCACC

GTCTCCCGCCGCACAGTCTCTAATG TTACACTCGGACCTCGTCGACATGGCTTTCTTGCCCCCTTTGATGGCATTGTCCCCTCCTAAGTTTCGAGCTGGACTCAGTGCAGAGACTAGCCTCCAGTACCCACC

GTCTCCCGCCGCACAGTCTCTAATG TTACACTCGGACCTCGTCGACATGACGATGGTTTGTATGGTCTTGCCTAGCCCCCCTCCTAAGTTTCGAGCTGGACTCAGTGCAGAGACTAGCCTCCAGTACCCACC

GTCTCCCGCCGCACAGTCTCTAATG TTACACTCGGACCTCGTCGACATGATCTTCCTCCCACGTGGACTGGTCATATGGCCTCCTAAGTTTCGAGCTGGACTCAGTGCAGAGACTAGCCTCCAGTACCCACC

GTCTCCCGCCGCACAGTCTCTAATG TTACACTCGGACCTCGTCGACATGATCCACACTGTGGTTGATGATGCGGTGGACCCTCCTAAGTTTCGAGCTGGACTCAGTGCAGAGACTAGCCTCCAGTACCCACC

GTCTCCCGCCGCACAGTCTCTAATG TTACACTCGGACCTCGTCGACATGTACTTCTCCTCCATCTCAGCATAGTGCGGGCCTCCTAAGTTTCGAGCTGGACTCAGTGCAGAGACTAGCCTCCAGTACCCACC

GTCTCCCGCCGCACAGTCTCTAATG TTACACTCGGACCTCGTCGACATGCAGCCCAGGACCTTCTTCTTCTTCCTTCCTCCTCCTAAGTTTCGAGCTGGACTCAGTGCAGAGACTAGCCTCCAGTACCCACC

GTCTCCCGCCGCACAGTCTCTAATG TTACACTCGGACCTCGTCGACATGCCGGCCTCTCTTTAGTTTCTTGGTGCGGACCCTCCTAAGTTTCGAGCTGGACTCAGTGCAGAGACTAGCCTCCAGTACCCACC

GTCTCCCGCCGCACAGTCTCTAATG TTACACTCGGACCTCGTCGACATGGCCCTCTTTGGTTTTGGCTCTTCGGATGGGCCTCCTAAGTTTCGAGCTGGACTCAGTGCAGAGACTAGCCTCCAGTACCCACC

GTCTCCCGCCGCACAGTCTCTAATG TTACACTCGGACCTCGTCGACATGTGACTTCTGTTCCACTTGCTTTTGCCCCCCCCTCCTAAGTTTCGAGCTGGACTCAGTGCAGAGACTAGCCTCCAGTACCCACC

GTCTCCCGCCGCACAGTCTCTAATG TTACACTCGGACCTCGTCGACATGTCCGGTTCCATATTCACTGCCCCCACTCTCCTCCTAAGTTTCGAGCTGGACTCAGTGCAGAGACTAGCCTCCAGTACCCACC

GTCTCCCGCCGCACAGTCTCTAATG TTACACTCGGACCTCGTCGACATGGACTTCTCCCGGTACTCATCTCGCTCAGAACCTCCTAAGTTTCGAGCTGGACTCAGTGCAGAGACTAGCCTCCAGTACCCACC

GTCTCCCGCCGCACAGTCTCTAATG TTACACTCGGACCTCGTCGACATGTGTCATCCTTATCAGGCATCCTGTCACCCCCCTCCTAAGTTTCGAGCTGGACTCAGTGCAGAGACTAGCCTCCAGTACCCACC

GTCTCCCGCCGCACAGTCTCTAATG TTACACTCGGACCTCGTCGACATGACCAATGCCCCCACCATTTCCTTCTCCTTCCCTCCTAAGTTTCGAGCTGGACTCAGTGCAGAGACTAGCCTCCAGTACCCACC

GTCTCCCGCCGCACAGTCTCTAATG TTACACTCGGACCTCGTCGACATGTCTCCCCAATTCTGCTGTTCTTCTCTGGCTCCTCCTAAGTTTCGAGCTGGACTCAGTGCAGAGACTAGCCTCCAGTACCCACC

GTCTCCCGCCGCACAGTCTCTAATG TTACACTCGGACCTCGTCGACATGCCCCCACCGCCAATGTAGGATCTTCTGCACCTCCTAAGTTTCGAGCTGGACTCAGTGCAGAGACTAGCCTCCAGTACCCACC

GTCTCCCGCCGCACAGTCTCTAATG TTACACTCGGACCTCGTCGACATGTTCTCCCCTTCTTCCTCTCCCTCCTCTTCCCTCCTAAGTTTCGAGCTGGACTCAGTGCAGAGACTAGCCTCCAGTACCCACC

GTCTCCCGCCGCACAGTCTCTAATG TTACACTCGGACCTCGTCGACATGCTTTCCCCGAAGCTTCTTGCGTCCATCAGGCCTCCTAAGTTTCGAGCTGGACTCAGTGCAGAGACTAGCCTCCAGTACCCACC
