## Supplementary legends for "Exon Junction Complex dependent mRNA localization is linked to centrosome organization during ciliogenesis"

### Supplementary figure legends

**Figure S1. Fluorescence intensity measurement of EJC proteins around centrosomes, increase of eIF4A3 and Y14 upon differentiation and their colocalization with nuclear speckle markers.** Quiescent mNSC (a, c) and multiciliated ependymal (b, d) cells were stained for eIF4A3 (a, b) or Y14 (c, d). Centrosomes were labeled by FOP antibody and primary cilia and centrioles were stained by poly-glutamylated tubulin antibody. Nuclei were stained by Hoechst. Images result from maximum intensity projections of 12 z-stacks acquired at every 0.5 μm. Fluorescence intensity of eIF4A3 (a) and Y14 (c) were determined in a 2 μm circle around centrosomes and base of cilia. In multiciliated ependymal cells, 2 μm circles were manually selected at the base of multicilia, and fluorescence intensities of eIF4A3 (b) and Y14 (d) were determined. Scale bars are 3 μm. Relative fluorescence intensities of eIF4A3 (Figure1a, b) and Y14 (Figure 1c, d) in the nucleus were determined in Hoechst stained area and plotted in panel e and f as described in legend of Figure 1. RPE1 cells were stained with eIF4A3 and SC35 (g) and Y14 and 9G8 (h) antibodies. Scale bars represent 10 μm (g, h). Images from white dashed squares from g and h are shown in i and j, respectively. Relative fluorescence intensity profile of eIF4A3 with SC35 (i) and Y14 with 9G8 (j) along the red line on nuclear speckle were plotted. Average fluorescence intensity of each protein on the line is set to 1.0.

**Figure S2. Y14, core component of EJC accumulates around centrosome during quiescence, while MLN51 broadly exist in cytosol.**

Proliferating (a, e) and quiescent (b, f) RPE1 cells were stained for Y14 (a, b) or MLN51 (e, f). Primary cilia and centrioles were stained by poly-glutamylated tubulin antibody (a, b, e, f). Centrosomes were labeled by FOP antibody (e, f). Nuclei were stained by Hoechst. Right panels show enlarged images of the white dashed square in left. Scale bars in left and right panels are 10 μm and 3 μm, respectively (a, b, e, f). The fraction of cells with detectable Y14 (c) was determined in either proliferating or quiescent RPE1 cells. Error bars correspond to S.D. Quantifications of Y14 fluorescence intensities were performed as described in the legend of figure 1 except that average fluorescence intensity of Y14 in proliferating cells is set to 1.0 (d). ** $P \leq 0.01$, and **** $P \leq 0.0001$, two-tailed t-test (c) and Mann-Whitney test (d). Three independent experiments were performed.

**Figure S3. Y14 around centrosome, RPE1 cell cycle and general translation activity upon cell cycle re-entry.**

Quiescent RPE1 cells were incubated with 10 % serum containing media during the indicated times, and centrosomal Y14, cell cycle and translation efficiency were analyzed by immunofluorescence (a, b), flow cytometry (c, d) and SUnSET (e), respectively. Primary cilia and centriole were stained by poly-glutamylated tubulin antibody. Nuclei were stained by Hoechst. Right panels show enlarged images of the white dashed square in the left panel. Scale bars in the left panels are 10 μm, and scale bars in right panels are 3 μm (a). Quantification of Y14 fluorescence intensities was performed as described in the legend of figure 1. Average fluorescence intensity of Y14 in cells with 0 hr incubation is set to 1.0. n.s $P > 0.05$ and **** $P \leq 0.0001$, Mann-Whitney test (b). Representative cell cycle profiles at each incubation times are depicted (c). Cell cycle was determined by Hoechst staining. The proportion of cells in each phase was analyzed by FlowJo from four independent experiments. Error bars correspond to S.D (d). Nascent peptides were visualized by western blot with puromycin antibody (e), and relative translation activity was normalized by total protein staining (e, f). Translation efficiency at 0 hr is set to 1.0. Three independent experiments were performed (a, b, e, f).

**Figure S4. Stress granules and general translation independent EJC accumulation around centrosome.**

Quiescent RPE1 cells stably expressing centrin1-GFP were stained for stress granule marker protein TIA1 and either eIF4A3 or Y14 (a). Lower panels show enlarged images marked by white dashed square in the upper panel (a). RPE1 cells were treated with sodium arsenite or not (b) prior to fixation and stained with eIF4E and TIA-1 (b). Quiescent RPE1 cell (c, e) or proliferating RPE1 cells (g, i) were treated with DMSO, Puromycin, or Cycloheximide prior to fixation and stained with either eIF4A3 (c, g) or Y14 (e, i) antibodies. Centrioles and cilia were stained by poly-glutamylated tubulin antibody (c, e, i) and centrosomes were labeled with FOP antibody (c, g). Nuclei were stained by Hoechst. Scale bars in the upper and lower panels are 10 μm and 3 μm, respectively (a). Scale bars in b and scale bars in c, e, g, i are 10 μm and 3 μm, respectively. Quantifications of fluorescence intensities of eIF4A3 (d, h) and Y14 (f, j) were performed as described in the legend of figure 1. The average fluorescence intensities of eIF4A3 or Y14 in DMSO treated cells are set to 1.0. The numbers of cells analyzed from three independent experiments are depicted (d, f, h, j). n.s P > 0.05, Mann-Whitney test.

**Figure S5. RNA-dependent localization of assembled EJC around centrosome.**
Y14 antibody stained quiescent RPE1 cells treated with indicated siRNAs (a). Permeabilized proliferating RPE1 cells (d, e) and quiescent (g, h) were incubated with RNAse A or not prior to fixation and stained for EDC4 and DDX6 (d, e) or Y14 (g, h). Primary cilia and centriole were stained by poly-glutamylated tubulin antibody (a, g, h). Nuclei were stained by Hoechst. Upper (a) or right (g, h) panels show enlarged images marked by white dashed square in upper (a) or left panels (g, h). Scale bars in upper (a) or left (g, h) and lower (a) or right panels (g, h) are 10 μm and 3 μm, respectively. Scale bars in d and e are 10 μm. Yellow arrows represent EDC4 and DDX6 double positive bodies (d). Quantifications of fluorescence intensities for Y14 in the nucleus (b) were determined in Hoechst stained area. Relative fluorescence intensity of Y14 around centrosomes (c, i) or in speckles (j, k) were determined in 2 μm circle around centrosome (c, i) or 1 μm on speckles (j, k), respectively and plotted as described in legend of figure 1. One 1 μm circle per cell were manually selected in nuclei. The average fluorescence intensity of Y14 in Ctrl siRNA treated cells (b, c) or Buffer treated condition (i-k) is set to 1.0. Number of EDC4 and DDX6 double positive bodies/cell was determined in either buffer or RNaseA treated cells (f). Error bar means S.D (f). n.s P > 0.05 and **** P ≤ 0.0001, Mann-Whitney test (b, c, i, j, k) and two-tailed t-test (f). Three independent experiments were performed**.**

**Figure S6. Dynein dependent transport along microtubule is required for Y14 accumulation around centrosomes.**

Proliferating RPE1 cells were treated with either DMSO or Nocodaole (a, b) or either

DMSO or ciliobrevinD (c) and stained with eIF4A3 and β-tubulin (a, b) or GM130 (c) antibodies. Scale bars are 10 μm (a-c). Y14 antibody stained quiescent cells treated with either DMSO or Nocodazole (d), either DMSO or CiliobrevinD (e), or chilled quiescent cells subjected to a microtubule regrowth assay (f). Primary cilia and centriole were stained by poly-glutamylated tubulin antibody (d, e) and microtubules were visualized by α-tubulin antibody. Nuclei were stained by Hoechst. Lower panels show enlarged images marked by white dashed square in the upper panel (d-f). Scale bars in upper and lower panels represent 10 μm and 3 μm, respectively (d-f). Quantification of fluorescence intensities of Y14 (g-i) was performed as described in the legend of figure 1. The average fluorescence intensities for Y14 in DMSO treated cells (g, h) or in pre-incubated quiescent cells (i) are set to 1.0. n.s $P > 0.05$, ** $P \leq 0.01$, *** $P \leq 0.001$, and **** $P \leq 0.0001$, Mann-Whitney test. Three independent experiments were performed.

**Figure S7. A high-throughput smiFISH screen identifies mRNAs localizing to the base of cilia and translating ribosome is required for localization of *NIN* and *BICD2* mRNAs.**

In a high-throughput smiFISH screen, quiescent RPE1 cells stably expressing Centrin1-GFP were stained for each indicated RNA and Arl13b. Nuclei were labeled with DAPI. Images were processed by maximum intensity projections of 35 z-stacks acquired at every 0.35 μm. Scale bar represents 10 μm (a). Quiescent RPE1 cells stably expressing centrin1-GFP were stained by probes against *BICD2* mRNA (b) or *NIN* mRNA (c) after 30 min incubation with either DMSO, cyloheximide, or puromycin. Right panels display enlarged images of the white dashed square in the left panel. Scale bars in the left panels are 10 μm, and scale bars in right panels are 3 μm. Images result from maximum intensity projections of 14 z-stacks acquired at every 0.5 μm (b, c). Proportion of cells displaying centrosomal *BICD2* (d) or *NIN* (e) RNA pattern is depicted. Error bars correspond to S.D. Three independent experiments were performed. n.s $P > 0.05$ and **** $P \leq 0.0001$, two-tailed t-test.

**Figure S8. Both *NIN* and *BICD2* mRNAs are associated with EJC during quiescence and knock-downs of eIF4A3 and Y14 alter pericentriolar material organization.**

The association of EJC with centrosomal (*NIN* and *BICD2*) and intronless (*SFM3B5* and *SDHAF1*) mRNAs in quiescent RPE1 cells was determined by RNA immunoprecipitation (RIP)-qPCR analysis of eIF4A3, Y14, and Rab5. Enriched mRNA levels in pull down fraction compare to input from two independent experiments are depicted as % input (a). Quiescent RPE1 cells treated with indicated siRNAs were analyzed for *NIN* mRNA levels by RT-qPCR. Average *NIN* mRNA levels normalized by *GAPDH* mRNA levels from three independent experiments are depicted. Ctrl siRNA treated condition is set to 1.0 (b). Error bar means S.D (a, b). Quiescent RPE1 cells transfected with Ctrl, eIF4A3 or Y14 siRNA (c, f) were stained for pericentrin and γ-tubulin (c) or for β-tubulin and PCM1 (f). Images were resulted from maximum intensity projections of 6 z-stacks acquired at every 0.5 μm (c). Lower panels are enlarged images marked by white dashed square in the upper panels (c, f). Scale bars in upper panels of c and f are 10 μm and scale bars in the lower panels of c and f are 3 μm and 5 μm respectively. Quantifications of fluorescence intensities for pericentrin (d) and γ-tubulin (e) were performed as described in the legend of figure 1 except that the average fluorescence intensities of pericentrin and γ-tubulin in Ctrl siRNA treated cells are set to 1.0. Images corresponding to 120 cells from three independent experiments were analyzed. n.s $P > 0.05$, **** $P \leq 0.0001$. Two-tailed t-test (a) and Mann-Whitney test (d, e).

**Figure S9. Depletion of MAGOH but not of MLN51 affects NIN protein level at centrosomes and centrosome organization.**

Knock down efficiency of siRNAs was determined by either Western blotting (a) or qPCR (b, c). Relative protein (a) or RNA level of MAGOH (b) and MLN51 (c) normalized by GAPDH is depicted. Ctrl siRNA treated condition is set to 1.0 (a-c). Quiescent RPE1 cells transfected with indicated siRNAs were stained for NIN (d) or PCM1, FOP, and poly-glutamylated tubulin (f). Nuclei were stained by Hoechst. Lower panels are enlarged images marked by white dashed square in the upper panels. Scale bars in the upper and lower panels are 10 μm and 3 μm, respectively (d, f). Images were processed by maximum intensity projections of 15 z-stacks acquired at every 0.5 μm (d). Quantification of fluorescence intensities of NIN were performed as described in the legend of figure 1. The average fluorescence intensity of NIN in Ctrl siRNA treated cells is set to 1.0 (e).

Proportion of cells with ectopic centriolar satellite with FOP upon the siRNA treatments is represented (g). Proportion of ciliated cells upon the indicated siRNA treatments (h). Error bars correspond to S.D (b, c, g, h). n.s $P > 0.05$, * $P \leq 0.05$, ** $P \leq 0.01$, and **** $P \leq 0.0001$, Mann-Whitney test (e) and Two tailed t-test (b, c, g, h). Three independent experiments were performed.

**Table S1. High-throughput smFISH screen identifies transcripts displaying specific localization in quiescent RPE1 cell.**

Sequence reference number and gene name of analyzed mRNAs are depicted with subcellular localization patterns.

**Table S2. Sequence of smFISH probes in high-throughput smFISH screen**

Sequences of smFISH probes for each transcript are depicted.
